# Capturing the Hierarchically Assorted Modules of Protein Interaction in the Organized Nucleome

**DOI:** 10.1101/2022.08.14.503837

**Authors:** Shuaijian Dai, Shichang Liu, Chen Zhou, Fengchao Yu, Guang Zhu, Wenhao Zhang, Haiteng Deng, Al Burlingame, Weichuan Yu, Tingliang Wang, Ning Li

**Affiliations:** Division of Life Science, The Hong Kong University of Science and Technology, Hong Kong SAR, China; Department of Chemical and Biological Engineering, The Hong Kong University of Science and Technology, Hong Kong, China; Tsinghua-Peking Joint Centre for Life Sciences, Centre for Structural Biology, School of Life Sciences and School of Medicine, Tsinghua University, Beijing 100084, China; The HKUST Shenzhen-Hong Kong Collaborative Innovation Research Institute, Futian, Shenzhen, Guangdong, 518057, China; Department of Electronic and Computer Engineering, The Hong Kong University of Science and Technology, Hong Kong SAR, China; Department of Pharmaceutical Chemistry, University of California, San Francisco, San Francisco, United States

**Keywords:** Nucleome, Protein-protein interaction, qXL-MS, Modulome, Community, CHAMPION

## Abstract

Nuclear proteins are major constituents and key regulators of the topological organization of nucleome. To elucidate the global connectivity of nucleomic proteins and to decipher the hierarchically organized modules of protein interaction that are involved in nucleomic organization and nuclear events, both formaldehyde and CBDPS crosslinkers were applied sequentially on the *in vivo* prefixed nuclei to perform a double <u>c</u>hemical <u>crossl</u>inking with <u>m</u>ass <u>s</u>pectrometry (XL-MS) analysis. The integration of dimethyl-labelling with XL-MS generated a quantitative XL-MS workflow (qXL-MS) that consequently identified 5,340 cross-linked peptides (crosslinks) from nucleome. These crosslinks were construed into 1,297 nuclear <u>p</u>rotein-<u>p</u>rotein interactions (PPIs), from which discovered were 250 and 26 novel interactors of histones and nucleolar box C/D snoRNP complex, respectively. MONET-based modulomic analysis of their Arabidopsis orthoglous PPIs constructed 27 and 24 master <u>n</u>uclear <u>p</u>rotein interaction <u>m</u>odules (NPIMs) that contain the condensate-forming protein(s) and the intrinsically <u>d</u>isordered region (IDR)-containing proteins, respectively. These NPIMs successfully captured the previously reported nuclear protein complexes and nuclear bodies in nucleome. Surprisingly, modulomic analysis showed that these NPIMs were hierarchically assorted into four communities of NPIMs in nucleome including Genome Community and Nucleolus Community. The qXL-MS-based quantitative interactomics finally revealed 17 Hormone-specific module variants participating in a broad range of nuclear events. Thus, this integrated pipeline of qXL-MS and MONET modulomics, named as CHAMPION, is capable of capturing both nuclear protein complexes and nuclear bodies, constructing the topological architecture of protein interaction modules and module variants in nucleome and probably of mapping the protein compositions of condensates.

**Highlights:**

1. The formaldehyde and CBDPS crosslinkers coupled qXL-MS discovered 5,340 crosslinked peptides. These crosslinks were construed into 1,297 nuclear <u>p</u>rotein-<u>p</u>rotein interactions (PPIs), protein components of which contained 250 and 26 novel interactors of histone octamer and nucleolar box C/D snoRNP complex, respectively, in the intricately organized nucleome.
2. The MONET-based modulomic analysis of these crosslinks captured 95 <u>n</u>uclear <u>p</u>rotein interaction <u>m</u>odules (NPIMs), a portion of which contain both the condensate-forming and the intrinsically <u>d</u>isordered region (IDR)-containing proteins. Especially, some NPIMs captured 6 previously reported nuclear protein complexes.
3. A number of Hormone-specific module variants were identified by modulomics upon hormone treatment using the hormone significantly up-regulated crosslinks from qXL-MS. Several PPIs and NPIMs have been substantiated with alternative biological experiments.
4. This CHAMPION pipeline has partitioned these NPIMs into four hierarchically and topologically organized communities in nucleome. The molecular functions of those proteins partitioned into C1 and C2 community are specialized in genome organization and nucleolar functions, respectively.

## Introduction

Nucleus is the central organelle of eukaryotic cells that stores the hereditary information and harbors thousands of nuclear proteins interacting with DNA and RNA molecules to coalesce into the intricately organized genome, consisting of chromatin fibers, topologically <u>a</u>ssociated <u>d</u>omains (TADs) and chromatin loops. These nuclear proteins participate in the formation of a wide spectrum of membrane-less biomolecular condensates ranging from speckles, paraspeckles, nuclear bodies to nucleoli (Strom and Brangwynne, 2019). These chromatin fibers, liquid and solid nuclear bodies are compartmentalized either within the chromosomal territories, subnuclear compartments or the interchromatin spaces (Fraser and Bickmore, 2007), comprising the entire content inside nucleus, or called nucleome (Dekker et al., 2017), which is surrounded by a nuclear envelope punctated with nuclear pores. A highly diverse array of cytosolically synthesized nuclear proteins pass through the pores spatiotemporally into nucleome to regulate the self-organization of genome, chromosomal refolding, DNA replication and repair, gene expression, RNA processing, ribosome biogenesis and other nuclear events (Meier et al., 2017; Rao et al., 2014a; Tang et al., 2020).

Among these nuclear events, those nuclear protein groups may play an arguably dominant role (Misteli, 2020). Generally speaking, the chromatin-binding proteins largely participate in regulation of the chromatin polymer-polymer interactions, in enhancement of the formation of chromatin loops and chromosomal domains as well as in defining of the overall conformation of chromatin fibers (Szabo et al., 2019). For examples, the chromosomal loop structure is often mediated by the CCCTC-binding factor (CTCF) and cohesin complexes (Hnisz et al., 2016), whereas the heterochromatin protein 1 HP1α promotes the compartmentalization of constitutive heterochromatin by organizing repetitive sequences into compacted assemblies around nucleolus (Larson et al., 2017; Strom and Brangwynne, 2019). Even the dynamic local motion and transient interaction of chromatins are in fact mediated by chromatin-forming proteins, which associate and dissociate with chromatins in rapid cycles (Szabo et al., 2019).

Histone proteins, as one group of well-known nuclear proteins incorporated into nucleosomes as subunits, are complexed with DNA molecules to form the basic chromosomal fibers of genome (Bannister and Kouzarides, 2011; Taverna et al., 2007; Zhang and Reinberg, 2001). The multiple post-translational modifications (PTMs) occurring at the tail regions of these histone proteins encrypt the essential epigenetic information, or called “histone code” (Strahl and Allis, 2000). The histone modifying enzymes are another group of well-known nuclear proteins, called writers and erasers, responsible for encrypting and erasing the histone code, respectively. The epigenetic information encoded by histone PTMs can be translated into a variety of downstream molecular events by proteins of specific functions, called readers, ranging from dynamic changes of chromosome conformation, replication to gene expression (Musselman et al., 2012; Spitz and Furlong, 2012; Taverna et al., 2007). To profile histone PTMs and identify histone-interacting proteins, such as transcription factors (TFs), both proteomics and chromatin immunoprecipitation followed by sequencing (ChIP-seq) have been applied to enable the unbiased and comprehensive analysis of DNA-interacting and DNA association proteins (Baker, 2012; Ding et al., 2013; Lu et al., 2021; Park, 2009). However, little is known about the overarching profiles and interaction networks of chromosome-interacting proteins, genome organization proteins, histone-modifying enzymes and gene expression regulators from a nucleome-wide perspective.

Other than these chromosome-modifying and chromatin architecture-organizing proteins (Hnisz et al., 2017; Yamamoto and Schiessel, 2016), the third group of nuclear proteins, the homotypic proteins, are the major drivers of the sub-organellar compartment, or called biomolecular condensate, formation processes. These nuclear proteins undergo liquid-liquid <u>p</u>hase <u>s</u>eparation (LLPS) *via* protein-protein interaction, protein-RNA and protein-DNA interactions, mediating the formation and stabilization of the chromatin interactions, TADs, the separation of euchromatin and heterochromatin as well as regulation of gene expression (Misteli, 2020; Zidovska, 2020). These nuclear condensate-forming proteins, like histones, promote the assembly of a wide array of membrane-less organelles (Banani et al., 2017). They include Cajal bodies that play a role in mRNA metabolism and <u>s</u>mall <u>n</u>uclear ribo<u>n</u>ucleo<u>p</u>rotein (snRNP) biogenesis (Riback et al., 2020), nuclear speckles that are rich in RNA metabolism-related factors and function to concentrate a wide range of gene regulation and RNA metabolism factors (Galganski et al., 2017), DNA damage foci where DNA damage response proteins concentrate, dicing bodies that are involved in <u>mi</u>croRNA (miRNA) processing, photobodies that mediate light signalling (Emenecker *et al*., 2020), and finally nuclear pore complexes that are responsible for molecular transports (Shin and Brangwynne, 2017). Some of these multivalent nuclear protein interactions are mediated either by intrinsically-<u>d</u>isordered regions (IDR) or the low-<u>c</u>omplexity <u>s</u>equences (LCS) (Murray et al., 2017; Nott et al., 2015). To comprehend the protein compositions of condensates, several proteomic experiments have been performed on Cajal bodies, speckles, dicing bodies and photobodies (Fang and Spector, 2007; Quail, 2021; Riback et al., 2020; Saitoh et al., 2004; Song et al., 2007). For an example, the proteomics analysis of Cajal bodies indicated the protein Coilin being the scaffold protein in the condensate-formation of Cajal body while Argonaute-4 (AGO4) and additional <u>s</u>mall interfering RNA (siRNA) complex components being conjugated as client molecules (Li et al., 2006). Similarly, in the case of proteomic study of nuclear speckles, it was found that the <u>s</u>erine/arginine-rich (SR) proteins are abundant components, playing essential roles in splicing regulation (Saitoh et al., 2004). These results have again clearly demonstrated the usefulness of proteomics in identification of protein components in diverse nuclear bodies or condensates.

Nucleolus, as the predominant membrane-less organelle in nucleus, is considered to be a conspicuous biomolecular condensate. It is a key nucleomic component where both ribosome RNA biogenesis and ribosome assembly occur (Pederson, 2011), and it is involved in many important biological processes (Emenecker et al., 2020; Lafontaine et al., 2021). The forth major groups of nuclear proteins are the nucleolar condensate-forming proteins that play an important role in the compartmentalization of nucleolus into three regions of condensates: fibrillar <u>c</u>enter (FC), <u>d</u>ense fibrillar <u>c</u>omponent (DFC) and <u>g</u>ranular <u>c</u>omponent (GC) *via* LLPS (Emenecker et al., 2020; Feric et al., 2016). Ribosome assembly factors, such as the box C/D small nucleolar RNA-associated methyltransferase <u>fib</u>rillarin (FIB) and other proteins, are well-known to participate in the assembly of DFC around FC (Xing et al., 2017; Yao et al., 2019). The R/G domains on FIB and rRNA transcription factor nucleolin are both necessary and sufficient to induce the LLPS during nucleolus formation (Emenecker et al., 2020; Feric et al., 2016). To elucidate the overall protein composition of nucleolar condensate, proteomic studies have been performed before and revealed 1400 and 1600 proteins associated with human and Arabidopsis nucleolus, respectively (Andersen et al., 2005; Palm et al., 2016; Stenström et al., 2020).

Provided with the extensive understanding of roles of nuclear proteins and nucleome organization, it still remains mysterious about how these nuclear proteins collectively partition, interact and organize in nucleome to drive and mediate the dynamic changes of chromosomal folding, 3-D genome organization, the assembly of nuclear body, the formation of biomolecular condensate and HUB of chromatin-RNA interactions, the histone code encryption and code-deciphering and regulation of gene expression. To address some of these questions from a different perspective, we decided to combine the *in vivo* chemical <u>crossl</u>inking-coupled <u>m</u>ass <u>s</u>pectrometry (XL-MS) (Iacobucci et al., 2019; Liu et al., 2018b; Yu and Huang, 2018) with a MONET-based (Tomasoni et al., 2020) modulomics to profile nucleomic protein-protein interactions (PPIs) and construct a nuclear protein interaction modulome, consisting of hierarchically and topologically organized protein clusters, complexes, subnuclear compartments (including nucleoli, nuclear condensates and nuclear bodies) and nucleosomes. The *in vivo* formaldehyde-fixing of interacting DNA molecules in living cells had been successfully applied in the proximity ligation of DNA fragments (3C, <u>c</u>hromatin <u>c</u>onformation <u>c</u>apture; Hi-C, <u>hi</u>gh-throughput <u>c</u>hromosome conformation capture) to elucidate the comprehensive 3-D genome topology and chromatin fiber interactions (Dekker et al., 2002; Nagano et al., 2013; Quinodoz et al., 2018) and to reveal high resolution architecture features and dynamic changes of chromatin loops and TADs in genome organization (Dekker and Mirny, 2016; Lieberman-Aiden et al., 2009; Misteli, 2020). Similarly, in the split-pool-based SPRITE studies, the *in vivo* formaldehyde-fixed interacting DNA and RNA molecules were adopted to acquire novel information about the interactions (or called HUBs) between chromatins and RNA-participated condensates (Quinodoz *et al*., 2018). Thus, in the present study, the *in vivo* formaldehyde-crosslinking was integrated with the *in organello*-crosslinking of nuclei using CBDPS (Makepeace *et al*., 2020) to capture those nuclear protein-protein interactions prefixed in nuclei (Chavez et al., 2020; Fasci et al., 2018). This double crosslinking-coupled quantitative XL-MS approach (qXL-MS) was considered to be complementary to the classical modulomics approaches where the constructions of protein interaction modules (Lin et al., 2015; Qin et al., 2021) were based on the well-known PPI data repositories, such as STRING and BioGRID (Oughtred et al., 2021; Szklarczyk et al., 2019). These repositories of PPI information were, by and large, acquired from many independent studies using yeast two-hybrid assay (Y2H), <u>a</u>ffinity <u>p</u>urification-<u>m</u>ass <u>s</u>pectrometry (AP-MS), protein complementary assays (PCAs) <u>F</u>örster resonance <u>e</u>nergy transfer, (FRET) and <u>p</u>roximity-<u>d</u>ependent labeling (PDL) (Altmann et al., 2020; Kerppola, 2006; Qin et al., 2021; Rao et al., 2014b), which may generally miss the important quantitative features for computing modulomes using a multi-layer modularity optimization algorithm (M1 algorithm) built in MONET toolbox (Tomasoni et al., 2020). Nevertheless, numerous modulomic analysis have been performed in the past based on the data of <u>p</u>rotein-<u>p</u>rotein interaction (PPI) and module-module interaction (Lin et al., 2012, 2015; Vella et al., 2018) as well as the data of multiple Omics (Silverbush et al., 2019). Especially, the rich data of nucleomic constituents (4DN Data Portal, Dekker et al., 2017) has permitted the development of a novel bioinformatic algorithm, named <u>MO</u>tif <u>C</u>lustering in <u>H</u>eterogeneous Interactomes (MOCHI), to discover the fundamental principles underlying genome organization and gene expression executed by a diverse nucleomic components like chromatins and nuclear proteins (Tian et al., 2020). Recently, the MONET toolbox has been applied successfully in recognizing disease-associated modules from the proteomic data (Johnson et al., 2022). The success of these modulomics exhibited in various applications encouraged us to explore the application of MONET toolbox in construction of nucleomic modulome using a new type of quantitative PPI data acquired from qXL-MS pipeline.

## Results

### qXL-MS analysis of nuclear proteins in nucleome

The qXL-MS-based quantitative interactomics (Fig 1A, S1-S4; See Star Methods for details) was performed on nuclei of a representative eukaryotic organism. The nuclei of this organism were first isolated from 0.2% formaldehyde-prefixed (1^st^ crosslinking) control (or untreated) and hormone-treated tissues, respectively. A second permeable, enrichable and cleavable crosslinker, CBDPS, was subsequently applied (2^nd^ crosslinking) onto these two batches of isolated nuclei samples in order to capture nuclear protein interactions on site. The key feature of this qXL-MS is the *in-organello* ligation by CBDPS of two segments of polypeptides that are in close contact (Fig S2) and already formaldehyde-prefixed in nucleome. Secondly, the *in vivo* formaldehyde-prefixed protein-protein interactions were maintained throughout the nuclei isolation procedure to allow CBDPS to capture what PPIs occurred in living cells. The CBDPS-crosslinked proteins were consequently digested, and the resulting crosslinked peptides (XL-peptides or crosslinks) were dimethyl labelled, mixed, SCX column-fractionated, streptavidin affinity-enriched and LC-MS/MS analyzed to generate 6 experimental replicates of qXL-MS data. As a reference, additional 6 experimental replicates of non-formaldehyde-prefixed but CBDPS-crosslinked nuclei were performed, crosslinks of which were analyzed by an identical LC-MS/MS protocol (Fig 1A, S1-S4). The crosslinks were searched and identified using a modified search engine mXLinkX (Fig S3 - S4; see Star Methods for details).

**Figure 1.**
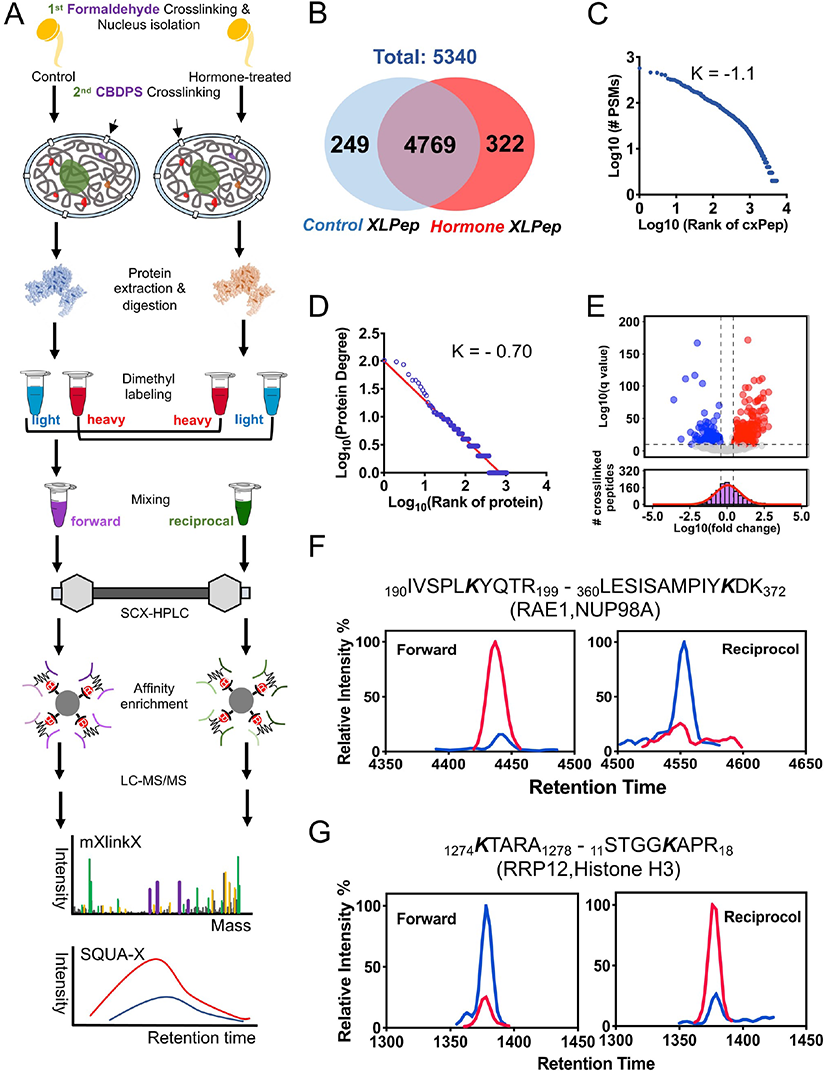
Proteomic profiling of XL-peptides and bioinformatic analysis of PPIs from nucleome. (A) General workflow of quantitative interactomics. The *in vivo* formaldehyde-crosslinked nuclei (1^st^ crosslinking) isolated from both two groups of plant seedlings were further crosslinked with CBDPS (2^nd^ crosslinking). The crosslinked proteins were extracted (Table S0a) and digested into peptides, labeled with isotopic CH_2_O and ^13^CD_2_O. The heavy isotope-labelled peptides from the hormone-treated and the light isotope-labelled ones from the control were mixed equally, defined as the forward mixing experiment (F), whereas the *vice versa* mixing as reciprocal experiment (R). SCX-HPLC stands for strong cation exchange - high pressure liquid chromatography. The XL-peptides were affinity enriched using streptavidin beads. LC-MS was used for analysis of XL-peptides. The mXlinkX (<u>m</u>odified <u>XlinkX</u>) and SQUA-X software was employed for the XL-peptide’s identification and quantification, respectively. (B) A Venn diagram shows the total number of identified control-(5,018) and hormone-related (5,091) XL-peptides, respectively (Table S1d). (C) Logarithm-Logarithm scale distribution of the number of PSMs’ of the XL-peptide over the entire population of XL-peptides (Table S1d). (D) Logarithm-Logarithm scale distribution of the degree of each crosslinked nuclear protein over the entire population of crosslinked nuclear proteins (Table S1j). K represents the slope of the linear regression. (E) Volcano plot (upper panel) and histogram (lower panel) of quantitative interactomics. The base 2 logarithm ratio is the average binary logarithmic ratio of MS1 isotopologue areas of the XL-peptides, while the q is the q-value determined from the Benjamini-Hochberg (BH-FDR) correction of the Student’s t-test. The histogram of base 2 logarithm ratios are fitted using a normal distribution (red curve). The vertical and horizontal dashed line indicates the mean ± 0.5 x SD (standard deviation) of the distribution and the cutoff of q ≤0.1, respectively. The data are listed in Table S1l. (F - G) XIC plot of the up- and the down-regulated XL-peptides of the hormone-treated tissues. Forward and Reciprocal stands for the mixing replicate of the light isotope-labeled control XL-peptides and the heavy isotope-labeled hormone-treated XL-peptides as well as the mixing replicate of the heavy isotope-labeled control XL-peptides and the light isotope-labeled hormone-treated XL-peptides, respectively. Bolded K marks the crosslinking site within an identified XL-peptide. Red lines represent the heavy isotope-labelled XL-peptides while the blue lines represent the light isotope-labelled XL-peptides. The number at the N-terminal end of XL-peptide indicates the amino acid position within a protein. (F) and (G) shows both the XL-peptides derived and XIC measured from RAE (Ribonucleic Acid Export 1)-NUP98A (Nucleoporin 98A) protein pair and RRP12 (Ribosomal RNA Processing 12)-H3 (Histone H3) protein pair, respectively. Data are listed in the Table S1l.

Eventually, a total of 9,289 crosslinks were identified from 12 experimental replicates (Table S1a-c; Fig S5A - C). Over 83.4% (7,754 out of 9,289) of crosslinks were identified from 6 experimental replicates (Fig S5B), indicating a relatively lower cost-effectiveness for the additional 6 experimental replicates in term of crosslinks’ identification. Given the advantage of double crosslinking in mapping of the *in vivo* PPI of nucleomic proteins, both bioinformatics and modulomics were only focused on 5,340 crosslinks derived from the formaldehyde-prefixed cells (Fig S5C), corresponding to 2,540 putative crosslinked nuclear proteins (Table S1d; Fig S5D). The differential interactomic analysis revealed 322 and 249 crosslinks from the untreated control and the hormone-treated samples, respectively (Table S1d; Fig 1B). The relationship between the frequency of PSM (or crosslink) identification and the ranking of each crosslink could be described by a Zipf’s law of a slope of −1.1 (Table S1d; Fig 1C). These crosslinks can be construed into protein-protein interactions (PPIs) according to a standard protocol (Liu et al., 2018a; Yu and Huang, 2018; See Star Methods for details). Filtering of these 2,540 putative crosslinked proteins with plant nuclear proteomes eventually confirmed 1,427 nuclear proteins (Table S1e-f; Fig S5E; Cooper et al., 2011; Hooper et al., 2017; Mair et al., 2019; Yin and Komatsu, 2015), among which 1,052 (74%) nuclear proteins of discrete polypeptide sequences participated in 1,297 hetero-PPIs (Table S1h-j; Fig S5D-E). The protein degrees calculated from the 1,297 hetero-PPIs also followed the Zipf’s law of a slope of −0.70 (Table S1j; Fig 1D). Differential interactomics found 1,155 and 1,181 PPIs from the untreated control and hormone-treated samples, respectively (Fig S5F). These 1,052 nuclear proteins were found to be functionally enriched (by GO enrichment analysis) in both mRNA- and chromatin-binding, and most abundant in functions of mRNA-binding, catalytic activity on RNA, DNA-binding, DNA-binding transcription factor and transcription regulator activity (Table S1m; Fig S6).

To identify hormone-specific nucleomic PPIs, 1,259 quantifiable and dimethyl-labeled crosslinks were selected from 3283 nuclear protein-specific crosslinks (filtered using plant nuclear proteome repository; Table S1l; Fig S5G). Quantification was performed using SQUA-X software on these crosslinks (Table S1l; Fig 1E-G; See Star Methods for details). As a result, there were 129 and 217 hormone down- and up-regulated crosslinks (including both intra- and inter-crosslinks; Fig S7) identified, respectively (Table S1l; Fig 1E, S5G), comprising of 115 and 121 nuclear proteins (Table S1l; Fig S5G), out of which 24 hormone down- and 48 hormone up-regulated inter-crosslinks were found and subsequently constructed into 20 Control- and 35 Hormone-specific PPIs, respectively (Table S1l; Fig S5G).

These 48 Hormone-specific proteins were found to participate in a broad number of nuclear events, including wound hormone biosynthesis process, the chromatin-binding, regulation of DNA and rRNA methylation, nucleosome assembly, negative regulation of gene silencing, snoRNA-binding, copper ion-binding and formation of nuclear pore, preribosome, Cajal body, cytosolic small ribosome subunits and nuclear speckles according to GO enrichment analysis (Table S1n; Fig S8).

### Interactome of histone octamer

The core histone octamer complex is made of two sets of histones proteins, H2A, H2B, H3 and H4. The histone writers and erasers as well as reader proteins interact with these core histones to regulate 3D conformation of chromatins and to participate in the control of the accessibility of regulatory proteins to DNA-surrounding histone octamer (Zhou et al., 2019). In this study, the qXL-MS profiling successfully captured 287 crosslinks from four core histone subunit proteins, among which there were 87 inter- and 200 intra-crosslinks (Table S2a; Fig S9A-B). The quantitative interactomics subsequently found that a majority of hormone significantly regulated crosslinks are intra-crosslinks (10 each of down- and up-regulated intra-crosslinks, Table S2a; Fig S9A-B), suggestive of hormone-induced conformational changes in histone octamer. Moreover, the qXL-MS-based interactomics identified 436 inter-crosslinks between histone subunits and other histone-binding proteins, which gave rise to 256 unique interactors of core histone octamer (Table S2b; Fig S9B), 250 of which were novel. Most interactors interacted with only one subunit of histone octamer complex (Fig S9B). The number of interactors on H2A, H2B, H3 and H4 protein was 51,165, 51 and 29, respectively. It is believed that both the protruding tail of H2B on chromatin and the acidic motif located on H2A-H2B surfaces (Fig 2; Arya and Schlick, 2006; Davey et al., 2017) may have made it possible for the highest number of interactors to be found on H2B (Fig S9B).

**Figure 2.**
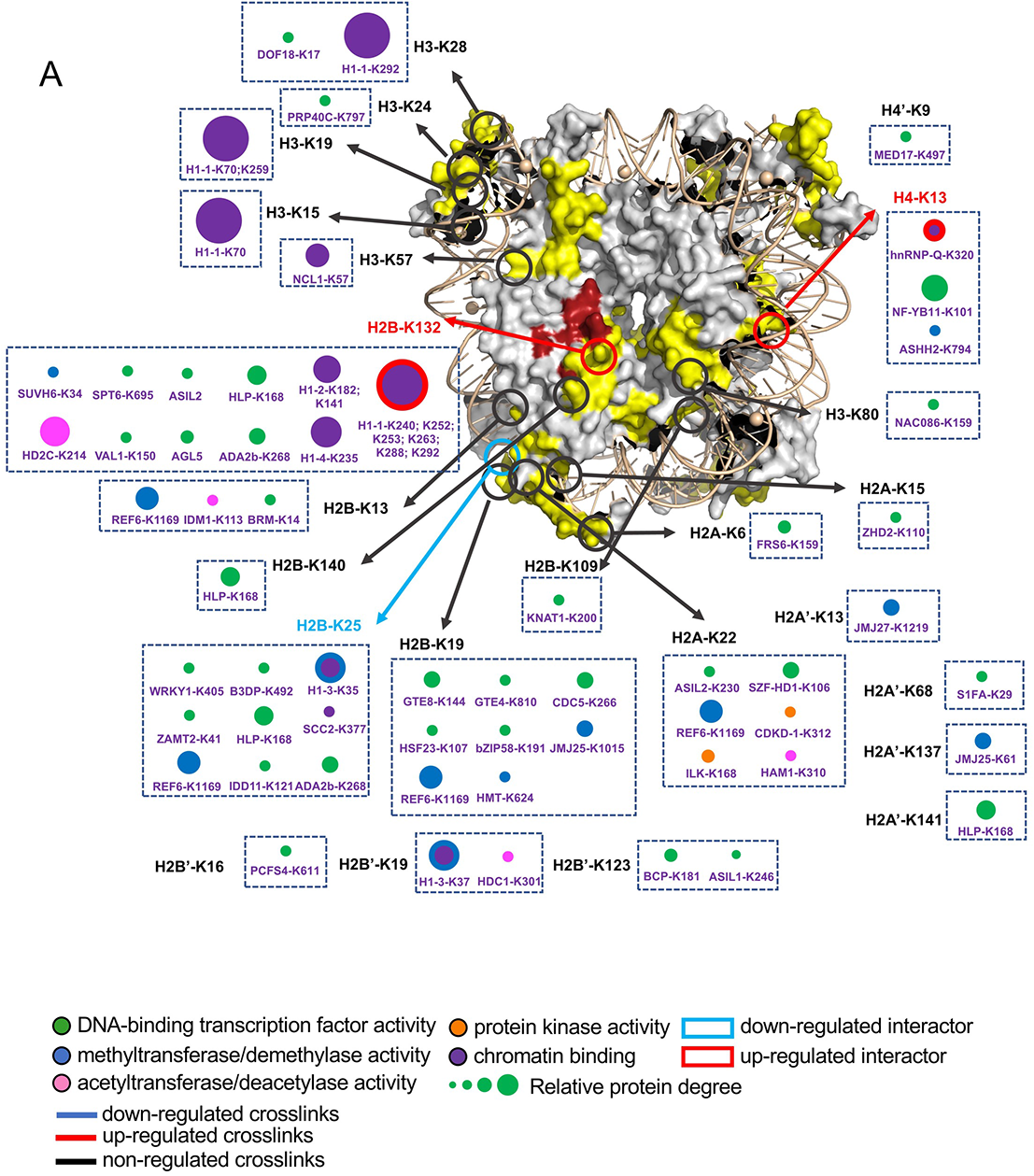
The interactome of histones. The surface representation depicts the histone octamer and DNA double helix, which was built using the homology modeling tool SWISS-MODEL using the X-ray structural data of the chicken nucleosome particles (PDB: 1EQZ). The yellow regions on octamer’s surface mark the XL-MS identified polypeptides from this study. Red region marks the acidic patch of histone octamer (Davey, et al., 2017). The circles and arrows indicate the crosslinking lysine sites on histone proteins marked by black letters. The red circles and arrows as well as blue circles and arrows indicate the hormone up- and down-regulated crosslinks, respectively (Table S1l, S2c). The interactors of histone proteins are shown as colored dots. A group of interactors that link with the identical sites of a histone are enclosed in a dashed box. The interacting lysine sites on interactors are labelled in purple letters. The transcription factor, methyltransferase/demethylase, acetyltransferase/deacetylase, protein kinase and chromatin-binding protein of histone interactors is marked in green, dark blue, pink, orange and dark purple, respectively. The hormone down- and up-regulated interactors (or dots) are marked with blue and red borders, respectively. The size of the node (interactor) represents the protein degree. The data are listed in Table S2c.

These interactors are functionally classified into 29 transcription factors, 12 histone-modifying enzymes *i.e.* 6 methyltransferases/demethylases, 4 acetyltransferase/deacetylases and 2 protein kinases as well as 7 chromatin-binding proteins (Table S2c, Fig 2). A majority of these enzymes and modifiers were linked with the N-terminal tail of histone H2A at K6, K15 and K22 site, H2B at K13, K19 and K25 site and H3 at K15, K19, K24 and K28 site, whereas within the acidic patches of nucleosome, these modifiers were linked with histone H2B at K132 and K140 site and H3 at K57 site (Table S2c; Fig 2; Kalashnikova et al., 2013). Both histone-modifying enzymes and transcription factors have shown a preference in connection with H2A-H2B dimer rather than H3-H4 dimer. Among all these crosslinking sites, K22 site of histone H2A and K19, K25 and K132 sites of H2B were hot spots. Enzymes crosslinking with K22 of H2A were two protein kinases, ILK1 and CDKD-1, and HAM1, a Histone Acetyltransferase of the MYST family 1. This group of enzymes may be involved in both phosphorylation and acetylation of H2A, which is well-known to play a critical role in response to DNA damage repair (Downs et al., 2000). At the hot spots, K19, K25 and K132, of H2B, we found 15 crosslinked transcription factors, 5 chromatin-binding proteins, 4 methyltransferases/demethylases and 1 acetyltransferases/deacetylases. The biological functions of these proteins were highly enriched in regulation of gene expression, whereas proteins involved in chromosome organization processes were two transcription factors, ADA2b and SPT6, two chromatin-binding proteins, the histone H1 and SCC2, and a demethylase REF6 (Fig 2).

Generally speaking, there were three interactors, *i.e.* chromatin-binding protein Histone 1.1 (H1.1; Fig S10A), a transcription factor HLP (Fig S10B) and a REF6 demethylase (Fig S10C), that were of the highest frequency in crosslinking with nucleosome (Table S2c). The hot spots of H1-1, HLP, and REF6 in crosslinking with histone octamer located at C- (9 out of 12), N- (5 out of 5), and C-terminus (4 out of 4) of these three interactors, respectively (Fig S10A-C). Among them, REF6 is a lysine-specific histone demethylase and a regulator of transcription of hundreds of genes in response to various stimuli (Cui et al., 2016). Especially, it is known that REF6 acts as a key mediator in the recruitment of Brahma (BRM) to its target gene loci (Fig 2; Li et al., 2016), and the qXL-MS indeed captured both REF6 and BRM linked to K13 site of H2B.

The CDKD-1 is a protein kinase reported not only to catalyze the phosphorylation of serine at the C-terminus of Arabidopsis RNA Polymerase II to regulate transcription (Hajheidari et al., 2012) but also being involved in cell cycle regulation and cell differentiation (Sterken et al., 2009). In this study, we found that CDKD-1 was crosslinked with histone H2A, demonstrating its possible participation in regulation of phosphorylation of histone H2A. Furthermore, the crosslinks between histone H4 and a heterogenous nuclear Ribonucleoprotein Q (hnRNPQ, or LIF2) were found to be up-regulated by hormone (Table S1k, S2c; Fig 2). Given that hnRNPQ is rapidly recruited to chromatin in response to <u>me</u>thyl <u>ja</u>smonate (MeJA), a wound-responsive hormone, and subsequently mediates the gene activation in Arabidopsis (Molitor et al., 2016), we believe that hnRNPQ protein may play a role in transcriptional switches in the hormone response of plant.

### Interactome of nucleolar box C/D snoRNP complex

The box C/D snoRNP complex is located within the dense fibrillar component (DFC) region of nucleolus and composed of four core subunit proteins, Fibrillarin (FIB), Nucleolar Protein 58 (NOP58), Nucleolar Protein 56 (NOP56) and Small Nuclear Ribonucleoprotein 1 (SNU13). In this study, we identified 33 intra-crosslinks on three subunits, FIB, NOP58 and NOP56 (Table S3a; Fig 3A, S11A) and 11 inter-crosslinks from four subunits of box C/D snoRNP (Table S3a; Fig 3A, S11B). Quantification of these crosslinks revealed 2 hormone down-regulated (2 intra-crosslinks) and 5 up-regulated crosslinks (3 inter-crosslinks and 2 intra-crosslinks; Table S1k, S3a; Fig 3, S11A-B), suggestive of a conformation change in the DFC sub-compartment under hormone treatment.

**Figure 3.**
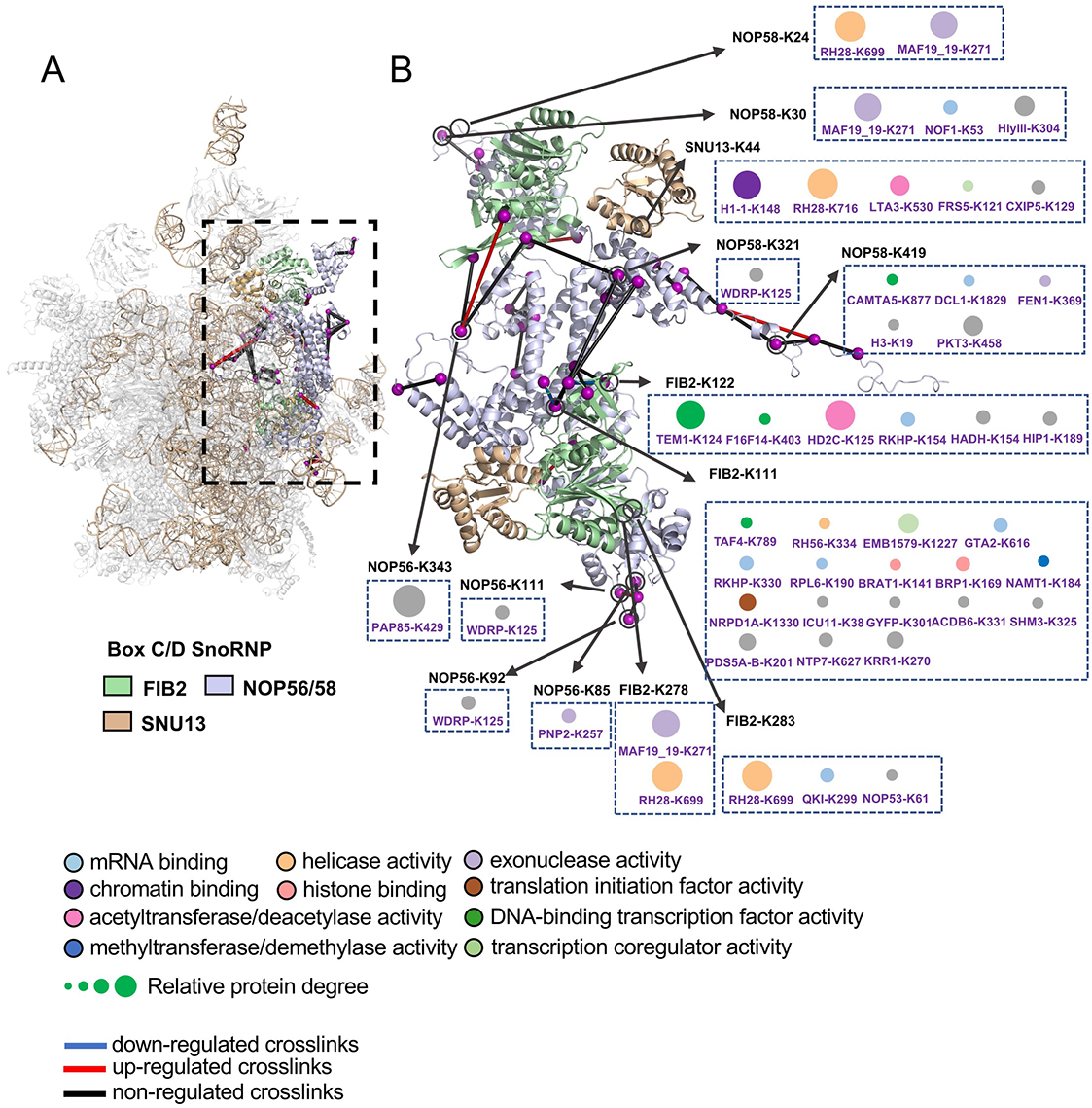
The interactome of box C/D snoRNP complex. The ribbon structure depicts 90S pre-ribosome (A) and the close-up view depicts the box C/D small nucleolar ribonucleoprotein (snoRNP) complex (B), which were built using the homology modeling tool SWISS-MODEL using the Cryo-EM structural data of the 90S pre-ribosome from *Chaetomium thermophilum* (PDB: 5OQL). The RNA and proteins on the 90S pre-ribosome complex are marked in grey and yellow, respectively, while the rRNA 2’-O-methyltransferase fibrillarin (FIB), nu cleolar protein 56/ nucleolar protein 58 (NOP56/NOP58) and small nuclear ribonucleoprotein 13 (SNU13) of box C/D snoRNP is marked in light green, light purple and light brown, respectively. The blue, red and black line within the complex indicates the hormone down-, up-, and non-regulated crosslinks, respectively (Table S1l, S3a-b). The purple balls, black circles and arrows indicate the crosslinking lysine sites on box C/D snoRNP proteins marked by black letters. The interactors of box C/D snoRNP proteins are shown as colored dots. A group of interactors that link with the identical sites of a box C/D snoRNP protein are enclosed in a dashed box. The interacting lysine sites on interactors are labelled in purple letters. The mRNA binding, chromatin binding, helicase activity, translation initiation factor activity, DNA-binding transcription factor activity, histone binding, transcription coregulator activity, acetyltransferase/deacetylase activity, methyltransferase/demethylase activity and exonuclease activity proteins are marked in light blue, purple, wheat, brown, dark green, light coral, light green, pink, dark blue and light purple, respectively. The size of the node (interactor) represents the protein degree. The data are listed in Table S3a-b.

Other than those crosslinks occurred among subunits of box C/D snoRNP complex, there were 57 inter-crosslinks found between the nucleolar protein complex and the complex-interacting proteins. A total of 41 interactors were derived from these inter-crosslinks (Table S3b; Fig 3B, S11B), among which 26 interactors were novel according to previously reported Arabidopsis nucleolus proteome (Palm et al., 2016). FIB interacted with 27 outside protein interactors while the remaining three components, NOP58, SNU13 and NOP56, had 8, 5 and 4 interactors, respectively. The hot spots of FIB for physical interaction with interactors were K111 and K122 sites, which were crosslinked with 17 and 6 proteins, respectively. It was interesting to find that all these FIB’s interactors were crosslinked to sites outside of the intrinsically <u>d</u>isordered region (IDR, or called a glycine- and arginine-rich domain, GAR domain; Fig S11C) even though some intra-crosslinks still occurred in the IDR region (Table S3a-b; Fig 3B, S11C). This may be due to the fact that GAR domains of FIB, which participated in condensate formation, were lack of ample lysine residues. These two features of GAR domain prevented the crosslinking to occur (Yao et al., 2019). In the case of NOP58, its hot spot is K44, which had 5 interactors, while the rest lysine sites K30, K24 and K321 had 3, 2, and 1 interactors, respectively. All 5 interactors of SNU13 were crosslinked at K44 site. NOP56 had 4 interacting sites, K85, K92, K111, and K343 (Table S3a; Fig 3B). One interactor crosslinked at a site. These 41 interactors were all classified into proteins possessing the functions of mRNA-binding, chromatin-binding, acetyltransferase/deacetylase activity, methyltransferase/demethylase activity, helicase activity, exonuclease activity, translation initiation factor activity, DNA-binding transcription factor activity and transcription co-regulator activity (Table S3b; Fig 3B, S11B).

As a prominent interactor of the box C/D snoRNP complex, the DEAD-box ATP-dependent RNA Helicase 28 (RH28) was simultaneously crosslinked with FIB, NOP58 and SNU13 subunits, consistent with the reported protein interactions occurred between RH28 and NOP58 or FIB2 (Krogan et al., 2004). This RNA helicase activity is required for an efficient 2′-O-methylation and mediates the dynamics of box C/D snoRNP complexes on pre-ribosomes (Aquino et al., 2021). It may work in concert with Histone Deacetylase 2C (HD2C) and another interactors (Fig 3) to regulate the activity of FIB rRNA 2’-O-methyltransferase (Chen et al., 2018).

### Hierarchical and topological organization of nucleomic modulome of protein interaction

To map the landscape of the organization of modules of protein interaction in nucleome, 1,297 soybean nuclear hetero-PPIs, comprising of 1052 nuclear proteins, were firstly converted into the corresponding 1,211 Arabidopsis orthologous hetero-PPIs (Table S4a-b) containing 877 Arabidopsis orthologous proteins (Table S4a; Fig S5D-E) because of the availability of the well-known Arabidopsis interactome database (BioGRID) and the convieneint validation system for the functions of nuclear proteins (McWhite et al., 2020; Oughtred et al., 2021). Subsequently, these highly selected PPIs were subjected to modulomic analysis by MONET toolbox (see Star Methods for details), in which the abundance of PPIs (the intensity of edge, defined by the number of crosslinks) played a determinate role in capturing 95 master <u>n</u>uclear <u>p</u>rotein interaction <u>m</u>odules (NPIMs, ≥ 2 components per module; Table S4c; Fig 4, S13) from the intricately organized nucleome.

**Figure 4.**
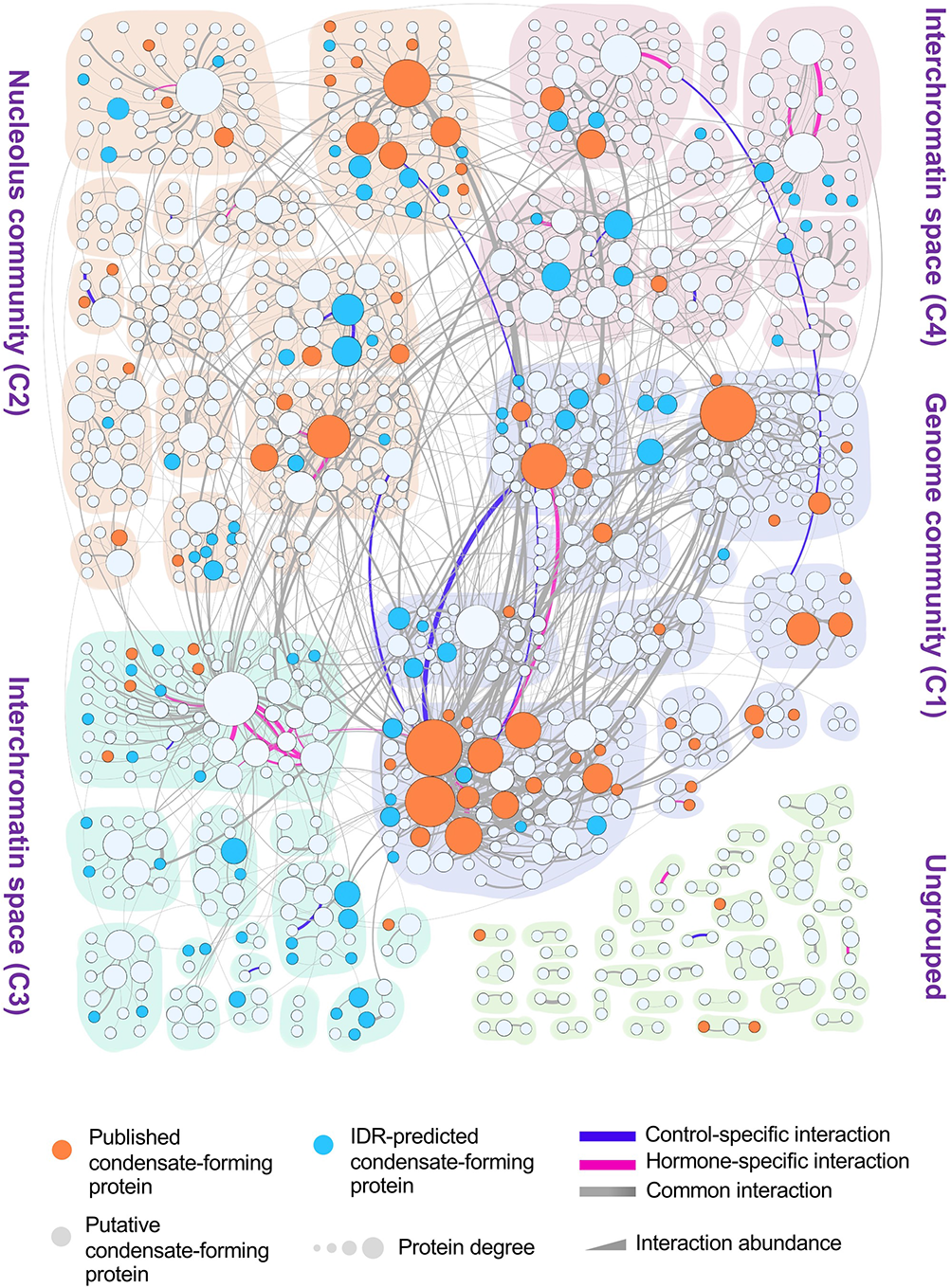
Hierarchical organization of nuclear proteins into NPIMs and communities in nucleome. Diagram shows the classification and organization of 877 highly conserved nuclear proteins into 95 nuclear protein interaction modules (NPIMs) and 4 communities in nucleome using MONET toolbox (Table S4c). The NPIM is defined as a master module, which can be reorganized into module variants temporospatially and conditionally. The higher order module of NPIMs is defined as community. The C1 (light purple) and C2 community (light brown) is defined as the genome community and nucleolus community, respectively. The C3 (light cyan) and C4 community (light pink) represents NPIMs that might locate within interchromatin spaces. The unclassified and non-community forming NPIMs are marked in light green. The dot and line stand for the protein node and the interaction edge within a protein-protein interaction graph, respectively. The size of the node represents the degree of protein. The thickness of line represents the abundance of interaction between two leading proteins The orange, light blue and light grey node represents the literature-annotated condensate-forming, the intrinsically disordered region (IDR)-containing and unknown putative condensate-forming proteins, respectively. The blue, red and grey edge represents the control-specific, hormone-specific, and common protein interaction, respectively.

To discover if there exists a higher order of architecture of NPIMs, *i.e.* <u>m</u>odule-<u>m</u>odule interaction (MMI) in the nucleome graph (Fig S14), the converging nodes and hybrid edges were used by MONET toolbox to generate a module of modules, named as “Community” (Fig S14). Interestingly, a total of 55 master NPIMs were eventually constructed into four higher-order communities (Fig 4, S15). As shown in Fig S15, the Community 1, 2, 3 and 4 (or called C1, C2, C3 and C4) comprises of 17, 15, 14, and 9 NPIMs, respectively (Fig 4). To gain insights into the functions of four nucleomic communities, GO enrichment analysis were performed on the highly interactive modular proteins of these four communities (Table S4d; Fig S16), revealing that nuclear proteins in Community 1 (C1) were largely involved in chromosome organization, chromatin assembly and DNA packaging by the Biological Process, localized in chromatin, chromosome, protein-DNA complex and nucleosome by the Cellular Component and finally specialized in protein chromatin-binding, methylated histone-binding, histone demethylase activity and DNA-binding transcription factor activity by the Molecular Function. Thus, the C1 community was defined as Genome Community (Fig 4, S15). In contrast, nuclear proteins in C2 community were enriched in ribosome biogenesis, rRNA methylation, regulation of gene silencing, and box C/D RNA processing by the Biological Process, in box C/D RNP complex, pre-ribosome and nucleolus by the Cellular Component and eventually in RNA methyltransferase activity by the Molecular Function. It was therefore defined as Nucleolus Community (Fig 4, S15). As to the C3 community, the enriched protein functions include RNA polymerase II binding-related functions and gene expression process, suggesting a regulatory role played by the C3 nuclear proteins in gene expression. However, C4 community did not show any functional enrichment. Continued acquisition of PPI data from nucleome may help clarify the major functions of NPIMs in this community. Taken together, the integration of qXL-MS-based interactomics with MONET-supported modulomics, named as CHAMPION (<u>C</u>apturing the <u>H</u>ierarchically <u>A</u>ssorted <u>M</u>odules of <u>P</u>rotein Interaction in the <u>O</u>rganized <u>N</u>ucleome) pipeline, is able to spatially localize nuclear proteins into four hierarchically and topologically assorted communities of discrete nuclear functions in nucleome. Modules of proteins in Genome Community might be topologically confined to chromosomal territories while modules of proteins in either Nucleolus Community or C3 and C4 communities may be compartmentalized into interchromatin spaces (Fig 4, S15).

The next issue is to elucidate whether the components of a NPIM represent a cluster of nuclear proteins either performing the same nuclear events or assembling into the same physical complexes. By comparing NPIMs with those well-documented protein complexes, condensates and nuclear bodies involved in various nuclear events, it was interesting to find that there were six NPIMs overlapping with the well-reported protein complexes spatially disseminated into four PPI communities of nucleome (Table S4c). For examples, all components of NPIM4-42 were actually subunits of DNA-dependent RNA polymerases II, suggesting that this module plays a role in transcription, whereas most components of NPIM4-56 were found to contain subunits of Prp19 complex involved in both activating the precatalytic spliceosome and catalyzing splicing. Moreover, the components of both NPIM0-47 and 0-77 overlapped with the protein components of proteasome while NPIM2-57 and 0-63 overlapped with those of 40S ribosome and small nuclear ribonucleoprotein complex, respectively (Table S4c).

In addition, we also found components of many NPIMs matching with those of diverse nuclear bodies, including Cajar body, dicing body, nuclear speckle, and photobody (Table S4c). For an example, NPIM2-7 contains the core components of box C/D snoRNP complex playing functions in preribosomal RNA methylation, among which FIB is a key member of the center for rRNA processing (Xing et al., 2017; Yao et al., 2019) and the N-terminal GAR domain of FIB protein is essential and sufficient for LLPS (Berry et al., 2015; Feric et al., 2016). Interestingly, several additional components of NPIM2-7 are also well-known to be involved in RNA-processing. They include several RNA helicases (*i.e.* RH28, RH56 and DCL1), rRNA processing proteins (*i.e.* NOP53, 56 and 58, KRI1 and NHP2) and other RNA-binding proteins (*i.e.* GTA2 and KH26) as well as CP2 and DCL1 proteins involved in RNA-processing during gene-silencing. It is therefore conceivable that NPIM2-7 indeed captured a nuclear body consisting of a cluster of nuclear proteins involved in RNA-processing, which may play a role in cell cycle control as some components of this module, for instances, CDC48PR and CCAR1, are well-known to participate in cell cycle control (de los Santos-Velázquez et al., 2017).

Since unravelling of protein composition of a condensate is still a challenging issue, the results of CHAMPION workflow were utilized to decipher the identity of protein molecules in various condensates. For another example, Super-Enhancers (SEs) were clusters of enhancer proteins, collectively assembled into high density transcription apparatuses to promote robust gene expression (Sabari et al., 2018). Previous studies have shown that SE condensates facilitate compartmentalization and concentration of transcription components at the specific gene sites, which is facilitated by the phase-separating properties of IDRs on transcription factors and cofactors. MONET-based modulomic studies revealed two cases where a NPIM may contain several components of a condensate. In NPIM2-4, both GTE6 (an ortholog of human Bromodomain-containing protein 4) and MED1 have been documented to form phase-separated droplets together, which consequently sequestrate other transcriptional factors to boost gene expression (Table S4c; Sabari et al., 2018), whereas in NPIM0-53, HAC1 of p300/CREB-binding protein family interacted with WRKY3. These components have been documented previously to drive co-condensation and regulate transcriptional-bursting and co-operative gene expression (Table S4c; Ma et al., 2021).

For another example, out of three transcription coactivators (by GO annotation), MBF1B, MED30 and HAM1, modulomics was able to partition all three modular components into NPIM1-8, in which MBF1B and MED30 were indeed found to interact with each other (Table S4c; Fig 4). At the same time, more transcription factors like S1FA, lysine-specific demethylase JMJ25 and the trihelix transcription factor ASIL2L as well as a subunit of Paf1/RNA Polymerase II Complex CTR9 were all colocalized into NPIM1-8 (Table S4c; Fig 4). The enrichment of these transcription factors and coactivators into the same module (NPIM1-8) supports that modulomic analysis may indeed capture novel scaffold and client condensate-forming proteins into a putative SE condensate. Similarly, another observation was made on NPIM1-9, in which found were two transcription coregulators, *i.e.* CAMTA4 and SWRC4 as well as three transcription factors, *i.e.* HMGB1, RR2 and the tri-helix transcription factor ASIL2. All these transcription regulators possess the function of gene expression regulation, suggestive of a potential role for these components in forming a putative SE condensate (Table S4c), especially that the HMGB1 is well-known to contain a large Intrinsically Disordered Region (IDR), which may serve as the contact point between condensate-formation of proteins (Nott et al., 2015), further supporting the hypothesis that NPIM1-9 may capture components of a putative condensate.

By further comparing components of NPIMs with the documented condensate-forming proteins, we found that each of 27 NPIMs (28% of the total NPIMs) contain at least one condensate-forming protein (Table S4c; Fig 4, S15A). For example, 13 NPIMs (NPIM0-33, 0-59, 0-63, 0-75, 1-3, 2-4, 1-14, 2-5, 3-39, 3-41, 3-89, 4-6 and 4-10) consisted of proteins localized in nuclear speckles that are both known as interchromatin granule clusters or splice speckles (Table S4c) and involved in gene regulation, RNA metabolisms, pre-mRNA-processing and mRNA export. Furthermore, two Phytochromes (PHYs) proteins, PHYA and PHYE, were found in NPIM3-2 and NPIM3-27, respectively (Table S4c). These phytochromes (PHYs) are localized in photobodies (or plant-specific nuclear bodies) containing photoreceptors and condensate-forming components that play a role in mediating the light signal transduction. Moreover, NPIM1-1, 1-3, 4-6, 1-8, and 1-36 contain core histone proteins, revealing a possible correlation with chromatin formation, which has been reported to have an intrinsic LLPS capability. Among these modules, NPIM4-6 contains the histone variant H2AX required for an efficient repair of DNA double-strand breaks when modified by C-terminal phosphorylation. The LHP1 in NPIM2-4 is an ortholog of human HP1α, which has been shown to promote the compartmentalization of constitutive heterochromatin (Larson et al., 2017; Strom and Brangwynne, 2019). Recent reports also demonstrated that histone H1, the most interactive protein in NPIM2-5, can go through the LLPS when its lysine-rich C-terminal tail is mixed with DNA, promoting condensation of heterochromatin (Gibson et al., 2019; Turner et al., 2018). In addition, NPIM1-9, 1-17, and 1-25 consist of known transcription factor-associated condensates, whereas NPIM1-18 is composed of several GBPLs that were found to undergo phase-transition to control transcriptional responses. Plant La1 protein in NPIM3-43 is an ortholog of yeast La protein 1 (La1). It may form droplets upon association with RNA. La1 is also a factor of ribosome biogenesis that functions essentially in rRNA processing (Palm et al., 2019). Furthermore, both the interactive proteins, CRWN1 and 4, of NPIM2-13, which are the plant nuclear lamina homologs, may form nuclear lamina responsible for the condensation of inactive chromatins in TADs (Ulianov et al., 2019). Many transcriptional repressors, including histone deacetylases in NPIM2-12, were linked to the nuclear lamina (Ulianov et al., 2019). The protein GBPL3 in NPIM1-18 has been reported to form GBPL Defense-Activated Condensates (GDACs) in Arabidopsis (Table S4c). In the same module, discovered were three additional transcription factors, SHP2, HB22 and ULT2 together with a THO5 protein. The molecular functions of these modular components appeared to be consistent with the well-known function of GDAC in recruiting the regulatory proteins to control the gene expression in plant defense (Huang et al., 2021), Thus, it is concluded that NPIM1-18 may captured additional components of the GDAC condensate. These findings strongly suggest that NPIMs may capture various protein assemblies in nucleome.

All these findings also encouraged us to investigate if the MONET-determined NPIM may potentially reveal the protein composition of a novel condensate. Given that proteins of IDR may have a potential to form a condensate, the IDR and the prion-like domain were therefore predicted from all modular protein components (Table S4c) by both MobiDB and PLAAC programs (See Star Methods for details). As a result, 42 NPIMs were found to contain a component possessing the predicted IDR/prion-like domain, out of which 18 NPIMs actually contain both IDR-possessing and condensate-forming protein components (Table S4c; Fig 4, S15A). Interestingly, the percentage of this class of NPIMs (both IDR-containing and condensate-forming) was higher in both C3 and C4 communities than that in both C1 and C2 communities. In contrast, the percentage of NPIMs containing the well-known condensate-forming proteins in both C1 and C2 communities is higher than those in both C3 and C4 (Table S4c; Fig 4, S15A). One example of IDR protein-containing NPIM, which may capture the putative condensate, is the NPIM1-46. In this module, both MAC2 and TMA7 components contain large IDRs (Table S4c). At the same time, there were 3 more components found to associate with the RNA-binding (Table S4c). It is therefore speculated that NPIM1-46 might have captured components of an RNA-related condensate.

### Partitioning of Control- and Hormone-specific module and community variants

To reveal how a hormone treatment might alter the architecture of modules and communities of nucleomic proteins, we segregated the overall PPIs into the untreated Control (20 Control-specific Arabidopsis ortholog PPIs; Table S5a-b) and the Hormone-treated (31 Hormone-specific Arabidopsis ortholog PPIs; Table S5c-d) datasets according to the information of hormone significantly regulated crosslinks (Table S1L). Separate modulomic analysis of these two specific datasets plus 1,160 common PPIs dataset resulted in 79 modules and 4 communities for each of Control and Hormone samples (Table S5e-f; Fig 5A). To identify Control- and Hormone-specific module variants and community variants, we pair-wisely compared the components of two sets of 79 module variants and 4 community variants using Jaccard Coefficient (JC; Table S5g; Fig 5B-C; see Star Methods for details), and consequently identified both 17 Control-specific and 17 Hormone-specific module variants and a pair of Control- and Hormone-specific Community variant (JC < 0.6; Table S5e-g; Fig 5B-C, S17-S18; Tang et al., 2021). The rest 62 modules of JC larger than 0.6 from both Control and Hormone PPI datasets were classified as the Steady-State (unchanged by hormone treatment) Module (Table S5e-g). Comparison of components between Control and Hormone community revealed that the C4 community was altered dramatically by hormone treatment while the C1, C2 and C3 communities remained as the Steady-State Community (Table S5e-f; Fig 5C) according to the JC calculation.

**Figure 5.**
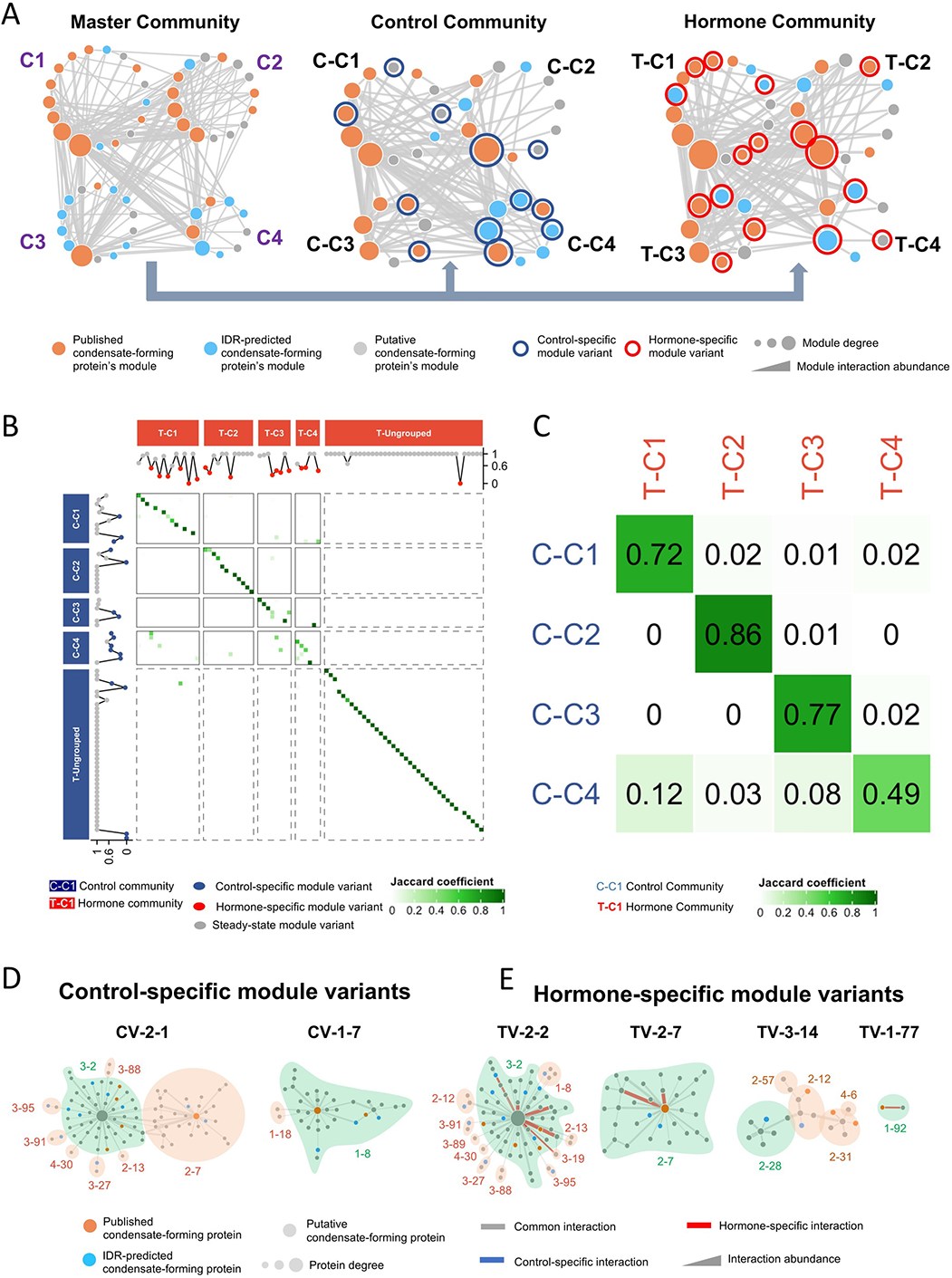
Heatmap and topological analysis of Control and Hormone-specific modules and communities. (A) The schematic topological graph represents 4 communities of master (left panel), Control (middle panel) and Hormone modules (right panel), respectively. The node and edge represents the NPIM (or variant of NPIM) and the interaction among modules (MMI, <u>m</u>odule-<u>m</u>odule interaction) within the nucleus graph, respectively. The higher order module of modules is defined as community. The size of the node represents the degree of module. The orange, light blue and light grey node represents the literature-annotated condensate-forming, the intrinsically disordered region (IDR)-containing and unknown putative condensate-forming modules, respectively. The thickness of line represents the abundance of module-module interaction, which is a count of PPIs between these two modules. The dark blue and red circle marks the control- and hormone-specific module variant, respectively. C1, C2, C3 and C4 represents the community 1, 2, 3 and 4, respectively. The prefixes of communities, “C-” and “T-”, stand for Control and Hormone dataset, respectively. The data are in Table S4c and S5e-f. (B - C) The heatmap represents the module (B) and community (C) comparison between the Control (generated by Control-specific and Common *Arabidopsis* ortholog PPIs; rows) and Hormone (generated by Hormone-specific and Common *Arabidopsis* ortholog PPIs; columns) dataset. The similarity between two modules is evaluated by Jaccard Coefficient (JC), which is determined by the number of identical components divided by the total number of unique components. The green color of the heatmap stands for the level of Jaccard Coefficient. C1, C2, C3 and C4 represents the community 1, 2, 3 and 4, respectively. Ungrouped are those modules that fail to be integrated into a community. The prefixes of communities, “C-” and “T-”, stand for Control and Hormone dataset, respectively. The dark blue, red and grey dots of (B) stand for the Control-specific (JC <0.6), Hormone-specific (JC<0.6) and the steady-state modules variants (JC≥0.6), respectively (Table S5g). (D - E) Representatives of Control- (D) and Hormone-specific (E) module variants. The code (in black) marks the graph index of module variants. The green and brown code indicates the master module graph index of arbitrarily defined scaffold and client submodules, respectively. The green and light brown background color denotes the scaffold and client submodule, respectively. The orange, light blue and light grey node represents the literature-annotated condensate-forming, the intrinsically disordered region (IDR)-containing and unknown putative condensate-forming proteins, respectively. The size of the node represents the protein degree. The blue, red and grey edge represents the Control-specific, Hormone-specific, and Common PPI, respectively (Table S5e-g).

To investigate how the master NPIMs are reorganized into module variants upon a hormone treatment, we named the Control- and Hormone-specific module variant as CV and TV, respectively. For example, both master NPIM3-2 and NPIM2-7 were merged into a Control-specific module variant, CV-2-1, in the Control PPI dataset-assembled communities while they were segregated into two independent hormone-specific module variants, TV-2-2 and TV-2-7, in the Hormone PPI dataset-assembled communities (Table S5e-f; Fig 5D-E, S17-S18). When the protein compositions of CV-2-1 and TV-2-7 were compared, it was found that both module variants contained the protein FIB (originated from NPIM2-7), a key component of nucleolar DFC. The other 4 co-residing chromosome organization proteins (originated from NPIM3-2) like RAD50, UBN1, ARP7 and ARID5, were colocalized into CV-2-1 with FIB. In contrast, these components were missing from TV-2-7 module variant, suggesting that the hormone plays a role in decreasing the interaction of these chromosomal organization-related proteins with nucleolar proteins. As nucleolus plays a role in maintaining the organization of its surrounding genome (Lindström et al., 2018), this finding of modulomics suggested that the hormone may affect spatial organization of nucleolus-associated chromosomal domains. In addition, more RNA-processing proteins (*i.e.* RNA helicase and ribosome biogenesis proteins) were also assembled into the module variant CV-2-1 as comparing to module variant TV-2-7, indicating that the hormone may suppress rRNA biogenesis and ribosome assembly in nucleolus (Table S5e-f). Together, these modulomic results indicated that the hormone influenced both the assembly of nucleolar protein complexes and the chromosome organization around nucleolus. Likewise, another example of module reorganization is TV2-2 variant, in which the chromosomal organization proteins, RAD50, UBN1, ARP7 and ARID5, merged with 3 stress-related proteins, like ILK1, GAPC1 and NDK1, originated from NPIM1-8, NPIM2-12, and NPIM3-19, respectively, to regulate gene expression.

One more example of hormone-triggered module reorganization is TV-3-14 variant, in which 3 condensate-forming proteins, RH45, DYRK2A and AGO10 were sequestrated from NPIM2-12, NPIM2-31 and NPIM2-31, respectively (Table S5e-f; Fig 5D-E, S17-S18). Upon the hormone treatment, both AGO10 (a subunit of RNA-Induced Silencing Complex) and RH42 (a helicase required for pre-mRNA splicing by spliceosome) merged into the variant TV-3-14. As the production of siRNA is required the spliceosome-associated complex (Dumesic et al., 2013), the formation of variant TV-3-14 might be involved in forming a novel Hormone-specific condensate of functions related to activities of RISC and spliceosome. Similarly, hormone also promoted the formation of Hormone-specific module variant TV-1-77, which was composed of only two components, RAE1 and NUP98A. Both of them are well-known mRNA export factors on the nuclear pore complex. The formation of hormone-induced module variant may reflect a hormonal regulation of the mRNA export process *via* modification of nuclear pore subunit composition (Table S5e-f; Fig 5D-E, S17-S18). Taken together, it is evident that the CHAMPION pipeline is able to capture the dynamic reorganization of protein interaction modules in nucleome upon a hormone stimulation.

To further analyze the possible functions of the module variants that are topologically confined to each nucleomic community, we selected 3, 2, 2, and 5 Control-specific module variants from C-C1, C-C2, C-C3 and C-C4 community, respectively, and 6, 3, 4 and 3 Hormone-specific module variants from T-C1, T-C2, T-C3 and T-C4 community, respectively (Table S5e-f; Fig 5A-B) for function comparisons. GO enrichment analysis were separately performed on these two sets of module variants (Table S5h; Fig S19). It was found that in C-C1 community, the components of Control-specific module variants were specifically and functionally enriched in chromatin and nucleosome organization, spliceosome complex, nuclear lumen, catalytic complex, ribonucleoprotein complex, and non-membrane-bounded organelle as expected, whereas those proteins of the Hormone-specific module variants in T-C1 community were enriched in RNA metabolic process, mRNA and RNA-binding, pyrophosphatase activity and nuclear pore, suggesting an association of these hormone-induced modules with gene expression. In C-C2 community, the components of Control-specific module variants were specifically enriched in nuclear body and large ribosomal subunit, while that of Hormone-specific module variants in T-C2 community were enriched in mRNA export from nucleus and sister chromatid cohesion. In C-C3 community, although there was no GO annotation enriched Control-specific module variants, the Hormone-specific module variants in T-C3 community were specifically enriched in mitotic sister chromatid cohesion, nucleic acid-binding, misfolded protein-binding, protein-folding chaperone, snRNA-binding, ATP hydrolysis activity, catalytic activity, cohesion complex, and ribosome. Finally, in C-C4 community, the components of Control-specific module variants were specifically enriched in nucleobase-containing compound metabolic process, peptide biosynthetic process, translation, RNA processing, mRNA-binding, cohesion-loading activity, hydrolase activity, glycogen synthase activity, cohesion complex, ribosomal subunit, intrinsic component of nuclear inner membrane, tricaboxylic acid cycle enzyme complex and amyloplast, while that of Hormone-specific module variants in T-C4 community were enriched in ribosome large subunit biogenesis, regulation of DNA repair, meristem maintenance, regulation of response to DNA damage stimulus, maturation of SSU-rRNA, lipid modification, rRNA processing, oxidoreductase activity, and 90S pre-ribosome.

In summery, the modulomic analysis of 4.2% Control- and Hormone-specific PPIs (20 Control-specific PPIs and 31 Hormone-specific Arabidopsis ortholog PPIs out of 1,211 Arabidopsis ortholog PPIs) produced a total of 21.5% module variants within nucleome graph (*i.e.* 17 Control-specific + 17 Hormone-specific module variants out of 79 Control + 79 Hormone modules (both of which include 62 Steady-State Modules). Bioinformatic GO enrichment analysis of these module variants found that the Hormone-specifically interacting protein clusters (modules) may collectively participate in a broad range of nuclear events from RNA processing to RNA export, ribosome biogenesis, sister chromatids separation and DNA damage repair.

To reveal the extent of functional reorganization of modular components at a community level, the variation and commonality of highly interactive nuclear protein components within each community between Control and Hormone dataset were investigated. To do that, the top 10% and 50% of nuclear proteins of the highest protein degree were first selected out from these two PPI datasets (Table S5i-l; Fig S20). Comparison of these highly interactive protein components within each community by GO enrichment analysis revealed that there existed the greatest commonality between Control and Hormone sample by Cellular Component analysis, whereas a much higher level of variation found in between the two datasets by both GO Biological Process and Molecular Function analysis (Table S5i-l; Fig S20). Among all four communities, the function of highly interactive proteins of C4 community was the most variable between Control and Hormone sample (Table S5i-l; Fig S20), which was consistent with the component comparison among communities conducted above. In contrast, a certain degree of similarity was observed between the C-C1 community and the T-C3 community. Vice versa, the function of the C-C3 community is similar to that in the T-C1 community according to the GO enrichment analysis by Biological Process and Cellular Component (Table S5i-l; Fig S20), suggesting that hormone may promote the protein components’ exchange among communities.

### Confirmation and validation of qXL-MS and modulomics analysis results

To access the reliability of the mXlinkX, we firstly performed the CBDPS-crosslinking of synthetic peptides, Fmoc-KELDDLR and Fmoc-EAKELIEGLPR, and Bovine Serum Albumin (BSA, Table S0b-c; Fig S4). The intra-crosslinks of BSA were mapped to the crystal structure of BSA subunits (PDB: 3V03). Both Cα-Cα distances between pairs of crosslinked lysine residues and those between randomly assigned lysine residues were measured, leading to a different Cα-Cα distance distribution between the crosslinked and randomly assigned lysine residues (Fig S4). However, all of the Cα-Cα distances among the crosslinked lysine residues fell within the maximum constraint imposed by the CBDPS crosslinker (∼38Å; Makepeace et al., 2020). To further confirm the intra-crosslinks, we screened all 533 nuclear proteins containing intra-crosslinks and consequently measured the distances of these crosslinked sites against the data of these proteins deposited in the <u>P</u>rotein <u>D</u>ata <u>B</u>ank (PDB, Berman et al., 2000; Burley et al., 2021) with an identity threshold of 0.9. Eventually, 61 nuclear proteins were selected to measure the Cα-Cα distances between pairs of crosslinked lysine residues (Table S6a; Fig 6A). As a result, the measurement revealed about 99% of the distances agreeing well with the constraint of CBDPS crosslinker (Table S6b; Fig 6B). Taken together, the spatial distances of pairs of crosslinked lysine residues on a protein subunit match well with those measured from their protein structure information.

**Figure 6.**
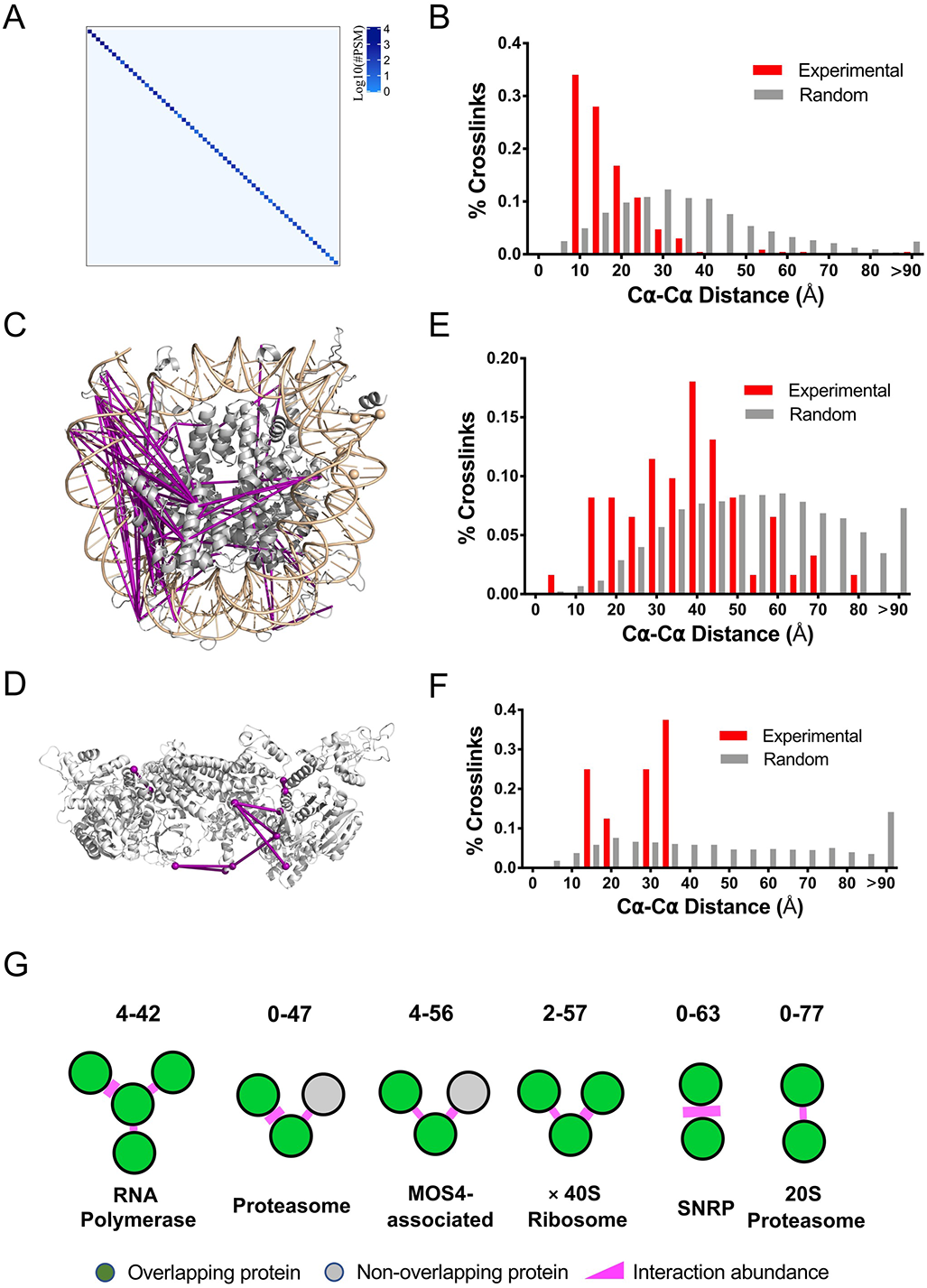
Bioinformatic confirmation of crosslinks and PPI within 3D structure of protein and NPIM. (A) Heatmap represents the intra-crosslinks within each one of 61 nuclear monomeric proteins. The blue color of dots represents the PSM counts of the crosslinks (Table S6a). (B) The distribution of a carbon distances between the crosslinked lysine pairs from intra-crosslinks. The red bars represent the distribution of experimental a carbon distances of the crosslinked lysine pairs while the grey bars represent that of randomly selected lysine pairs (Table S6b). The data of a carbon distances were collected from Protein Data Bank (PDB). (C - D) The ribbon structure and crosslinks of the histone octamer in combination with DNA double helix (C; Table S6c) and box C/D small nucleolar ribonucleoprotein (snoRNP) complex (D; Table S6d). The 3D structures of both protein and protein complex were built by the homologue modeling tool SWISS-MODEL. Purple line and dot represents the crosslink and crosslinked lysine site, respectively. The structural data of histone octamer and snoRNP complex were obtained from the chicken nucleosome particle (PDB: 1EQZ) and the 90S pre-ribosome of *Chaetomium thermophilum* (PDB: 5OQL), respectively. (E - F) The distribution of a carbon distances between the crosslinked lysine pairs from histone octamer (E) and box C/D snoRNP (F). The red bars represent the distribution of experimental a carbon distances of the crosslinked lysine pairs while the grey bars represent that of randomly selected lysine pairs (Table S6e-f). (G) Schematic diagram of 6 NPIMs overlapping to the conserved plant protein complexes. The nodes and purple edges represent proteins and protein-protein interactions, respectively. The green and grey node stands for the overlapping and non-overlapping protein of NPIM with the reported subunits of the conserved plant protein complexes, respectively (Table S6g).

As to the confirmation of inter-crosslinks, we firstly mapped the inter-crosslinks onto the 3D structures of histone octamer and nucleolar box C/D snoRNP complex, which were built by the SWISS-MODEL using chicken nucleosome particle (PDB: 1EQZ) and the 90S pre-ribosome of *Chaetomium thermophilum* (PDB: 5OQL), respectively (Table S6c-d; Fig 6C-D). Among subunits of these two protein complexes, the Cα-Cα distances measured between the crosslinked lysine residues showed a distribution different from that of the randomly selected lysine residues of the protein 3D structures (Table S6e-f; Fig 6E-F). Measurement of 61 inter-crosslinks among subunits of histone octamer structure, 43% of which fell within the constraint of CBDPS (Table S6e; Fig 6E). In contrast, 57% inter-crosslinks have a Cα-Cα distance exceeding the maximal constraint of CBDPS because these inter-crosslinks might actually occur on residues of the flexibly moving histone tail. In terms of 8 inter-crosslinks occurred among subunits of nucleolar box C/D snoRNP complex, all had the Cα-Cα distances falling within the expected constraints (Table S6f; Fig 6F). To further confirm the inter-crosslinks, 1,297 hetero-PPIs obtained from this experiment were searched against both STRING and BioGRID databases (Table S1h). It was found that a total of 77 PPIs had already been documented in these PPI repositories (Table S1h). The third class of confirmation for the inter-crosslinks came from the comparison between the CBDPS-crosslinked proteins and the subunits of the gel filtration-determined protein complexes, such as RNA Polymerase II, 40S Ribosome, MOS4-associated, Proteosome, small nuclear ribonucleoprotein particle (SNRP) and 20S Proteosome (Table S6g; Fig 6G; Makepeace et al., 2020).

Other than the bioinformatic confirmation of hetero-PPIs, we also applied the super-resolution microscopy to validate the hetero-PPIs. A protein-protein interactions both in between hnRNPQ and histone H4 as well as RAE1 and NUP98A were selected. Consequently, we generated four antibodies, anti-hnRNPQ, anti-H4, anti-RAE1 and anti-NUP98A. As a result, a 568 nm dye and a 750 nm dye were employed to label both anti-hnRNPQ and anti-RAE1 antibodies as well as both anti-H4 and anti-NUP98 antibodies, respectively. The dSTROM microscope was used to validate the PPIs of these four interactors (Fig 7A-B, S21A-D, S22-S25). Both hnRNPQ and NUP98A were chosen to serve as the negative control for the pair of non-interacting proteins (Fig S21A). The titration experiments using the anti-hnRNPQ, anti-H4, anti-RAE1 and anti-NUP98A antibodies firstly demonstrated the specificities of these antibodies in recognizing the target proteins (Fig S21B-C). The colocalization (yellow) of hnRNPQ (green) and histone H4 (red) indeed occurred under microscopic observation of both untreated control and hormone-treated plant cells (Fig 7A, S21D, S22-S23). There was a significant difference in the colocalization coefficient between the interacting protein pair, hnRNPQ-H4 and the non-interacting one, hnRNPQ-NUP98A, supporting the PPI of hnRNPQ (green) and histone H4 (red) (Fig 7A, S21D, S22-S23). At the same time, the signals of these two proteins overlapped well with that (cyan) of cell nuclei, further confirming the nuclear localization of these two proteins (Fig 7A). Similar to the hnRNPQ - histone H4 pair, the interaction and localization of RAE1-NUP98A protein pair was also validated by the colocalization and nuclei signals (Fig 7B, S21D, S24-S25).

**Figure 7.**
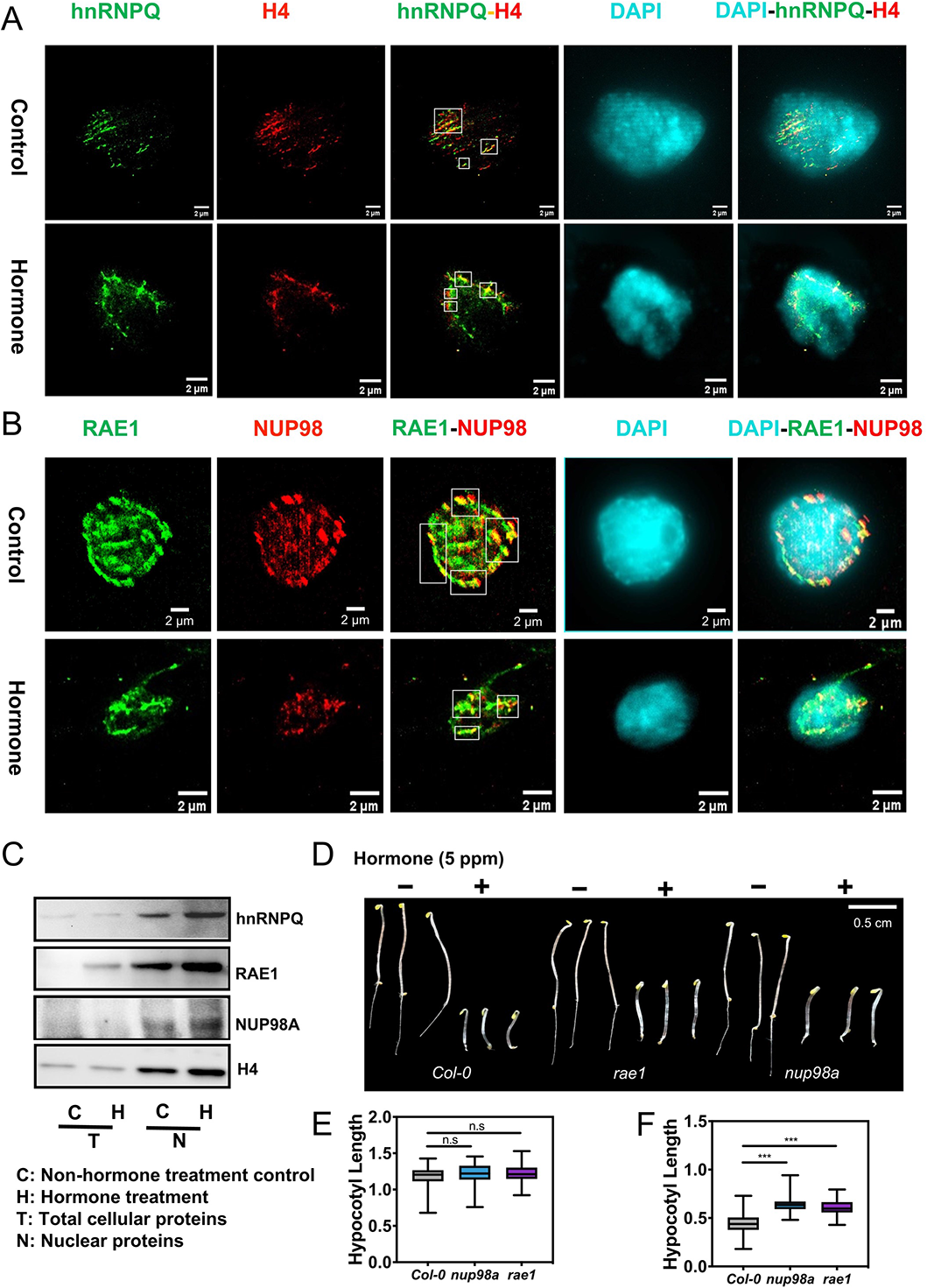
Microscopic confirmation of PPI and validation of hormone-specific module variant. (A - B) STORM super-resolution imaging of PPIs of hnRNPQ with histone H4 and RAE1 with NUP98A. The upper and lower five photos in panels A and B are the imaging results of non-hormone-treatment control and hormone-treated nuclei, respectively. Both hnRNPQ and RAE1 proteins are shown as green color (from Alexa Fluor 568 dye), whereas H4 and NUP98A proteins are shown as red color (from Alexa Fluor 750 dye). The colocalized protein pairs are shown in yellow color. DAPI-stained nuclei are marked in cyan color. (C) Western blot analysis of proteins extracted from nuclei of both non-hormone treatment control and hormone treatment tissues. Protein hnRNPQ, H4, RAE1 and NUP98A was identified using anti-hnRNPQ, -H4, -RAE1, and -NUP98A polyclonal antibodies, respectively. C and H stands for proteins extracted from control and hormone-treated tissues, respectively. T and N stands for the total cellular and nuclear proteins, respectively. (D) Triple response phenotypes of the loss-of-function mutants, *rae1* and *nup98a*, of the model plant *Arabidopsis*. Representative phenotypes of the etiolated seedlings of 4-day-old *Col-0*, *rae1* and *nup98a Arabidopsis* grown without (-) and with hormone treatment (+). (E - F) Measurement of *Arabidopsis* hypocotyl length under non-hormone treated control (E) and hormone treated condition (F). The grey, blue and purple box represents *Col-0*, *rae1* and *nup98a*, respectively. Three biological replicates were performed. Average values and error bars (±SEM) are shown. n.s., *, **, and *** as p ≥ 0.05, p < 0.05, p < 0.01, and p < 0.001, respectively.

In addition, the immunoblot assays were also employed to validate the hormone up-regulation of the inter-crosslinks occurred in between hnRNPQ and histone H4 as well as in between RAE1 and NUP98 (Fig 7C, S26). The protein histone H4 served as a loading control for both untreated Control and Hormone-treated samples (Fig 7C, S26). The protein amount changes of hnRNPQ, RAE1 and NUP98A were quantitated accordingly (Fig 7C, S26). As a result, the protein levels of hnRNPQ, RAE1 and NUP98A were increased under Hormone treatment as compared with the untreated Control, which was consistent with the qXL-MS results (Fig 7C, S21E, S26).

To confirm the NPIMs generated by modulomic analysis, we compared the NPIMs with the well-known established plant protein complexes and found that NPIM4-42, 0-47, 4-56, 2-57, 0-63, and 0-77 were highly overlapping with the gel filtration-determined protein complexes: RNA Polymerase, 40S Ribosome, MOS4-associated, Proteosome, SNRP, and 20S Proteosome complexes, respectively (Table S6g; Fig 6G). In addition, it was found that NPIM 2-4 and 0-53 contained the co-condensation components of established SE condensates (Table S4c). Taken together, MONET-based modulomics had successfully captured a portion of components of both stable protein complexes and the biomolecular condensates.

Finally, the biological role of a Hormone-specific module variant, TV-1-77, containing RAE1 and NUP98A, was investigated by a seedling hormone triple response assay on the wild type *Col-0* as well as *rae1* and *nup98a* mutants of Arabidopsis (Fig 7D, S27-S28). The hypocotyl length of 3-day-old Arabidopsis seedlings grown in darkness was measured for both untreated Control and Hormone-treated plants. The results showed that there was no difference between these three lines of seedlings under untreated control condition (Fig 7E). In contrast, there was a significant difference existing in between *Col-0* and *rae1* as well as in between *Col-0* and *nup98a* under the hormone-treated conditions (Fig 7F), suggesting that both modular components play a role in the hormone response of Arabidopsis.

## Discussion

### Selection of qXL-MS pipeline for study of crosslinks in nucleome

The qXL-MS workflow consists of multiple sequentially performing components (Liu et al., 2018b). Once a biological system is chosen for the study of special biological questions, the ultimate interactomic results are basically determined by a combination of the arbitrarily selected wet and dry laboratory methods, ranging from a choice of <u>C</u>hemical crosslinking (1^st^ C) for identification of XL-peptides to the chemical labeling either of proteins or peptides for quantitation, to the <u>C</u>hromatographic (2^nd^ C) separation and enrichment of XL-peptides for MS/MS analysis, and finally to the choice of software used for <u>C</u>omputational analysis (3^rd^ C) of MS data and quantitation (See Reviews by Chavez *et al*., 2016; Yu and Huang, 2018; Matzinger and Mechtler, 2021). Out of different biological systems studied so far, the subcellular organelles, such as mitochondria and nuclei, were often chosen to perform qXL-MS-based interactomic studies (Bartolec et al., 2020; Chavez et al., 2020; Fasci et al., 2018; Makepeace et al., 2020; Schweppe et al., 2017). In these qXL-MS experiments, various types of MS non-cleavable (*i.e*. BS^3^) and MS-cleavable (*i.e*. DSSO, CBDPS, BDP-NHP) crosslinkers have been applied to determine and quantitate the interactomes of organellar proteins (Fasci et al., 2018; Linden et al., 2020; Liu et al., 2018a). Given that formaldehyde is a general tissue fixation reagent that has been introduced into the qXL-MS research for some times (Liu et al., 2018b; Yu et al., 2019; Zhu et al., 2016a), the *in vivo* formaldehyde-crosslinking of living cells was purposely hybridized with the *in-organello* CBDPS-crosslinking of nuclei to generate a unique double crosslinking approach for qXL-MS application (Fig 1A). This double crosslinking method combined the extraordinary characteristics of formaldehyde both in rapid penetration of multiple cellular membrane systems and in fixation of *in vivo* architecture of protein-protein interactions in nucleome with the efficient crosslinking capability of CBDPS, which is a membrane permeable, tag enrichable, MS-cleavable, lysine site-specific and homo-trifunctional protein crosslinker (Makepeace et al., 2020). This combination was evidently successful in identifying a total of 10,169 unique crosslinks, containing 5,340 and 4,829 repeatable (PSM > 1) and non-repeatable (PSM = 1) XL-peptides (crosslinks), respectively (Table S1d). Throughout the bioinformatic and modulomic analysis, only the repeatable crosslinks were used.

The use of formaldehyde for *in vivo* fixation of the macromolecule interaction has also been manifested successfully in the proximity ligation of 3-C, Hi-C and SPRITE methods, where the macromolecule interactions, such as DNA to DNA or RNA to DNA interactions, were prefixed by formaldehyde in living cells and subsequently processed to elucidate chromatin-chromatin interactions and chromatin-RNA condensate interactions (Dekker et al., 2002; Nagano et al., 2013; Quinodoz et al., 2018). The reason why formaldehyde is so widely used as a fixative agent in study of interaction and subcellular localization of various macromolecules is due to its chemical reactive properties. It acts as reactive electrophilic species together with its derivatives to readily interact with various functional groups of cellular nucleic acids, polysaccharides and especially proteins to preserve the global organization and composition of cellular macromolecules in organelles, cells and tissues (Fox et al., 1985; Thavarajah et al., 2012). This aldehyde agent is also known to have a rapid and strong membrane penetration ability (5 to 10 min across plasma membrane, Fox *et al*., 1985; Toews *et al*., 2008), thus fixing the organization and composition of macromolecules including proteins in their actual physical positions, under liquid-liquid <u>p</u>hase <u>s</u>eparation (LLPS), at times of treatments or during nuclei isolation. CBDPS, on the other hand, has an arm-length of 11.4 Angstrom (measured by ChemDraw software, Makepeace *et al*., 2020), the size of which is small enough to flow, arguably, freely through the 52 - 107 Angstrom-diameter channel of nuclear pores (Schuller et al., 2021) to increase its access to nuclear proteins in nucleome. The only disadvantage of this double crosslinking is that the primary crosslinking site of both formaldehyde and CBDPS is the lysine residue on protein. We therefore deliberately reduced the formaldehyde concentration to 0.2% instead of using a routine concentration of 1% (Yu *et al*., 2019) in order to ameliorate the competition problem existed in between two crosslinkers. The extent of this competition was in fact evaluated by comparing the qXL-MS results from both the non-formaldehyde-prefixed and the formaldehyde-prefixed tissues. For examples, a total of 17,776 unique XL-peptides (crosslinks), containing 7,754 repeatable and 10,022 non-repeatable crosslinks, were identified from the CBDPS-crosslinked nuclear proteins (Table S1c). This number of unique crosslinks was 1.8-fold of that (10,169) of the formaldehyde-prefixed and CBDPS-crosslinked nuclear proteins, suggesting that the use of formaldehyde to prefix tissues could substantially reduce the discovery of nuclear PPIs 44%. Another measurement on the effect of formaldehyde in reducing the crosslink’s identification is the size of local interactomes of nuclear pore protein, NUA. This nuclear pore anchor protein was found to interact with 20 and 26 non-nuclear and nuclear proteins, respectively, under the direct *in-organello* CBDPS-crosslinking (Table S1h; Fig S29B). However, the number of these interactors substantially decreased to 8 and 14, respectively, in the *in vivo* formaldehyde-prefixed nuclei sample (Fig S29B). To circumvent this lysine site competition problem, it is, therefore, speculated that the switching of protein crosslinker CBDPS to another non-amine reactive crosslinker, such as DHSO targeting acidic amino acids (Gutierrez et al., 2016) or DAU targeting cysteine (Iacobucci et al., 2018), may help further increase the discovery number of unique crosslinks from the formaldehyde-prefixed nucleome in the future experiments.

Having said that, an alternative explanation on why formaldehyde reduced the discovery of crosslinks may result from the availability of the reactive amine moiety on a lysine residue of protein. In our previous study on Arabidopsis acetylproteomics (Liu et al., 2018c), the size of acetylproteome obtained from the formaldehyde-prefixed organism was found to be significantly larger than those discovered from the non-formaldehyde-crosslinked organism. It was speculated that the formaldehyde-prefixed living cells might contain a lower level of deacetylase activity so that a higher number of acetylated lysine residues might be maintained on organellar proteins during the nuclei isolation process. The extensive acetylation of nuclear proteins at lysine sites resulting from formaldehyde crosslinking of cellular components might prevent *in-organello* CBDPS crosslinking of nuclear proteins so that the identified crosslinks were relatively fewer. In contrast, if living cells were not subjected to the formaldehyde-prefixation, the broken cells would release a large number of deacetylases to remove acetylation from lysine residues, which would expose a lot of free amines to CBDPS crosslinker so that more CBDPS-crosslinked peptides would be identified.

As mentioned above, a choice of protein crosslinker used in various applications is often coupled with a corresponding XL-peptide identification search engine. The CBDPS was initially applied in XL-MS studies with a help of both light- and heavy-coded isoforms for XL-peptide identification (Makepeace et al., 2020). This pair of isotopically coded crosslinks were usually identified using Qualis-CL software. Since crosslinks were further dimethyl labelled at the N-termini for quantification purpose, this pair of isotope-coded crosslinkers would dramatically increase, in theory, the complexity of computational analysis. Thus, both Qualis-CL software and heavy isotope-coded CBDPS were not adopted in our experiment. Instead, we decided to integrate the use of less expensive light isotope-coded CBDPS with the dimethyl-labeling of peptides during qXL-MS analysis. To analyze the multiple chemical-labelled crosslinks, a common and widely used XlinkX search engine was therefore selected to perform the CBDPS-crosslinked peptide identification because this software, first introduced to the qXL-MS field by Heck’s group in 2015 (Liu et al., 2015), was able to analyze the cleavable DSSO-crosslinked peptides using the MS data collected from both CID/HCD and ETD detectors. The combinational use of CID/HCD and ETD detectors has been shown to enhance the identification of crosslinks (Liu et al., 2015). To make it suitable for identification of CBDPS-crosslinked peptides, a few modifications need to be made on XlinkX (See Star Methods for details). At the end, the integration of all these modifications into XlinkX made a modified software mXlinkX, the validation of which was conducted using both synthetic peptides and BSA proteins before it was fully applied in this qXL-MS pipeline (Fig S4; see Star Methods for details). The online link of the open-source scripts can be found in Key Resource Table.

### Considerations during the construction of modules and communities of protein interaction

Modulomics has been developed in the past years to aim at generation of the subgraphs linking with biological insights from a large dataset of PPIs, cell signaling cascades and gene regulatory networks. For an example, during the construction of the transcription-associated modules, a novel bioinformatics tool, MOCHI, was built to integrate the chromatin interaction and gene regulatory networks into identifying <u>h</u>eterogeneous interactome <u>m</u>odules (HIMs, Tian *et al*., 2020). Similarly, many other computational algorithms have also been proposed to perform datamining of these PPI databases in order to identify the protein interaction modules to gain the biological insights (Choobdar et al., 2019; Huttlin et al., 2017; Wu et al., 2020). Silverbush *et al*. generated a statistical framework to integrate multiple Omics data, PPIs, mutual exclusivity of mutations and copy number alterations, transcriptional coregulation as well as RNA co-expression to identify cancer diver pathways (Silverbush et al., 2019). In addition, MTGO integrated the PPIs and GO knowledge to identify topological and functional modules (Vella et al., 2018). Especially, Lin *et al*. utilized the PPIs of multiple species to identify the conserved proteins (called core components) from module variance. Both PPI evolution scores and interface evolution scores were introduced into this moduomic analysis (Lin et al., 2015). However, a large part of PPI datasets used by these modulomic approaches were retrieved from the public repositories of HIPPIE, BioGRID, InAct and STRING, which, however, lack the information about the abundance of PPI. In the present study, we selected the M1 method from the MONET toolbox as the module detection method. The M1 method is based on the optimization of a quality function, modularity, which is a relative quality measure for the partition of a network into modules and communities (Arenas et al., 2008; Newman, 2004). Unlike the traditional modularity optimization approaches, this algorithm utilizes a multiple resolution screening of the modular structure by searching the graph at multiple topological scales to overcome the resolution limit of modularity. Particularly, the algorithm introduced a resistance parameter *r* that prevents nodes from joining modules. The *r* = 0 is corresponding to the original scale, by which modularity was defined by Newman, while the positive and negative value of *r* reveals the substructures and superstructures of network, respectively. The M1 method will fit a resistance value *r* to the data to produce an optimized network for module detection. MONET itself is also unique in a sense that MONET toolbox (Tomasoni et al., 2020) was one of the three top-performing modulome construction programs, which was selected from the Disease Module Identification DREAM Challenge consortium (Choobdar et al., 2019). More importantly, this computational program has been demonstrated to be useful in recognizing disease-associated modules from a proteomic data collection (Johnson et al., 2022).

In the construction of modules and communities of nucleome graph, MONET utilized two pairs of variables, edge (protein-protein interaction, PPI) and the abundance of PPI as well as hybrid edge (module-module interaction, MMI) and the abundance of MMI, respectively. Since PPI information were construed from the information of crosslinks, a strong protein-protein interaction in nucleome is believed to result from more abundant crosslinks discovered. A question therefore raised is whether the abundance of nuclear proteins in nucleome is positively correlated to the abundance of PPIs. To address this question, we specifically performed a proteomic analysis on the crosslinked nuclei and identified 6,674 nuclear proteins after a filtration by plant nuclear protein repositories (Table S1j-k; Fig S29C). The abundances of these filtered nuclear proteins were determined using the Proteomic Ruler method (Wiśniewski et al., 2014). As a result, it was found that the abundances of these crosslinked nuclear proteins were distributed throughout the entire range of nuclear protein levels (Table S1j-k; Fig S29C). The median abundance of protein components of hetero-PPIs had no significant difference from that of the whole nuclear proteins (Table S1j-k; Fig S29D). However, the median abundance of proteins of homo-PPIs was 1.5 times higher than that of protein components of hetero-PPIs and that of nuclear proteins (Table S1j-k; Fig S29D), suggesting that crosslinking by CBDPS often occurred among subunits of abundant homodimers or oligomers of more stable interaction, leading to discovery of more abundant PPIs. As the abundance of PPI and the degree of the crosslinked protein components showed a lower correlation (0.25), it is also indictive that proteins of higher abundance were not necessarily highly interactive (Table S1j; Fig S29E). Further Examination of these crosslinks showed that the number and the type of crosslink was unrelated to the abundance of nuclear protein (Table S1j; Fig S29F-G). Taken together, the more abundant nucleomic proteins may not produce more abundant hetero-PPIs. The intensity of protein interaction may determine the abundance of PPI (or the abundance of a crosslink discovered). However, if an interaction strength is given between two components, the abundance of PPI will increase if two components or one of the two would increase in concentration under inductive conditions as demonstrated by the abundance of both hnRNPQ-H4 and RAE1-NUP98A interactions (Fig 7C).

When the abundance of PPI was included in modulomic analysis, this type of modulomics is considered to be weighted (Arenas et al., 2008; Liu et al., 2018a). If the unweighted modulomics is applied in the construction of nucleomic modulome, would it make a difference from what the weighted one may generate? To address this question, we arbitrarily reduced the abundance of all (1,211) Arabidopsis nuclear protein orthologous PPIs into a uniform integer one during the MONET computation process. It was quite interesting to find that the MONET-based modulomics actually captured 70 modules and 3 communities, which shared very little similarity with those modules and communities built using the same set of both PPI data and the quantitative information of these PPIs (Table S7a-d), suggesting that the unique feature, *i.e.* the abundance, of PPIs construed from crosslink data of qXL-MS, can make a difference in construction of the architecture of modulome. This finding also led us to speculate if the crosslinks of one-time-discovery (PSM = 1) acquired from qXL-MS would also alter the construction of modulome dramatically. To that end, we built modulomes using 1,878 and 3,274 Arabidopsis orthologous hetero-PPIs derived from dataset of PSM = 1 and dataset of PSM>=1 (Table S7a, S7e-j), respectively, each of which eventually generated 71 modules and 4 communities as well as 107 modules and 4 communities, respectively (Table S7a, S7e-j). None of these two modulomes exhibited Nucleolus Community. Moreover, these two modulomes showed very little similarity with that built using 1,211 PPIs derived from the repeatable crosslinks (PSM > 1; Table S7f, S7i). Taken together, it was concluded that the MONET-based modulome construction is sensitive to the abundance of PPI, and only repeatable crosslink data should be included in the modulomic analysis.

*In vivo* formaldehyde-prefixation of protein interaction has become a choice of interactomic practice for maintaining the native PPIs at or near native physiological levels (Liu et al., 2018b; Yu et al., 2019). To understand how the formaldehyde prefixation may influence the construction of modulome, we constructed 118 modules and 4 communities using the filtered nuclear 1,398 Arabidopsis orthologous hetero-PPIs (997 protein components) derived from crosslinks of non-formaldehyde-prefixed nuclei samples and compared them with that of formaldehyde-prefixed samples (Table S8a-g; Fig S30A-B). To our surprise, only 11% (10/95) NPIMs were common modules between these two datasets (Table S8c-e; Fig S30B). This percentage is much lower than that 41% (3,805/9,289) of common crosslinks found in between non-formaldehyde-prefixed and formaldehyde-prefixed samples (Fig S5B-C). Furthermore, GO enrichment analysis of the highly interactive proteins of 4 communities of both samples found Genome Community to be similar in between the two nuclei samples while the protein components of the rest three communities were totally different (Table S8f-g; Fig S30C-D). It is therefore evident that the formaldehyde-prefixation of living cells in the CHAMPION pipeline plays an essential role in capturing the real time on-going protein interactions in nucleome.

Having considered all these influential factors on construction of nucleomic modulome, the characteristic 4-Community architecture of protein interaction modulome might also be a stochastic product of MONET software even though the MONET-constructed modules and communities exhibited their preference for specific biological functions such as Genome Community and Nucleolus Community. To eliminate the possibility of randomness, we arbitrarily generated 1,000 Erdős–Rényi (ER; Erdös and Rényi, 1959) random modulomes using the equivalent number of nuclear proteins and hetero-PPIs used in this study (Fig S31A-B). As a result, none of the ER random-networking generated more than 40 modules, only about 6% of which produced four communities. These findings suggested that NPIMs and communities built from PPI data construed from crosslinks were not the consequences of random protein interactions (Fig S31C-D).

The PPI data generated by qXL-MS is apparently different those previously reported PPI data collected from Y2H, AP-MS, PCA, FRET and PDL approaches (Altmann et al., 2020; Kerppola, 2006; Qin et al., 2021; Rao et al., 2014b). A related question would be how much the source of PPI data can influence the construction of modulome. To address this concern, MONET-based modulomics was applied to analyze the Arabidopsis nuclear PPIs deposited in BioGRID (4,960 hetero-PPIs comprising of 2,067 nuclear protein components; Table S9a-d; Fig S32A). As a result, 156 modules and two communities were found, and there existed a large discrepancy in both the number of modules and communities between the two sources of PPI data (Table S9a-g; Fig S32B-D). No common community was identified based on the GO enrichment analysis between these two datasets. One explanation for such a great difference was that PPIs deposited in the BioGRID database were collected from different cell types, by different laboratories using various *in vivo* methods (such as heterologous yeast system) and under different conditions (Wu et al., 2020). An additional explanation may be that PPI data collected from molecular biological approaches cannot provide the abundance of PPI, which has been demonstrated to play an essential role in building the native protein interaction modulome of nucleome. In contrast, the *in vivo* qXL-MS protocol rather provided a uniform data real time data on both protein degree (degree of node) and abundance of PPI (intensity of edge) to MONET toolbox-based modulomic analysis even though these XL-MS collected PPI data used in this modulomic study in fact came from a mixture of multiple cell types in a tissue and the constructed modulome may represent a converged architecture of modular organization of nuclear proteins.

Finally, we question if the hierarchically assorted 4-Community of modular protein interactions in nucleome are universally present in the representatives of two kingdoms of organisms and if the 4-Community architecture of modular protein interactions in nucleome is both crosslinker- and software-independent. To address these questions, we performed the MONET-based modulomic analysis on the nuclear PPIs acquired from the human cell XL-MS, in which 3,312 PPIs comprising of 1,580 protein components of modules were reported by both Fasci *et al*., 2018 and Wheat *et al*., 2021 groups (see Star Methods for details). To our surprise, the modulomics, again, constructed 4 communities consisting of 150 modules (Table S10a-d; Fig S33A). Although no common module and community was observed between human and plant according to the JC comparison of modular components (Table S10a-e; Fig S33B-C). However, the GO enrichment analysis of highly interactive components in Genome Community of both datasets showed a high similarity (Table S10f-g; Fig S33D). These findings strongly suggested that the characteristic 4-Community of protein interaction modulome in nucleome, found by the qXL-MS with modulomics workflow, were neither CBDPS crosslinker-specific nor search engine-specific nor organism-dependent because those human cell PPI data were collected from different crosslinkers, DSSO and DSBSO, and XL-MS pipelines (Fasci *et al*., 2018; Wheat *et al*., 2021;Table S10).

### Implication of the CHAMPION pipeline in mapping the landscape of modular nucleome

The nucleomic proteins are known to participate in many nuclear events, ranging from chromatin organization, chromosome interactions, chromosome modification to DNA replication, regulation of gene expression and finally to the formation of membraneless sub-organellar compartments, nucleolus, nuclear bodies, biomolecular condensates and HUBs of DNA-RNA interactions (Strom and Brangwynne, 2019). Since the modular cell biology theory (Hartwell et al., 1999) predicted that these nucleomic proteins may interact with each other in a modular fashion to perform specific nuclear activities, the questions consequently raised are (1) how to systematically map the physical organization and distribution of nuclear proteins in nucleome and (2) how to associate modules of protein interaction with various nuclear events. Addressing these questions is certainly a challenging issue. Application of hormone treatment on plant cells further complicated the construction of 3-dimentional (3-D) nuclear protein interaction modulome to a 4-dimentaional (4-D) modulomics issue (Fig 4 and 5). To address these questions, the significantly regulated crosslinks (SRCs) and crosslinks, sequestered by qXL-MS pipeline that was performed on the hormone-treated cells, undoubtedly provided key information about the degree of nuclear protein components, the abundance of PPIs and Hormone-specific PPIs necessary for building the dynamic 4-D modulomes (Fig 5). Construing the SRCs into Hormone-specific (or called case-specific) PPIs allowed the MONET-based modulomics to capture the novel module and community variants specific to hormone treatment and permitted us to monitor the dynamic changing behaviors of modules and communities (Fig 5) of nucleomic proteins. We thus named the integrated workflow of both qXL-MS-based interactomics and MONET-based modulomics specially as CHAMPION pipeline.

Indeed, the CHAMPION pipeline has unexpectedly identified a characteristic 4-Community modulome, consisting of 55 nuclear protein interaction modules (NPIMs), from nucleome of plant cell (Fig 4). Those 250 novel interactors of core histone octamer and 26 novel interactors of BoxC/D snoRNP, identified from this qXL-MS (Fig 2 and 3), are believed to contribute significantly to the formation and organization of Genome and Nucleolus Community, respectively. The successful segregation of protein interactions present in the molecules-crowded (Bancaud et al., 2009) and phases-separated (Strom and Brangwynne, 2019) nucleome into discerned protein interaction communities by CHHAMPION pipeline demonstrated that the physical positions of hundreds and thousands of nucleomic proteins, one day other macromolecules (such as DNA and RNA), can be mapped once into discerned nucleoplasm territories and spaces in nuclei though a proteomic experiment. This type of proteomics-based and high-throughput PPI measurement technology should be complementary to the microcopy-based and slow-throughput biomolecular fragment complementation (BiFC) technique (Kerppola, 2006), by which PPIs are routinely investigated in cells (Yu and Huang, 2018). In addition, we hereby hypothesize that what the shortest path of two nuclear proteins positioned by modules and communities to the nucleome graph is what the centimorgan unit between two genes separated on a chromosome to the genetic distance. The graph index used here to define the position of a protein in nucleome may be applied, one day, to calculate the shortest path of two protein nodes in nucleus.

Another important finding of the CHAMPION workflow is the mapping of 27 NPIMs (out of 55 community-forming modules, 49%) containing modular components involved in forming biomolecular condensates (Table S4c), and, specially, is the finding of Super-Enhancer (SE) condensates from nucleome (Ma et al., 2021; Sabari et al., 2018). These nuclear bodies usually have dynamic behaviours, and contain highly dynamic components although they usually appear as overall stable structures (Phair and Misteli, 2000). As the composition of biomolecular condensates are difficult to be analysed using traditional biochemistry methods due to the difficulties in isolation of condensates (Ditlev et al., 2018). Because both formaldehyde and CBDPS are capable of penetrating through multiple membranes into condensates (Sabari et al., 2018; Yang et al., 2020), this CHAMPION pipeline was presumably able to capture many putative new components of various known condensates (Fig 4 and 5). For an example, in the Genome Community (C1), NPIM1-9 contained a transcription factor MHGB1 of a large Intrinsically Disordered Region (IDR) functioning as a contact domain for condensate-forming (Murray et al., 2017; Nott et al., 2015). At the same time, this module captured other known transcription factors like RR2, ASIL2, CAMTA4 and SWRC4 (Table S4c), suggesting a putative super-enhancer condensate probably forming in cells to regulate gene expression. Similarly, NPIM1-18 might capture a GDAC condensate with GBPL3 as scaffold protein and new modular components like SHP2, HB2, UTL2 and THO5 interactors as putative client proteins (Table S4c; Fig S13). In Nucleolus Community (C2), NPIM2-7 contains a well-known scaffold condensate-forming FIB protein, which is a core member of box C/D snoRNP and it has a N-terminal GAR domain for LLPS (Feric *et al*., 2016; Emenecker *et al*., 2020). Again, this module seemed to have captured RNA helicase subunits, rRNA processing proteins, gene silencing proteins and other RNA binding proteins as client proteins from a putative condensate participating RNA processing activities in nucleoli. Since the qXL-MS might have captured crosslinks occurred in different types of cells, the current NPIM may be a combinatorial module converged from a mixture of heterogenous module variants of many cell types or from the heterogeneity of genome organization in individual cells (Finn and Misteli, 2019) or from the inherently stochastic transcriptional bursting occurred in nuclear spaces (Hager et al., 2009; Misteli, 2001). It is therefore postulated that a temporal- and spatial-specific NPIM may have a smaller size of module of less number of modular components in a specific type of cell. Integration the qXL-MS with single cell transcriptomics (Longo et al., 2021) performed on the same specimens under the same conditions may help resolve this problem. Especially, the combination of SPRITE (Quinodoz et al., 2018) and ChIP-seq (Park, 2009) with the CHAMPION pipeline will generate a much higher resolution landscape on the hierarchically and spatiotemporally organized 4-Dimensional architectures of RNA, DNA and protein molecules in nucleomes of an organism upon treatments and at various developmental stages.

## Acknowledgments

This work was supported by grants: 31370315, 31570187, 31870231, 32070205 from the National Science Foundation of China, 16102422, 16103621, 16101114, 16103817, 16103615, 16100318, 16101819, 16101920, AOE/M-403-16, R4012-18, C6021-19EF, and GRF 16306919 from the RGC of Hong Kong, ITS/480/18FP and MHP/033/20 from the Innovation and Technology Commission (ITC) of the Hong Kong S.A.R. government, Hetao Shenzhen-Hong Kong Science and Technology Innovation Cooperation Zone project (HZQB-KCZYB-2020083) and the internal fund supports from HKUST.

## Author contributions

SJD conducted most of experiments. NL, SCL, WZ, TLW, AB and HD contributed to the establishment of MS conditions for analysis of XL-peptides and XL-MS protocol. WCY, CZ and FCY contributed to the computational analysis of crosslinks. NL, TLW, WCY, AB and SJD contributed to MS data analysis. NL and DSJ analysed data and wrote the manuscript. NL designed research. All authors have read and agreed to the published version of the manuscript.

## Notes

The mass spectrometry data have been deposited to the ProteomeXchange Consortium via PRIDE the with a data set identifier PXD034796.

Username

Password: (It will be public-available after acceptance of paper)

## STAR Methods

### Lead Contact and Materials Availability

Further information and requests for resources and reagents should be directed to and will be fulfilled by the Lead Contact, Prof. Ning Li.

### Experimental model and SUBJECT details

*Glycine max L. cv* Zhonghuang #39 (Soybean)

### Method Details

Sources of chemicals are indicated in the Key resources table.

### Plant Material and Growth Condition

The seeds of an experimental plant (*Glycine max L.* cv Zhonghuang#39) were sterilized in a solution of 75% ethanol (v/v) and 3% hydrogen peroxide (v/v) for 2 min with several gentle inversions. After decanting the sterilizing mixture, plant seeds were rinsed with double-distilled water for 5 times. Subsequently, the sterilized seeds were immersed in double-distilled water at room temperature for 12 hr in darkness. After that, a hundred of seeds were placed on a moisture 4-layer gauze inside an air-tight container (Length × Width × Height, 288 mm × 230 mm × 184 mm). All these experimental seeds were further divided into two batches. One batch of seeds geminated in containers filled with air as control seedlings while the other batch of seeds in containers fumigated with a plant hormone ethylene of 5 ppm. The concentration of ethylene in container was periodically measured by gas chromatography (model Perkin Elmer Clarus 580, PerkinElmer, Shelton, CT, USA). Both air-grown and hormone-treated seeds germinated in the plastic containers covered with aluminum foil to prevent light, which were incubated in a greenhouse at 23 +/- 1 °C for 7 days. In total, 6 pairs of air-grown and hormone-treated etiolated seedlings were harvested.

### Prefixing of Seedling Tissue with Formaldehyde (1^st^ crosslinking)

Three replicates of etiolated hypocotyls of both the untreated (Control) and the hormone-treated (Hormone) seedlings were harvested, immediately sliced into 1.5-mm thick discs and subsequently immersed in a buffer containing 10 mM sodium phosphate pH7.0, 0.4 M sucrose, 1 mM ethylenediaminetetraacetic acid (EDTA), 1 mM phenylmethylsulfonyl fluoride (PMSF), 0.1% formaldehyde, 0.1% para-formaldehyde (Keene *et al*., 2006; Liu *et al*., 2018a). The formaldehyde was infiltered into plant cells through 5 rounds of pressure-shifting process between 600 Torr (5 min) and atmospheric pressure (1 min) within 30 min to *in vivo* fix macromolecules in plant cells. The formaldehyde-prefixed tissues were rinsed for 3 times with double-distilled water and harvested in liquid nitrogen (Table S0a; Fig S1). The frozen tissues were stored at −80 °C freezer for nuclei isolation.

In contrast, another three replicates of etiolated hypocotyls of both the untreated (Control) and the hormone-treated (Hormone) seedlings were harvested without being dissected and immediately stored at −80 °C freezer for nuclei isolation (Table S0a; Fig S1).

### Nuclei isolation and CBDPS-crosslinking (2^nd^ crosslinking)

A hundred of grams each of frozen tissues of Control and Hormone seedlings (either with formaldehyde-prefixed or without) were ground into fine powders in liquid nitrogen, which were subsequently homogenized in 3 volumes (v/w) of Cell Lysis Buffer containing 20 mM 4-(2-hydroxyethyl)-1-piperazineethanesulfonic acid (HEPES, pH 7.4), 25% glycerol, 20 mM KCl, 2 mM EDTA, 250 mM sucrose, 1 mM PMSF and 1×Complete EDTA free protease inhibitors cocktail. The nuclei homogenate was filtered sequentially through 120 μm and 38 μm nylon meshes. The nuclei-containing filtrates were collected (Kaufmann et al., 2010; Saleh et al., 2008) and centrifugated at 2000 xg for 10 min (Cheng et al., 2009).

The nuclei pellets were rinsed with 3 volumes (v/w) of Crosslinking Buffer, which contained 20 mM HEPES pH 8.0, 25% glycerol, 2.5 mM MgCl_2_, 250 mM sucrose, 5% Dimethyl sulfoxide (DMSO) (Carrier et al., 2011). The 1.2 grams of nuclei samples were crosslinked again with an amine-reactive crosslinker <u>c</u>yanur <u>b</u>iotin <u>d</u>i<u>p</u>ropionyl <u>s</u>uccinimide (CBDPS; arm length measured by ChemDraw: 11.4 Å; molecular weight: 739.1512) obtained from Creative Molecules Inc (Petrotchenko et al., 2011). The crosslinker was freshly prepared in DMSO to achieve a stock solution of 50 mM, which was added subsequently into nuclei at a concentration of 0.5 μM with 6 rounds (3 min / round) with rotation. As a result, it was estimated that the amount of CBDPS crosslinker added to the nuclei sample should reach a final concentration of 3 mM. The nuclei samples were crosslinked for 30 min at room temperature, followed by a centrifugation at 2000 g for 10 min. Immunoblot and fluorescence microscopy were used to measure the efficiency of nuclei isolation and CBDPS crosslinking as well as the integrity of nuclei (Fig S2).

The nuclei pellet was re-suspended in 5 volumes (v/w) of Protein Resuspension Buffer 1 containing 150 mM Tris-HCl pH 8.0, 20 mM EDTA, 20 mM ethylene glycol-bis(β-aminoethyl ether)-N,N,N′,N′-tetraacetic acid (EGTA), 50 mM NaF, 1% glycerol-2-phosphate, 1mM PMSF, 8 M urea, 2% glycerol, 1.2% TritonX-100 and 0.5% sodium dodecyl sulfate (SDS), 5 mM ascorbic acid, 5 mM dithiothreitol (DTT) to lyse the organelles (Li et al., 2009). Nuclei were disrupted by 10-sec pulses with a sonicating probe at 50% intensity for 10 min. Three volumes (v/v) of pre-cooled 12:1 (v/v) acetone/methanol solution was applied to the mixture for at least 4 hr at −20 °C to precipitate proteins. The protein pellet was subsequently air-dried and resuspended in 5 volumes (v/w) of Protein Resuspended Buffer 2 containing 50 mM Tris-HCl pH 8.0, 50 mM NaF, 1% glycerol-2-phosphate, 1mM PMSF, 8 M urea, 2% glycerol, 2% TritonX-100 (Li et al., 2009), followed by applying 10 mM DTT for 30 min, 40 mM IAM (protected from light) for 30 min and 10 mM DTT for 10min at room temperature. The solution was subsequently mixed with 3 volumes (v/v) pre-cooled 12:1 (v/v) acetone/ methanol solution for at least 4 hr at −20 °C (Table S0a; Fig S1). DC protein assay (Bio-Rad, Hercules, CA, USA) was performed to quantify the protein concentration using a Bovine Serum Albumin (BSA) standard curve.

### In-solution Digestion of Proteins and Dimethyl Labeling of the Crosslinked Peptides

Protein samples were air-dried and dissolved into Protein Re-suspension Buffer 3 containing 40 mM Tris-HCl, pH 8.0 and 6 M urea. The protein solution was diluted using a pre-heated (37 °C) trypsin digestion buffer (40 mM Tris-HCl, pH 8.0) to ensure the final concentration of urea was lower than 1 M. The protein samples were digested for two rounds. In the first round, the crosslinked protein samples were digested with trypsin protease at an enzyme-to-protein ratio of 1:50 (w/w) at 37 °C for 16 hours, whereas, in the second round, protein samples at an enzyme-to-protein ratio of 1:100 (w/w) at 37°C for 8 hr. The crosslinked peptides (XL-peptides) were desalted using C18 Sep-Pak cartridges (Waters, Manchester, UK) and concentrated using SpeedVac (Thermo Scientific Inc., Waltham, MA, USA). The XL-peptides derived from either Control or Hormone protein sample was dissolved in 100 mM sodium acetate (pH 5.5), each of which was divided into two parts equally and separately labeled with light (L) and heavy (H) isotope-coded formaldehyde chemicals, respectively. The light (L) isotope-labeled nuclear peptides of Control sample and the heavy (H) isotope-labeled nuclear peptides of Hormone sample were mixed equally as the forward experiment (F). The L isotope-labeled nuclear peptides from Hormone sample and the heavy isotope-labeled nuclear peptides from Control sample were mixed equally as the reciprocal experiment (R). The F and R mixtures were treated as two independent experimental replicates from a single biological replicate. The nuclear peptides were further desalted using the C18 Sep-Pak cartridges, followed by nuclear peptide concentration and acetonitrile (ACN) removal with SpeedVac (Table S0a; Fig S1). The nuclear peptides were stored at −80°C freezer before further use. In the end, a total of 6 biological replicates were performed, consequently generating 12 experimental replicates of nuclear peptide mixtures, among which 6 experimental replicates of nuclear peptides were generated from 3 biological replicates of double-crosslinked nuclei while the other 6 nuclear peptide samples from three replicates of CBDPS-crosslinked nuclei (Table S0a; Fig S1).

### Strong Cation Exchange-based High-performance Liquid Chromatography (SCX-HPLC)

The chromatographic separation was performed using a high-performance liquid chromatography (HPLC) system (Waters, Manchester, UK) coupled with an SCX column (200 × 9.4 mm, PolySULFOETHYL ATM, 5 µm, 200 Å, 209SE0502, PolyLCINC, Columbia, MD, USA). Both Buffer A (7 mM KH_2_PO_4_, 30% ACN, pH 3) and Buffer B (7 mM KH_2_PO_4_, 30% ACN, 350mM KCl, pH 3) were used as the mobile phases. The separation gradient was set as the following: Buffer A 100% to 90% in 10 min, 90% to 83% in 22 min, 83% to 68% in 8 min, 68% to 20% in 10 min, 20% to 0% in 2 min, 0% to 100% in 12 min and 100% for 30 min. For each biological replicate, 108 fractions (54 × F fractions and 54 × R fractions) were collected. The nuclear peptides were further desalted using the Oasis Hydrophilic-Lipophilic-Balanced (HLB) 1 cc cartridge (Waters, Manchester, UK), followed by peptide concentration and ACN removal with SpeedVac. The nuclear peptide samples were stored at −80 °C before further use.

### Affinity Enrichment of XL-peptides

The nuclear peptide samples were resuspended using 50 mM HEPES (pH 7.5) and subsequently incubated with the high-capacity streptavidin agarose resin (Pierce, Rockford, IL, USA) for 2 hr at room temperature. Washing was performed with a washing buffer consisting of 50 mM HEPES (pH 7.5). Afterwards, beads were incubated with 70% ACN and 0.5% FA for 1 hr to elute XL-peptides. The elution was performed twice to achieve a higher recovery rate. The XL-peptide samples were thereafter desalted using the Ziptip (MilliporeSigma, Burlington, MA, USA), followed by XL-peptide concentration and ACN removal with SpeedVac (Table S0a; Fig S1). In total, there were about 40 injections per experimental replicate (499 injections in total). The samples were stored at −80 °C before further use.

### Generation of Proteome Database of the Crosslinked Nuclear Proteins

The flow-through of avidin affinity column enrichment was collected and subsequently combined into 5 fractions from each experimental replicate. The nuclear peptide samples were further desalted using the Ziptip, followed by peptide concentration and ACN removal using SpeedVac. The air-dried peptide samples were resuspended into 100 µL of 0.1% formic acid and separated by a 120 min gradient elution at a flow rate of 0.3 µL/min with a Thermo-Dionex Ultimate 3000 HPLC system., which interfaced with a Thermo Orbitrap Fusion Lumos mass spectrometer. The analytical column was the Acclaim PepMapTM RSLC C18 Capillary column (75 µm ID, 150 mm length; Thermo Scientific Inc., Waltham, MA, USA). Mobile phase A consisted of 0.1% formic acid, and mobile phase B consisted of 100% acetonitrile and 0.1% formic acid. All spectral data were acquired in the Orbitrap mass analyzer. For the MS scans, the scan range was set to 300 - 2000 m/z at a resolution of 120 K, the automatic gain control target was set to 500,000 and the charge state was set as +2 - +8. For the MS/MS scans, the resolution was set to 30,000, the automatic gain control target was set to 300,000, the precursor isolation width was 1.6 Da, and the maximum injection time was set to 54 ms. The HCD normalized collision energy was 26%.

Raw mass spectrum data acquired was searched against the soybean protein database deposited in the NCBI (downloaded on 01/11/2018, containing 71,730 sequences) using Comet (version 2019.01 rev2). The default settings of the Comet were used unless stated otherwise. The decoy-searching mode was set to 1 (concatenated search). Carbamidomethylated cysteine was specified as a static modification, while methionine oxidation, lysine, and peptide N-term light (28.0313 Da) and heavy (34.0631 Da) isotope-coded dimethyl were included as variable modifications. The output format percolator file was enabled. FDR of the resulting Peptide Spectrum Matches (PSMs) were estimated using Percolator (version 3.02), and the q-value threshold was set as 0.01.

### LC-MS/MS Analysis of XL-peptides

XL-peptides were separated by a 120 min gradient elution at a flow rate of 0.3 µL/min with a Thermo-Dionex Ultimate 3000 HPLC system, which interfaced with a Thermo Orbitrap Fusion Lumos mass spectrometer. The analytical column was the Acclaim PepMapTM RSLC C18 Capillary column (75 µm ID, 150 mm length; Thermo Scientific Inc., Waltham, MA, USA). Mobile phase A consisted of 0.1% formic acid, and mobile phase B consisted of 100% ACN and 0.1% formic acid.

In the CID-MS/MS experiment, we selected the 10 most abundant precursors and subjected them to a sequential CID-MS/MS and ETD-MS/MS acquisition protocol. All spectral data were acquired from the Orbitrap mass analyzer. As to MS scans, the scan range was set to 300 – 2,000 m/z at a resolution of 120 K, the automatic gain control target was set to 500,000 and the charge state was set as +4 to +8. For the MS/MS scans, the resolution was set to 30,000, the automatic gain control target was set to 300,000, the precursor isolation width was 1.6 Da and the maximum injection time was set to 54 ms. At the same time, the CID normalized collision energy was 30%, the charge-dependent Electron Transfer Dissociation (ETD) reaction time was enabled, the orbitrap resolution was 30 K, the ETD automatic gain control target was set to 300,000, the scan range mode was auto normal, the precursor isolation width was 1.6 Da and the maximum injection time was set to 54 ms. For the HCD-MS/MS experiment, the settings were the same as the CID-MS/MS except the MS/MS activation type was set as HCD and the collision energy was 26% (Table S0; Fig S1).

### In-house Modification of XlinkX Scripts and Validation with CBDPS-crosslinked Peptides and Protein

The scripts of XlinkX (Liu et al., 2015, 2017) were downloaded from http://sourceforge.net/projects/xlinkx/. Five major modifications were made on XlinkX. (1) The scripts were firstly modified into a command-line version with parallel computing (using Parallel and doSNOW packages of R) to run on the High-Performance Clusters (HPC) with multiple cores (see Key resource table for details). The structures of data in the scripts were adjusted to save the Random-Access Memory (RAM) in computation. (2) The requirement for signature fragment ions was reduced from 4 to 3 during identification of XL-peptide as the fragmented peptide ions of the longer arm, harboring biotin tag, cleaved from the fragmented XL-peptide were not easy to be detected. (3) The parameters of CBDPS were added to the software, which included the mass of the shorter (54.0106 Da) and the longer (455.0878 Da) arms. (4) A derivative of CBDPS, containing biotin sulfoxide moiety, was integrated into XlinkX in searching for XL-peptides (the mass of the shorter and the longer arm was set as 54.0106 Da and 471.0820 Da, respectively), as it was previously reported that a crosslinker of biotin moiety could be derivatized into an oxidized form (called biotin sulfoxide moiety) either during the experimental procedures or MS analysis (Kang et al., 2009). All three organic compounds, biotin, biotin sulfoxide and CBDPS were subjected to LC-MS/MS analysis (Fig S3). As a result, no oxidized biotin derivative, biotin sulfoxide, was detected from both biotin and CBDPS, suggesting the oxidation of biotin occurred during the procedure of XL-peptide preparation from cells or under *in vivo* conditions. Moreover, the most insensitive fragment ion in MS/MS spectra of biotin (m/z 227.0847, 1+) and biotin sulfoxide (m/z 243.0797, 1+) fitted well with the biotin and biotin sulfoxide moiety of CBDPS measured from XL-peptides of this study (Fig S3), which further supported that the derivative of CBDPS measured from XL-peptides of this qXL-MS study was a biotin-sulfoxide moiety-containing CBDPS. (5) Finally, we increased the number of variable modifications from 1 to 4 to enable the searching of both light (28.0313 Da) and heavy isotope-coded dimethyl (34.0631 Da) because they were conjugated to both lysine residues and N-termini of XL-peptides during the initial step of XL-peptides preparation. In addition to these major modifications, we also modified the read_mgf function to use the peak lists from the output of Hardklor (Hoopmann et al., 2007, 2012). The functions for generating and comparing fragment ions were modified to be compatible with adding the number of variable modifications. Finally, an additional file for percolator FDR estimation was generated as previously described (Liu et al., 2017). As result, the new mXlinkX software only searched for inter- and intra-crosslinks instead of the mono-crosslinks including loop-crosslinks and single-crosslinks.

To access the reliability of the newly modified XlinkX (mXlinkX), the synthetic peptides, Fmoc-KELDDLR and Fmoc-EAKELIEGLPR, were crosslinked with CBDPS. These two peptides were synthesized by Minghao Biotechnology (Wuhan, Hubei, China). Fmoc-protected N-terminal amine was used to promote the crosslinking between two peptides at the side chains of lysine. The CBDPS-based crosslinking was performed using a previously described method (Liu et al., 2018b; Zhu et al., 2016b). The XL-peptides were labeled separately with light and heavy CH_2_O using the method described above, followed by desalting with Oasis HLB cartridges and concentrating with SpeedVac. Consequently, the chromatographic enrichment was performed with the HPLC system coupled with a SCX column. Both Buffer A (7 mM KH_2_PO_4_, 30% ACN, pH 3) and Buffer B (7 mM KH_2_PO_4_, 30% ACN, 350mM KCl, pH 3) were used as the mobile phases. The separation gradient was set as the following: Buffer A 100% to 90% in 10 min, 90% to 83% in 22 min, 83% to 68% in 8 min, 68% to 20% in 10 min, 20% to 0% in 2 min, 0% to 100% in 12 min and 100% for 30 min. The enriched peptides were collected and further desalted again using Oasis HLB cartridges. Subsequently, the streptavidin beads-based affinity enrichment (described above) was performed. The eluted peptides were enriched with Ziptip. Finally, the XL-peptides were analyzed using LC-MS/MS (as described in **LC-MS/MS Analysis of XL-peptides** section) and the peptide searching of the spectrum data was performed by the mXlinkX (Table S0b; Fig S4A-B). As a result, we obtained the MS/MS spectra of synthetic peptides, KELDDLR (⍰) - EAKELIEGLPR (β), that were crosslinked either with CBDPS of biotin moiety or CBDPS of biotin sulfoxide moiety (Fig S4 A-B). For peptides crosslinked with CBDPS of biotin moiety (Fig S4A), we observed a △m of 401.0762 Da between the longer (X_L_) and shorter arm (X_S_) of the cleaved crosslinker, and the four signature fragment ions identified were ⍰ + X_S_ (969.5130 Da, theoretical 969.5131 Da), β + X_L_ (1736.8161 Da, theoretical 1736.8170 Da), ⍰ + X_L_ (1370.5795 Da, theoretical 1370.5903 Da) and β + X_S_ (1335.7410 Da, theoretical 1335.7398 Da). On the other hand, the △m observed for CBDPS of biotin sulfoxide moiety was 417.0711 Da and the three fragment ions for the crosslinked synthetic peptides were ⍰ + X_S_ (969.5135 Da, theoretical 969.5131 Da), ⍰ + X_L_ (1386.5848 Da, theoretical 1386.5845 Da) and β + X_S_ (1335.7423 Da, theoretical 1335.7398 Da).

The ground truth of protein level was built by crosslinking of BSA with CBDPS according to the method described above (Table S0c; Fig S4C-D). Similarly, the crosslinked peptides either with CBDPS of biotin moiety or biotin-sulfoxide moiety were observed in the CBDPS-crosslinked BSA protein. For example, it was found that the BSA peptides, DTHKSEIAHR (⍰) and FKDLGEEHFK (β), crosslinked either with CBDPS of biotin moiety (Fig S4C) or biotin sulfoxide moiety (Fig S4D). For the MS/MS spectra of CBDPS of biotin moiety-crosslinked peptides, four signature fragment ions were found, which were ⍰ + X_S_ (1274.6219 Da, theoretical 1274.6367 Da), β + X_L_ (1759.7531 Da, theoretical 1759.7642 Da), ⍰ + X_L_(1675.7250 Da, theoretical 1675.7139 Da) and β + X_S_ (1358.6810 Da, theoretical 1358.6870 Da). Likewise, signature fragment ions ⍰ + X_S_ (1274.6580 Da, theoretical 1274.6367 Da), β + X_L_ (1775.7408 Da, theoretical 1775.7584 Da), ⍰ + X_L_ (1691.7121 Da, theoretical 1691.7081 Da) and β + X_S_ (1358.6882 Da, theoretical 1358.6870 Da) were observed for peptides crosslinked with CBDPS of biotin sulfoxide moiety. Moreover, the crosslinks of the BSA were mapped to the crystal structure of BSA dimer (PDB: 3V03). The distances between two alpha carbons (Cα) of the crosslinked lysine residues on BSA dimer were calculated (Fig S4E-F). The different distribution of the Cα-Cα distances between the crosslinked and randomly assigned lysine residues demonstrated a successful identification of crosslinks on BSA with the help of mXlinkX (Fig S4F). Moreover, the evaluation of mXlinkX was also performed on the CBDPS-crosslinked mitochondria MS data (Makepeace et al., 2020) with a similar MS/MS spectra observation with crosslinked synthetic peptides and BSA proteins (Fig 4 G-H), which further supported the reliability of mXlinkX in search for CBDPS-crosslinked peptides.

### Data Analysis and Processing

The raw MS data were converted and deconvoluted with Hardklor (version 2.3) (Hoopmann et al., 2007, 2012). The soybean plant protein database was constructed with both the previously published and the in-house analyzed proteome data. The mXlinkX was used for XL-peptide identification. For the *in-silico* protein digestion, the variable modifications were set as oxidation on “M”, and both light and heavy isotope-coded dimethyl (28.0313 Da and 34.0631 Da, respectively) were set on both “K” and the N-terminus of a peptide. The Carbamidomethyl on “C” was set as the fixed modification. The enzyme was set as trypsin. As for the crosslinking searching, the tolerance for MS and MS/MS searching were 10 ppm and 20 ppm, respectively. The short-chain monoisotopic mass of the crosslinker was set as 54.0106 Da while the heavy chain was set as 455.0878 Da and 471.0820 Da; The Precursor selection tolerance was 0.05 Da. The fragment types were set as “CID/HCD” and “ETD” respectively. The false discovery rates of the output were calculated by the Percolator (version 3.02) (Käll et al., 2007; The et al., 2016) and only the PSMs with q-value ≤ 0.01 was collected.

The subcellular localization of crosslinked proteins was obtained according to both the established soybean nuclear proteome (comprising of 7,378 soybean nuclear proteins) (Cooper et al., 2011; Yin and Komatsu, 2015) and Arabidopsis nuclear proteome (consisting of 4,246 Arabidopsis nuclear proteins) (Hooper et al., 2017; Mair et al., 2019). The crosslinked proteins that overlapped with anyone protein identification of both documented nuclear proteomes was defined as nuclear proteins. The nuclear XL-peptides or crosslinks were converted into nuclear Protein-Protein Interactions (PPIs). The nuclear PPIs were further divided into nuclear hetero- and homo-PPIs. The homo-PPI means that the crosslinked proteins on both ends of the CBDPS crosslinker are identical in polypeptide sequence, whereas hetero-PPI means that the two nuclear proteins crosslinked by CBDPS have discrete amino acid sequences. The abundance of crosslinked nuclear proteins was measured using the nuclear proteome data by the Proteomic Ruler (Wiśniewski et al., 2014). The hetero-PPIs were analyzed by Cytoscape (version 3.7.2) as an undirected network. The heatmap matrix of PPIs was conducted and visualized by the R package Pheatmap. PPIs documented from the literature were collected from two public PPI databases: STRING (Szklarczyk et al., 2019) and BioGRID (Oughtred et al., 2021). Only the interactions with a combined score of ≥ 700 were considered in the STRING database.

### Gene Ontology Analysis

The soybean proteins were converted into Arabidopsis ortholog by NCBI blastp (the hit with the lowest e-value). The Arabidopsis orthologs were used for the gene ontology analysis which was performed using the g:Profiler (Raudvere et al., 2019) with the default settings. The results of the GO enrichment analysis were visualized using the R package pheamap and ComplexHeatmap.

### Quantification of XL-peptides

The extracted ion chromatogram (XIC)-based quantification was conducted using SQUA-X software (Liu et al., 2018b, 2018c). The following criteria were used for selection of quantifiable XL-peptides: **1.** the number of PSMs of light dimethyl labeled XL-peptides ≥ 1; **2.** the number of PSMs of heavy dimethyl labeled XL-peptides ≥ 1; **3.** the number of experimental replicates ≥ 4; **4.** the number of identified PSMs from the forward/reciprocal experiments divided by the total number of PSMs ≥ 0.2. After the quantifiable XL-peptides were selected, the next procedure was to pair the ion chromatograms of light- and heavy-labelled XL-peptides. For the situation that both two differentially labelled XL-peptides were identified, their corresponding ion chromatograms were then calculated and paired directly. However, if only one of the light- or heavy-labelled XL-peptide was identified, the SQUA-X would calculate a theoretical m/z value and an isotopic pattern of the missed XL-peptide. After that, SUQA-X would find the ion chromatogram of the missed XL-peptide with the following criteria: (1) the observed m/z value should be equal to the theoretical m/z value with a mass tolerance of 0.05 Da; (2) the Pearson correlation of the observed isotopic pattern and theoretical isotopic pattern should ≥ 0.70; and (3) there should be an overlap between two retention time ranges of the light- and heavy-labeled ion chromatograms. Notably, SQUA-X used the half value of the minimum intensities among all extracted ion chromatograms to replace the zero intensity. Subsequently, the logarithm ratio of the light and heavy isotope-labeled crosslinks was calculated using the maximum intensities of the chromatograms (Cox and Mann, 2008). All logarithm ratios were then adjusted using the median value of the total population of logarithm ratios from the same replicate to circumvent mixing error during experiment, following by batch effect adjustment, one-sample student *t*-test and multiple hypothesis test correction using Benjamini-Hochberg procedure (Benjamini and Hochberg, 1995; Liu et al., 2018b, 2018c). Eventually, all the quantifiable XL-peptides were resulted with a mean value of corresponding logarithm ratios and a q-value by Benjamini-Hochberg procedure. The significantly regulated XL-peptides were selected with q-value ≤ 0.1 and the absolute value of log-ratio ≤ 0.5 × Standard Deviation (SD).

### Structural Modeling

The 3D protein structure prediction was performed using the homology modeling tool SWISS-MODEL. The structures of the histone octamer and DNA double helix were built through the SWISS-MODEL using the X-ray structural data of the chicken nucleosome particles (PDB: 1EQZ). The predicted structure with the highest sequence coverage was selected for analysis and display. The missing tail regions of histone octamer were modeled with the sequence alignment of the predicted histone octamer and individual core histone proteins (histone H2A, H2B, H3 and H4) by PyMOL. The structure of the box C/D Small Nucleolar Ribonucleoprotein (snoRNP) complex was built through the SWISS-MODEL using the Cryo-EM structural data of the 90S pre-ribosome from *Chaetomium thermophilum* (PDB: 5OQL). Both the surface and ribbon representatives of the 3D structures were generated with PyMOL version 2.30 (Schrödinger, LLC). The measurement of Cα-Cα distance was accomplished by PyMOL. Random Cα-Cα distances were calculated by all the possible lysine-lysine residue pairs of the corresponding protein structures. All the 2D crosslinking maps were generated with xiView (Graham et al., 2019).

### Terminology and Construction of the Hierarchically Assorted Modules of Protein Interaction

The NCBI blastp was firstly used to convert the soybean plant nuclear PPIs into Arabidopsis ortholog PPIs. The Arabidopsis ortholog PPIs were subsequently processed using R to fit the input format of the MONET toolbox (Arenas et al., 2008; Tomasoni et al., 2020). According to the graph theory (Al-Taie and Kadry, 2017), the two interacting proteins and the connection between the two proteins of a PPI were defined as nodes and an edge (or called a link), respectively. The number of connections on a protein node was defined as protein degree. The abundance of PPI was calculated using the following equation, which is the PSM count(s) of XL-peptide(s) (or called crosslink) that corresponded to a PPI (Liu et al., 2018a), as a single PPI could be described by multiple species of XL-peptides.

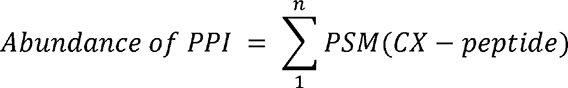

Both the PPI and the abundance of PPI were incorporated by the M1 algorithm of the MONET toolbox to construct the Nuclear Protein-Protein Interaction Modules (NPIMs; Fig 4, S14, S15). The formation of an NPIM required a minimum of two proteins (or called components). Moreover, the entire constituent components of an NPIM can be converted to a single converging node, and the PPI and the abundance of PPI between the two interacting NPIMs can be converted to hybrid edge, respectively (Table S4c; Fig. 5, S14, S15). Consequently, Module-Module Interaction (MMI, or called hybrid edge, defines a higher level of edge existing in between two interacting converging nodes (or two interacting modules). The abundance of MMI is the sum of the abundance of PPIs derived from all interacting components between these two modules. Again, both MMI and the abundance of MMI were further incorporated by the M1 algorithm of the MONET toolbox to produce a module of modules (or NPIMs), which was consequently defined as a Community. A higher level of organization of a Community is defined as a System (Fig 4, S14). The Graph Index, Community-NPIM-nuclear protein component (x-y-z), indicates the position of a nuclear protein within the System 1 (the protein interaction networks as a whole in the nucleus is referred to be System 1, Fig S14). Those NPIMs that failed to be integrated into a Community were defined as the Ungrouped (The graph index of this class of NPIM was defined as 0). The diagrams of NPIM and Community of nuclear proteins were depicted using Cytoscape.

### Bioinformatic Analysis of Arabidopsis Condensate-forming Proteins and Protein Complexes Among NPIMs

The biomolecular condensate-forming proteins were selected from both the literature and PhaSePro database (Mészáros et al., 2020). The polypeptide sequences of these proteins were searched with the eggNOG-Mapper against eukaryotic orthogroup (euNOG) HMMs to obtain their euNOGs accessions. The nuclear Arabidopsis ortholog proteins used in the modulomics study were simultaneously converted into euNOG by the same method. The Arabidopsis orthologs that have the same euNOGs as do the documented condensate-forming proteins were defined as the putative condensate-forming proteins (Table S4c; Fig. 4). The software, MobiDB (Piovesan et al., 2021) and PLAAC (Lancaster et al., 2014), were used to find the proteins possessing Intrinsically Disordered Regions (IDR) and prion-like sequences, respectively. The proteins that met one of the two criteria was considered as a predicted condensate-forming protein under conditions: 1) the IDR score in MobiDB ≥ 0.5; 2) the CORE score in PLAAC > 0. The plant nuclear protein complexes were firstly selected from a gel filtration-mass spectrometry (or CF-MS) study (McWhite et al., 2020). If the number of components of a plant nuclear protein complex have an overlapping equal to and larger than 60% of that of a NPIM, the NPIM is therefore defined to be a module capturing the protein complex (or JC > 0.6, Tang et al., 2021).

### Construction of Control- and Hormone-specific Module and Community Variant

The Control-specific PPI refers to those PPIs resulted from a combination of both the hormone significantly down-regulated XL-peptide(s) and the hormone unaltered one(s), whereas the Hormone-specific PPI stands for those PPIs derived from a combination of both the hormone significantly up-regulated XL-peptide(s) and the hormone unaltered one(s) (Table S5a-d). The Common PPI defines those PPIs that the hormone unaltered XL-peptide(s) or those hormone simultaneously and significantly down-and up-regulated XL-peptide(s). Based on these definitions, we segregated overall Arabidopsis ortholog PPIs into Control (including both Control-specific and Common PPIs; Table S5a-b) and Hormone (including both Hormone-specific and Common PPIs; Table S5c-d) PPI datasets. The modulomic analysis of Control and Hormone datasets using the MONET toolbox constructed Control module variants (CV, control variant) and Hormone module variants (TV, treatment variant), respectively (Table S5e-f; Fig 5A). Similarly, the modulomic analysis of converging nodes and merging edges that were congregated from the combination of PPI information of modular components from both Control and Hormone datasets generated Control and Hormone Community variants, respectively (Table S5e-f; Fig 5A).

The comparison between module variant CV and TV was accessed by the Jaccard Coefficient (JC, Tang et al., 2021) calculated based on the modular components’ identification using the following equation

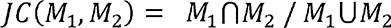

where M_1_ and M_2_ stands for CV and TV module variant, respectively.

After an exhausted comparisons between all CVs and all TVs, the module variant of a maximum JC < 0.6 was defined as Control- or Hormone-specific module variants, whereas the module variant of a maximum JC ≥ 0.6 as the Steady-State Module Variants (Table S5e-g; Fig 5B, S17-18; Tang et al., 2021).

A similar comparison method was performed at the Community level to determine the Control- or Hormone-specific Communities (Table S5e-f; Fig 5C). The differences between the Control and Hormone communities were also accessed by their GO enrichment analysis, which was firstly performed on the top 10% or 50% of the most interactive proteins (or proteins of the highest degree) of each Community. JC was applied to compare the significantly enriched GO terms (p ≤ 0.05) of each pair of Communities. Likewise, the Community of a maximum JC < 0.6 was defined as Community of a specific function. Heatmap was used to depict the level of difference and similarity between CV and TV Communities (Table S5i-l; Fig S20).

### Validation of Intra-crosslinks

All proteins with intra-crosslinks were searched against The Protein Data Bank (PDB) based on the primary sequence similarity. The PDB structures with ≥ 0.9 identities were fetched out as the validation sets. The measurement of Cα-Cα distance was accomplished by PyMOL. Random Cα-Cα distances were calculated by all the possible lysine-lysine residue pairs of the corresponding protein structures.

### Designing and Production of Polyclonal Antibodies

Rabbit polyclonal antibodies were raised against the synthetic peptide _914_YMDDSKKPPVGQGLNKP_930_ of nuclear pore complex protein NUP98A (<u>Nu</u>cleo<u>p</u>orin <u>98A</u>, XP_003539631.1) and _70_KRHRKVLRDNIQGIT_84_ of histone H4 (NP_001237495.1), whereas rat polyclonal antibodies against the synthetic peptide _306_CYDWSKGAENHNPAT_320_ of RAE1 (<u>R</u>ibonucleic <u>A</u>cid <u>E</u>xport <u>1</u>, NP_001242030.2) and _261_PTVTWADPKNSPDH_274_ of hnRNPQ (<u>H</u>eterogeneous <u>N</u>uclear <u>R</u>ibo<u>n</u>ucleo<u>p</u>rotein <u>Q</u>, XP_003521823.1). All polyclonal antibodies were produced in the GL Biochem company (Shanghai, China).

### Preparation of Plant Tissue Section and Immunostaining

The formaldehyde fixation and the section of soybean tissues were prepared as previously described (Paciorek et al., 2006). Briefly, the 7-year-old seedlings were cut into small pieces using razor blades (Leica) and fixed with 4% para-formaldehyde in PBS. The dehydration of tissue was achieved using different gradients of ethanol (25%, 50%, 75% and 96%), followed by paraffin wax embedding and microtome (HM325, Thermo Scientific Inc., Waltham, MA, USA) section. The materials were subsequently dewaxed using xylene and rehydrated with 99%, 90% and 50% ethanol. PBS containing 2% BSA was used for blocking the tissue sections at room temperature and further incubated with the anti-polyclonal antibody for 12 hr at 4 °C. After three times of washing (10 min per washing) of plant tissue sections with the microtubule-stabilizing (MTSB) buffer, containing 50 mM PIPErazine-N,N′-bis[2-ethanesulfonic acid] (PIPES), 5 mM MgSO_4_, and 5 mM EGTA (pH 6.9), these sections were incubated with 1:1000 goat antirat IgG (H+L), Alexa Fluor 568 (Thermo Fisher Scientific) and 1:1000 goat antirabbit IgG (H+L), Alexa Fluor 750 (Thermo Fisher Scientific) for 2 hr at room temperature. Consequently, the materials were washed 4 times with MTSB buffer (10 min per washing), followed by being incubated with a drop of ProLong antifade mountants (Thermo Scientific Inc., Waltham, MA, USA) at room temperature for 24 hr.

### Super-resolution Imaging and Colocalization Analysis

Super-resolution imaging was performed using the Single Molecular Localization Microscopy (SMLM) mode of ZEISS Elyra 7 with Lattice SIM for each pair of proteins. Samples were excited with a 561 nm diode-pumped solid-state (DPSS) laser (500 mW) and 642 nm diode laser (500 mW), respectively. The laser power mode was set as TIRF-uHP. The fluorescence signals were subsequently observed by an objective (Plan-Aprochromat 63x/1.4 oil DIC M27, ZEISS) with a 1.6 x lens. Emission light was collected by an electron-multiplying charge-coupled device (EMCCD, Andor, iXon 897) after passing through an emission filter SBS LP 560. The fast frame mode was enabled. The time series were set as 5000 frames for one super-resolution experiment after enabling the T-lapse. For image acquisition, the image size was set as 512 x 512 pixels. The 405 nm (50 mW) diode laser was used as activation power with an intensity of 1%, and the 561nm and 642nm lasers were served as excitation power with an intensity of around 20%. In addition, a 50 ms exposure time was used for imaging. Consequently, the blanking of the fluorescence dye molecules was recorded and subsequently form a super-resolution image with a 20 nm to 30 nm lateral resolution. Collected data were processed using the software Zen Lite (Zeiss) and ImageJ (Schneider et al., 2012). The colocalization of the proteins was accessed by Manders’ Overlap Coefficient (MOC, Dunn *et al*., 2011) value using color C package (Ahmed et al., 2019) of R.

### Immunoblot Assays

The plant proteins were extracted with Protein Extraction Buffer containing 125 mM Tris– HCl (pH 7.4), 1%SDS, 10% glycerol, 5 mM DTT, 50 mM NaF, 1% glycerol-2-phosphate, 20 mM EGTA, 20 mM EDTA, 1 mM PMSF, and EDTA-free protease inhibitors. The extraction solution was subjected to centrifugation at 13,200 rpm for 10 min at 4 °C to remove plant cell debris. The solution was then cooled on ice for 1 min and mixed with 3 volumes of cold acetone/methanol (12:1, v/v) mixture for 2 hr to precipitate proteins. The protein pellet was rinsed with a mixture of cold acetone/methanol/H_2_O (12:1:1.4, v/v/v) to remove residue pigment before it was air-dried and redissolved in a Resuspension Buffer (50 mM Tris–HCl, pH 6.8, 8 M Urea, 5 mM DTT, 1% SDS, and 10 mM EDTA). The final concentration of the proteins was measured using the protein DC assay and calculated based on a BSA protein standard curve. The resulting protein samples of 100 µg were loaded into each lane on a 10% SDS-PAGE gel before it was immobilized onto a PVDF membrane (GE Healthcare). The immobilized membranes were blocked with 5% (w/v) milk in TBST (Tris-buffered saline, 0.1% (v/v) Tween 20) for 1 hr at room temperature, followed by incubation with primary antibody for 1 hr at room temperature and 3 washes (10 min each) with TBST. The membranes were then incubated with secondary antibody for 1 hr at room temperature (anti-rabbit or anti-rat, conjugated with HRP, Bio-Rad), followed by 3 washes (10 min each) with TBST. LuminataTM Forte Western HRP Substrate (Millipore) was used for membrane exposure and signal detection. Immunoblot quantification was performed using ImageJ. The following primary antibodies: anti-hnRNPQ, anti-NUP98A, anti-RAE1 and anti-H4 were used in this study. The protein signal of histone H4 was used as the loading control.

### Biological Response Assay of Arabidopsis *nup98a* and *rae1* Mutant

Surface-sterilized etiolated *Col-0*, *rae1* and *nup98a* seeds of Arabidopsis were geminated on half-strength MS-agar plates under the dark condition for 1 day, followed by 5 ppm hormone treatment (Guzmán and Ecker, 1990; Hoffman et al., 1999) for 4 days at room temperature under darkness condition. The concentration of the hormone (ppm) was measured by Perkin Elmer Clarus 580 gas chromatography. For each independent replicates, approximately 40 - 50 hypocotyls for each genotype were measured by ImageJ and were represented as mean ± SEM. The comparison of the hypocotyl length between *Col-0*, and mutants (*rae1*or *nup98a*) was assessed using the two-tailed student’s *t-*test.

### Generation of Random Modules and Communities

The classical Erdős–Rényi model was used to generate the random protein-protein network with equivalent nodes and edges of the nuclear PPI network. In total, 1000 random networks were generated. The modulomic analysis (described above) was performed on the random PPI networks to generate Modules and Communities. The degree of components, number of modules and communities were analyzed using R.

### Comparison of Nuclear Modules and Communities Generated from Human XL-MS data

The human XL-MS data was retrieved from documented datasets provided online by Fasci *et al*., 2018 and Wheat *et al*., 2021. The human nuclear proteins were classified based on the protein localization information on THE HUMAN PROTEIN ATLAS. Only the hetero-PPIs derived from two distinct nuclear proteins were selected for further analysis. Modulomic analysis was thereafter performed on the human nuclear PPIs to generate Modules and Communities. The protein components of both human and plant datasets were converted into euNOGs to allow the direct comparison. Finally, the comparison between human and plant Modules and Communities were performed as described above.

### Statistical Analysis

The statistical significance of the results was assessed using two-tailed student’s t-test, with significance represented by *, **, and *** at p < 0.05, p < 0.01, and p < 0.001, respectively. Quantitative data were represented as mean ± SEM. Statistical methods used by SQUA-X to quantify the XL-peptides included the batch effect adjustment, t-test and BH-FDR (Liu et al., 2018b, 2018c).

### Nomenclature and Definition

*Protein-Protein Interaction (PPI)*: the protein interaction of protein A and B, while is determined by the XL-peptides of these two proteins.

*Hetero-PPI*: a protein-protein interaction derived from proteins of the discrete polypeptide sequences.

*Homo-PPI*: a protein-protein interaction derived from proteins of the same polypeptide sequences.

*PPI Abundance (or called weight)*: the PSM count(s) of XL-peptide(s) that was corresponding to the PPI.

*Node*: an intersection protein of a PPI network.

*Edge*: a connection between two nodes of a PPI network.

*Protein Degree*: the number of connections on a protein node of a PPI network.

*Component*: the modular protein component in a module, which is ranked according to the protein degree from high to low.

*Cell Graph*: a cluster of protein-interacting systems generated by modulomics.

*Module*: a cluster of interacting components generated by modulomic analysis.

*NPIM: Nuclear Protein-Protein Interaction Module*

*Converging Node*: a single node that is combined by the entire constitute components of a module.

*Hybrid Edge:* a combinatorial connection between two converging nodes (modules), which is determined by the combination of PPI information (containing both PPI and the abundance of PPI) of components positioned within these two modules under consideration.

*Module-Module Interaction (MMI)*: a hybrid edge

*MMI Abundance*: sum of the abundance of PPIs derived from all interacting components between these two modules in convern.

*Module Degree*: the number of connections on a module of a MMI network.

*Community*: a cluster of interacting modules generated by modulomics.

*Community-Community Interaction (CCI)*: a higher level of interaction existed in between two interacting communities, which is determined by the MMIs among the modules in these two communities.

*CCI Abundance*: sum of <u>MMI abundance</u> derived from all interacting modules between these two communities.

*Community Degree*: the number of connections on a community of a CCI network.

*Graph Index*: an index of a component within communities and modules, which follows the pattern of “community-module-component (x-y-z)”. Similarly, *Graph Index* of a module is defined as community-module (x-y).

*System*: a cluster of interacting communities generated by modulomic analysis. Nucleus is defined as System 1.

*System-System Interaction (SSI)*: a higher level of interaction existed in between two interacting systems, which is determined by the CCIs among communities of these two systems. There are multiple Systems in a cell graph of PPI.

*SSI Abundance*: sum of CCIs derived from all interacting communities between these two systems.

*Control-specific PPI*: a PPI that is resulted from a combination of both the hormone significantly down-regulated XL-peptide(s) and the hormone unaltered one(s).

*Hormone-specific PPI*: a PPI that is resulted from a combination of both the hormone significantly up-regulated XL-peptide(s) and the hormone unaltered one(s).

*Common PPI*: a PPI that is resulted from the hormone unaltered XL-peptide(s) or those hormone simultaneously and significantly down- and up-regulated XL-peptide(s).

*Master module or NPIM*: a module that is generated by the combination of PPIs of various conditions (*e.g.*, Control and Hormone PPIs in this study).

*Module variant*: a variation of master module that is generated by PPIs of a single condition.

*Control-specific module variant*: a Control module variant that has the maximum Jarccard Coefficient < 0.6 as compared with the components of all the Hormone modules.

*Hormone-specific module variant*: a Hormone module variant that has the maximum Jarccard Coefficient < 0.6 as compared with the components of all the Control modules.

*Steady-state module variant*: a Control/Hormone module variant that has the maximum Jarccard Coefficient ≥ 0.6 as compared with the components of all the Hormone/Control modules.

*Master community*: a community that is generated by the MMIs of master modules. *Community variant*: a variation of master community that is generated by MMIs of module variants.

*Control-specific community variant*: a Control community variant that has the maximum Jarccard Coefficient < 0.6 as compared with the components of all the Hormone communities.

*Hormone-specific community variant*: a Hormone community variant that has the maximum Jarccard Coefficient < 0.6 as compared with the components of all the Control communities.

*Steady-state community variant*: a Control/Hormone community variant that has the maximum Jarccard Coefficient ≥ 0.6 as compared with the components of all the Hormone/Control communities.

