## Supplemental Figures for "Capturing the Hierarchically Assorted Modules of Protein Interaction in the Organized Nucleome"

#### Supplemental Figure S1

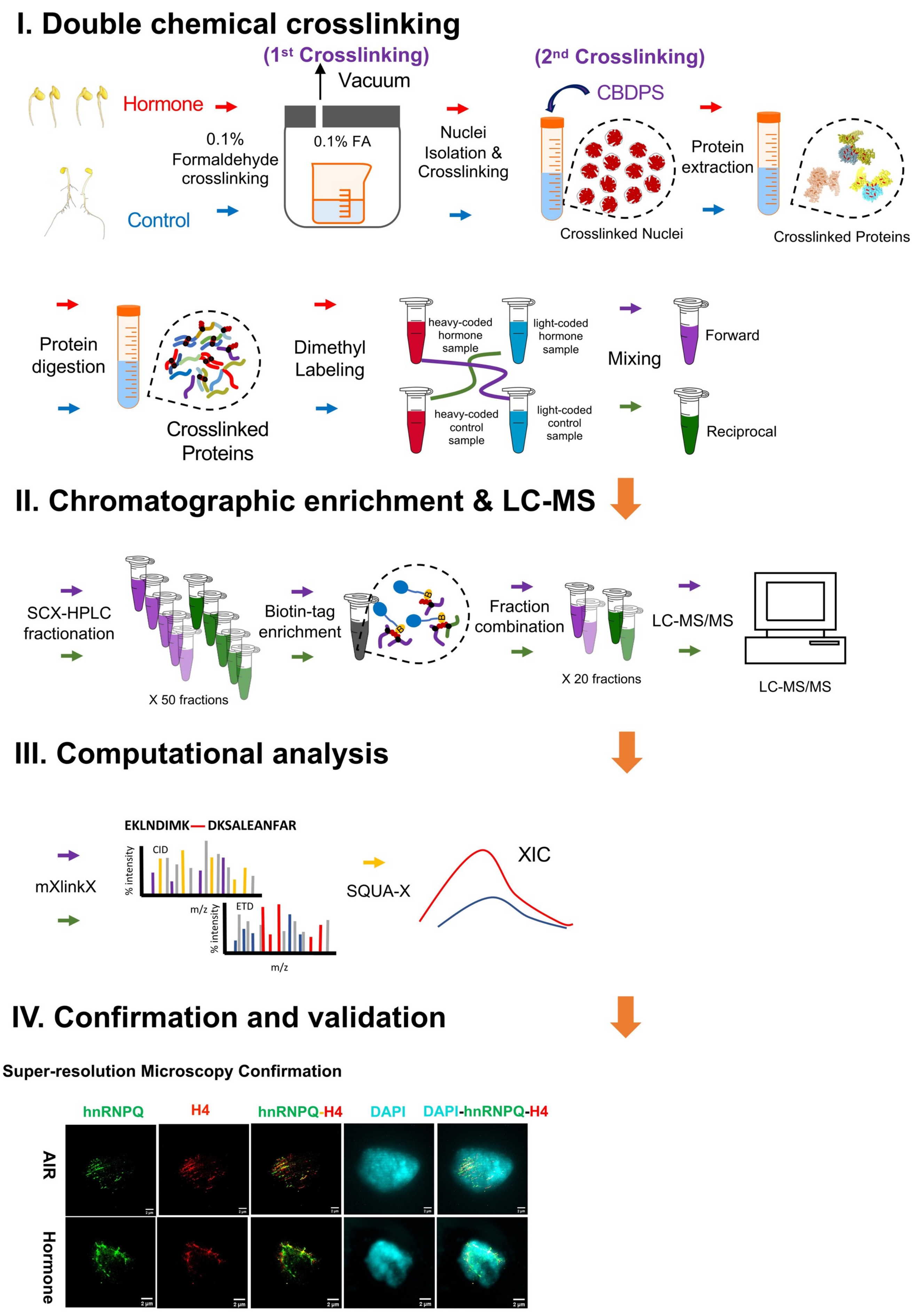

#### Supplemental Figure S1

#### The workflow of 4C quantitative interactomics

The workflow of 4C quantitative interactomics consists of (I) **C**hemical labelling: *i.e. in vivo* formaldehyde fixation (1$^{st}$ crosslinking) of the untreated and hormone-treated seedlings and *in organello* CBDPS crosslinking (2$^{nd}$ crosslinking) of isolated nuclei, followed by protein extraction and digestion as well as finally the dimethyl labeling and mixing of XL-peptides, (II) **C**hromatographic enrichment & LC-MS, which includes SCX-HPLC separation and biotin tag-based enrichment of XL-peptides and LC-MS/MS analysis; (III) **C**omputational analysis, including the XL-peptides searching by the modified XlinkX (mXlinkX) software and the stable isotope-labeling-based quantitation of XL-peptides by SQUA-X software, and (IV) **C**onfirmation and validation, including the confirmation of PPI using the super-resolution microscopy and data deposited in Protein Data Bank, and validation of the function of nuclear proteins using and molecular biology. The corresponding protein, peptide amount and yield is listed in **Table S0a.**

**Supplemental Figure S2**

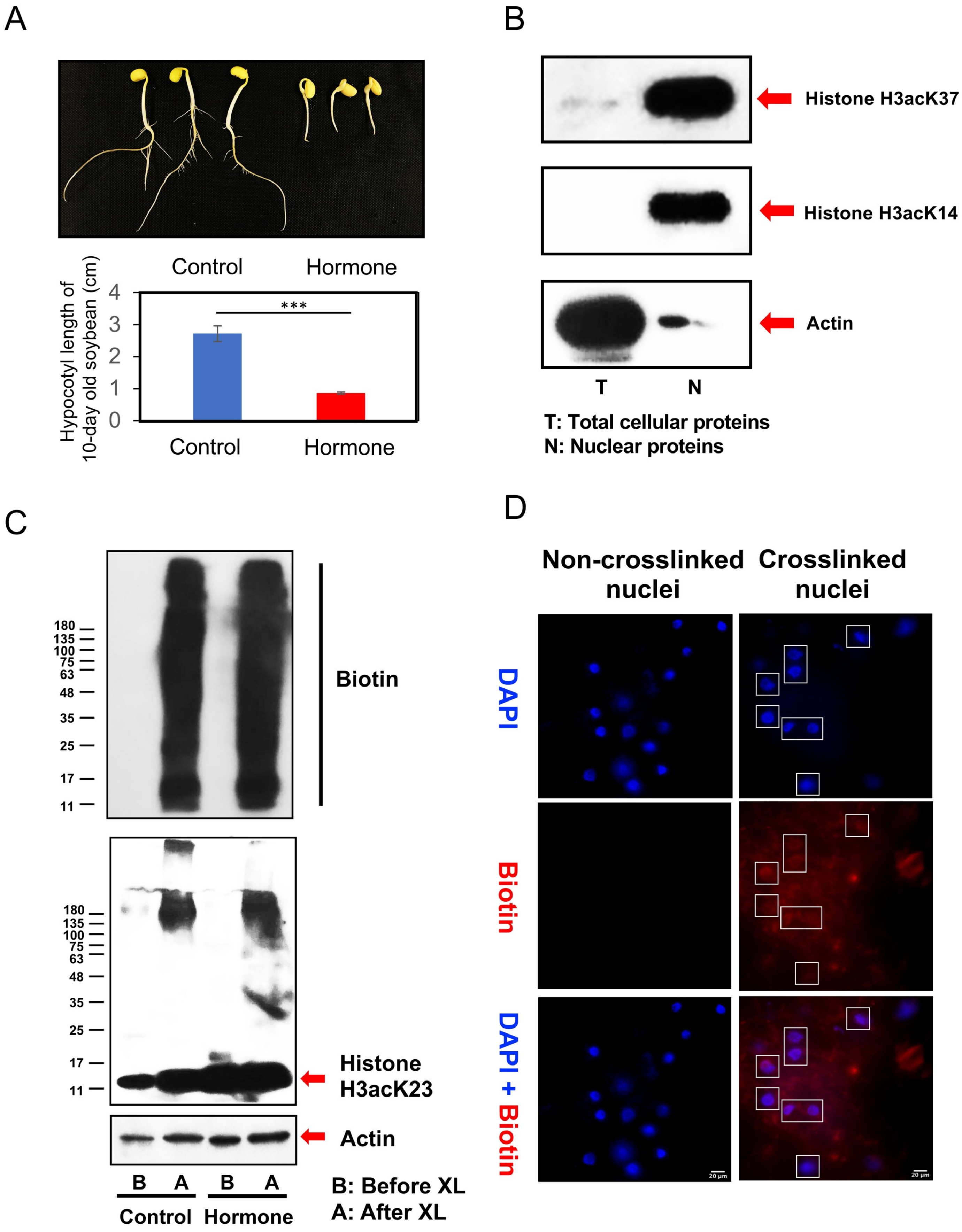

#### Supplemental Figure S2

#### Hormone treatment of plant seedlings and nuclear isolation

(A) The upper panel shows representative photos of the control and hormone-treated plant seedlings. The lower panel shows the difference of the hypocotyl length in between the air- and hormone-treated 10-day old seedlings. The error bar stands for the SEM. Student's *t*-test was performed. *** stands for $p < 0.001$.

(B) Western blots analysis of proteins isolated from tissues (left) and nuclei (right). The anti-histone PTMs, H3acK27 and H3acK14, and anti-actin antibodies were used to show the success of nuclei isolation. T and N stands for total cellular and nuclear proteins, respectively.

(C) Western blot analysis of the non-crosslinked and CBDPS-crosslinked nuclear proteins. The upper panel represents the overall crosslinker-labeled proteins using anti-biotin antibodies, whereas the lower panel represents the crosslinked histone protein complexes and histone monomer. The anti-biotin and anti-histone PTM, H3acK23, antibodies were used to detect the crosslinked proteins and crosslinked histone protein, respectively. Anti-Actin antibodies were used to monitor the protein loading in each lane. B and A indicates the protein sample before and after crosslinking of nuclei, respectively.

(D) Florescence microscopic study of representatives of non-crosslinked (left panels) and crosslinked nuclei (right panels). The upper, middle and lower panel represents the isolated intact nuclei (DAPI), the crosslinking status of isolated nuclei (anti-Biotin) and merged channels (DAPI + anti-Biotin), respectively. The white boxes represent the crosslinked nuclei.

### Supplemental Figure S3

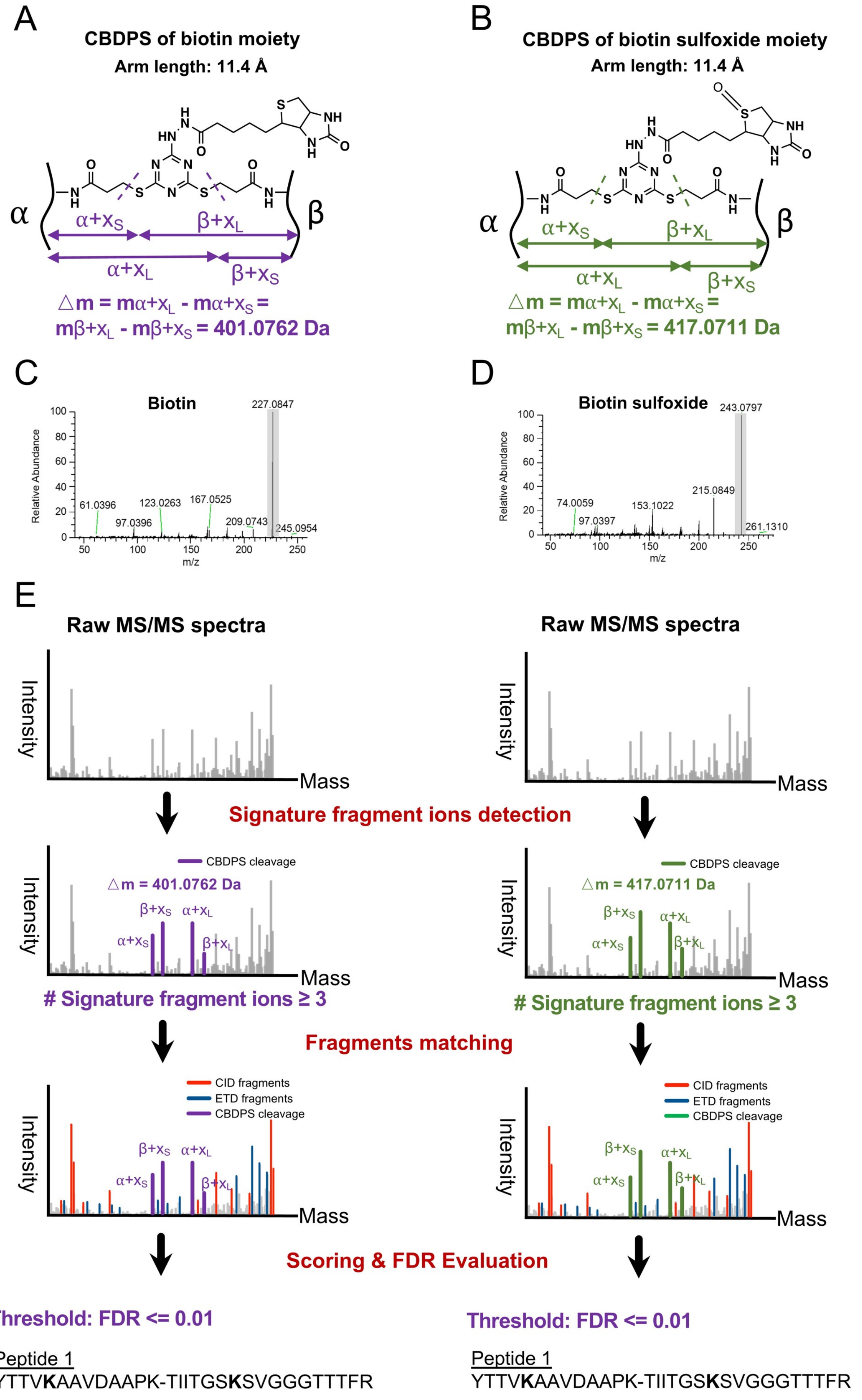

#### Supplemental Figure S3

#### CBDPS crosslinker and workflow of XL-peptides identification

(A - B) Schematic structure of CBDPS (denoted as x) crosslinked peptide α and β. Both CBDPS of biotin moiety (A) and CBDPS of biotin-sulfoxide moiety (B, Kang et al., 2009) were used in searching for XL-peptides by mXlinkX (See Star Methods). The arm length (11.4 **Å**) of the crosslinker is indicated in black letter. The purple and green letters present the information related to the XL-peptide of biotin moiety and biotin sulfoxide moiety, respectively. The dashed lines on the crosslinker stands for the cleavage site in MS. α and β stands for the two peptides crosslinked, whereas $x_S$ and $x_L$ stands for the shorter and the longer arm of crosslinker cleaved by MS, respectively. The four possible signature fragment ions are represented by $\alpha+x_S$, $\beta+x_L$, $\alpha+x_L$ and $\beta+x_s$. The delta mass is determined using an equation ($\triangle m = m\alpha+x_L - m\alpha+x_S = m\beta+x_L - m\beta+x_S$), where the $m\alpha+x_L$, $m\alpha+x_S$, $m\beta+x$ and $m\beta+x_S$ indicates the mass of peptide α plus longer arm of crosslinker, peptide α plus shorter arm of crosslinker, peptide β plus longer arm of crosslinker and peptide β plus shorter arm of crosslinker, respectively. The delta mass for CBDPS of biotin moiety and CBDPS of biotin-sulfoxide moiety is 401.0762 Da and 417.0711 Da, respectively, and they are differ by 15.9949 Da (Kang et al, 2009).

(C - D) MS/MS spectra of biotin (C) and biotin sulfoxide (D). The x and y axis represents m/z value and relative abundance of fragment ions, respectively. The most intensive fragment ions of biotin (m/z 227.0847, 1+) and biotin sulfoxide (m/z 243.0797, 1+) are marked in grey background.

(E) Workflow of identification of XL-peptides by CBDPS of biotin moiety (left panel) and that of biotin-sulfoxide moiety (right panel). The MS precursor ion is subjected to sequential CID-ETD fragmentation (Liu et al., 2015). The raw MS/MS spectra were firstly searched for the signature fragment ions, and spectra of the signature fragment ions ≥ 3 were selected for further processing. The purple and green peaks in the MS/MS fragmentation spectra represent the signature fragment ions resulting from CBDPS of biotin moiety and biotin sulfoxide moiety, respectively. The detected signature fragment ions determine the mass of both peptide α and peptide β, which was followed by fragments matching against the theoretical y-, b-, c-, z-ion series generated from XL-peptide α and β. Red and blue peaks indicate b-, y-ion series and c-, z-ion series, respectively. Eventually, the matched XL-peptides are both scored and evaluated by Percolator-based FDR (Käll *et al.*, 2007).

**Supplemental Figure S4**

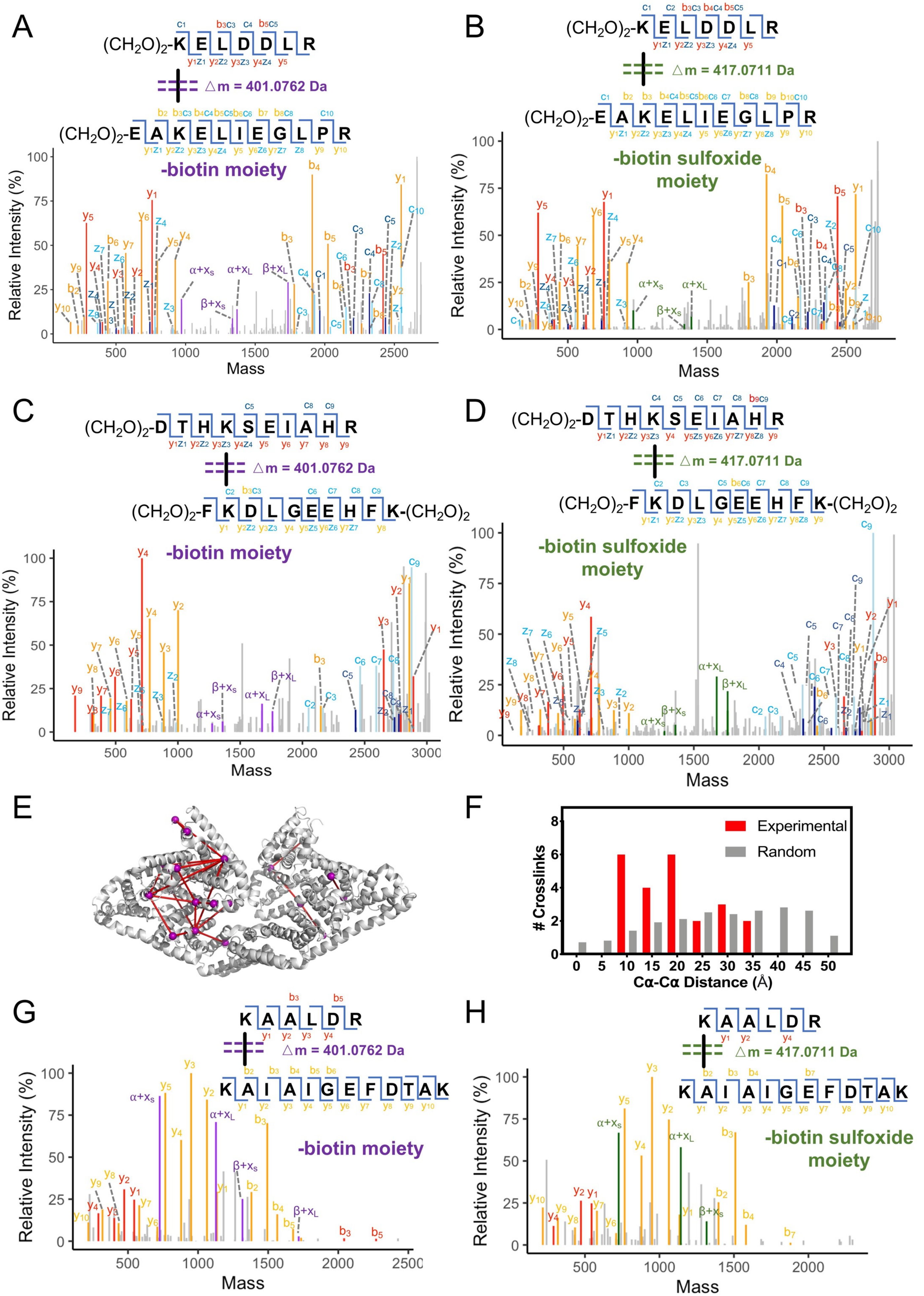

**Supplemental Figure S4**

**Testing of mXlinkX on synthetic peptides, BSA and the crosslinked mitochondria proteins**

(A - B) The MS/MS spectra of the dimethyl $(CH_2O)_2$ labelled synthetic peptides, KELDDLR (α) and EAKELIEGLPR (β), crosslinked with CBDPS of biotin moiety (A) and biotin sulfoxide moiety (B). The purple peaks stand for the signature fragment ions of the XL-peptide crosslinked by CBDPS of biotin moiety. The delta mass △m is 401.0762 Da between the longer and shorter arm of the crosslinker. The purple letters denote the signature fragments of $\alpha+x_S$ (969.5130 Da, theoretical 969.5131 Da), $\beta+x_L$ (1736.8161 Da, theoretical 1736.8170 Da), $\alpha+x_L$(1370.5795 Da, theoretical 1370.5903 Da) and $\beta+x_s$ (1335.7410 Da, theoretical 1335.7398 Da). Similarly, the green peaks stand for the signature fragments of the XL-peptide crosslinked by CBDPS of biotin sulfoxide moiety. The delta mass △m is 417.0711 Da between the longer and shorter arm of the crosslinker. The green letters denote signature fragments of $\alpha+x_S$ (969.5135 Da, theoretical 969.5131 Da), $\alpha+x_L$(1386.5848 Da, theoretical 1386.5845 Da) and $\beta+x_s$ (1335.7423 Da, theoretical 1335.7398 Da). The red letters ($y_x$, $b_x$) and peaks mark the y- and b-ion series from peptide $(CH_2O)_2$-KELDDLR, while the orange letters ($y_x$, $b_x$) and peaks represent the y- and b-ion series from peptide $(CH_2O)_2$-EAKELIEGLPR. The dark blue letters ($y_x$, $b_x$) and peaks mark the c- and z-ion series from peptide $(CH_2O)_2$-KELDDLR, while the light blue letters ($c_x$, $z_x$) and peaks represent the c- and z-ion series from peptide $(CH_2O)_2$-EAKELIEGLPR. The data are listed in Table S0b.

(C - D) The MS/MS spectra of the dimethyl $(CH_2O)_2$ labelled bovine serum albumin (BSA) peptides, DTHKSEIAHR (α) and FKDLGEEHFK (β), crosslinked with CBDPS of biotin moiety (C) and biotin sulfoxide moiety (D). The purple peaks stand for the signature fragment ions of the XL-peptide crosslinked by CBDPS of biotin moiety. The delta mass △m is 401.0762 Da between the longer and shorter arm of the crosslinker. The purple letters denote the signature fragments of $\alpha+x_S$ (1274.6219 Da, theoretical 1274.6367 Da), $\beta+x_L$ (1759.7531 Da, theoretical 1759.7642 Da), $\alpha+x_L$(1675.7250 Da, theoretical 1675.7139 Da) and $\beta+x_s$ (1358.6810 Da, theoretical 1358.6870 Da). Similarly, the green peaks stand for the signature fragments of the XL-peptide crosslinked by CBDPS of biotin sulfoxide moiety. The delta mass △m is 417.0711 Da between the longer and shorter arm of the crosslinker. The green letters denote signature fragments of $\alpha+x_S$ (1274.6580 Da, theoretical 1274.6367 Da), $\beta+x_L$ (1775.7408 Da, theoretical 1775.7584 Da), $\alpha+x_L$(1691.7121 Da, theoretical 1691.7081 Da) and $\beta+x_s$ (1358.6882 Da, theoretical 1358.6870 Da). The red letters ($y_x$, $b_x$) and peaks mark the y- and b-ion series from peptide $(CH_2O)_2$-DTHKSEIAHR, while the orange letters ($y_x$, $b_x$) and peaks represent the y- and b-ion series from peptide $(CH_2O)_2$-FKDLGEEHFK-$(CH_2O)_2$. The dark blue letters ($y_x$, $b_x$) and peaks mark the c- and z-ion series from peptide $(CH_2O)_2$-DTHKSEIAHR, while the light blue letters ($c_x$, $z_x$) and peaks represent the c- and z-ion series from peptide $(CH_2O)_2$-FKDLGEEHFK-$(CH_2O)_2$. The data are listed in Table S0b.

(E) The ribbon structure of BSA built by the crystal structure (PDB: 3V03). Red lines represent the crosslinks within the BSA dimer while the purple dots represent crosslinked lysine sites. The data is listed in Table S0c.

(F) The distribution of carbon distances between the crosslinked lysine pairs of BSA. The red bars represent the distribution of experimental *α* carbon distances of crosslinked lysine pairs of BSA while the grey bars represent that of randomly selected lysine pairs of BSA. The data is listed in Table S0c.

(G - H) The mXlinkX-searched MS/MS spectra of the peptides, KAALDR (α) and KAIAIGEFDTAK (β), measured from mitochondria proteins (Makepeace et al., 2020) that were crosslinked with CBDPS of biotin moiety (G) and biotin sulfoxide moiety (H). The purple peaks stand for the signature fragment ions of the XL-peptide crosslinked by CBDPS of biotin moiety. The delta mass △m is 401.0762 Da between the longer and shorter arm of the crosslinker. The purple letters denote the signature fragments of $\alpha+x_S$ (726.4121 Da, theoretical 726.4024 Da), $\beta+x_L$ (1717.7661 Da, theoretical 1717.7748 Da), $\alpha+x_L$(1127.4880 Da, theoretical 1127.4796 Da) and $\beta+x_s$ (1316.6787 Da, theoretical 1316.6976 Da). Similarly, the green peaks stand for the signature fragments of the XL-peptide crosslinked by CBDPS of biotin sulfoxide moiety. The delta mass △m is 417.0711 Da between the longer and shorter arm of the crosslinker. The green letters denote signature fragments of $\alpha+x_S$ (726.4089 Da, theoretical 726.4024 Da), $\alpha+x_L$(1143.4701 Da, theoretical 1143.4738 Da) and $\beta+x_s$ (1316.7089 Da, theoretical 1316.6976 Da). The red letters ($y_x$, $b_x$) and peaks mark the y- and b-ion series from peptide KAALDR, while the orange letters ($y_x$, $b_x$) and peaks represent the y- and b-ion series from peptide KAIAIGEFDTAK.

**Supplemental Figure S5**

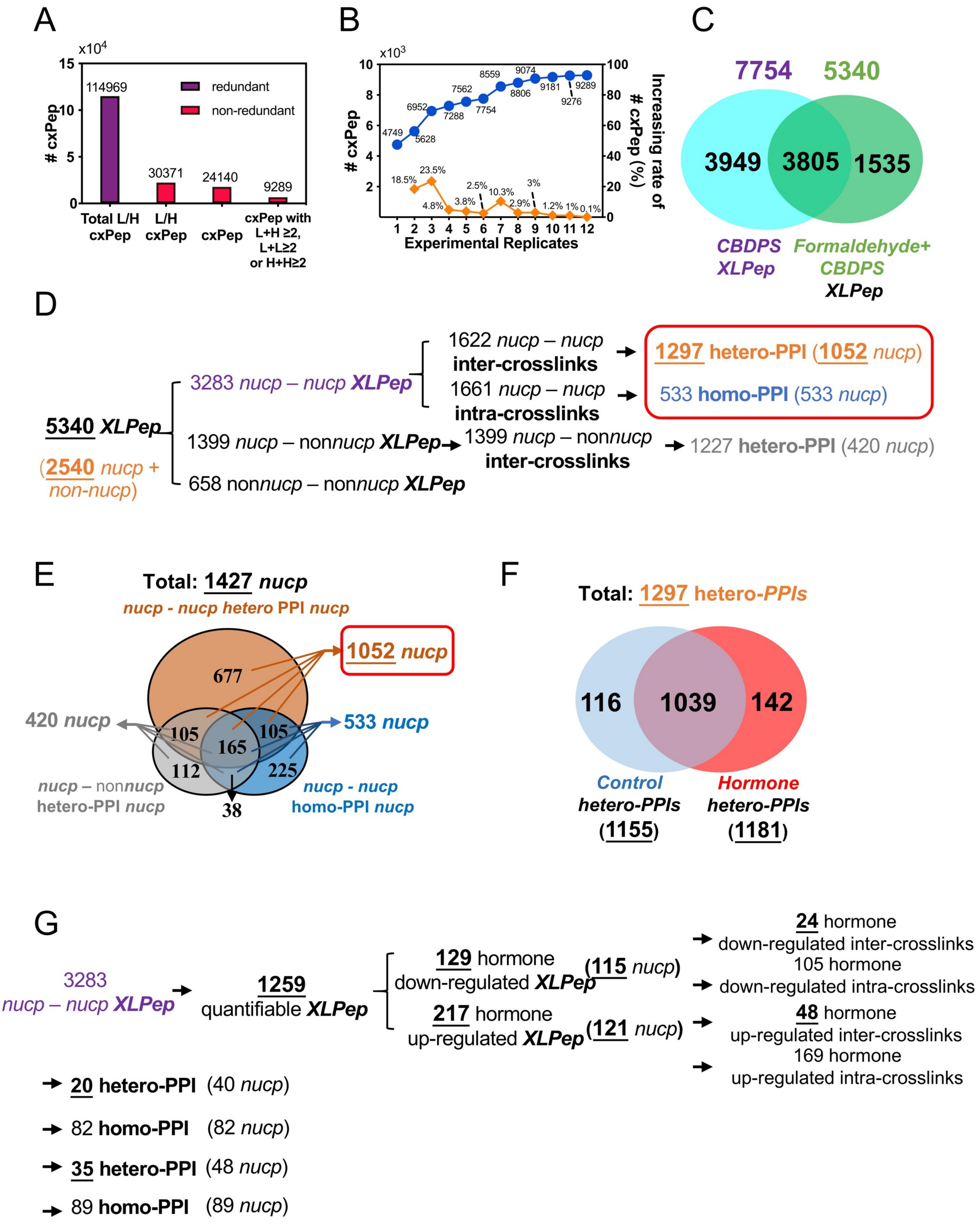

#### Supplemental Figure S5

#### Computational analysis of LC-MS/MS data of crosslinked peptides

(A) The bar represents the number (114,969) of redundant, label-dependent and non-repeatable crosslinked peptides of FDR ≤ 0.01, **Table S1a**), the number (30,371) of non-redundant, label-dependent and non-repeatable crosslinked peptides of FDR ≤ 0.01 (**Table S1b**), the number (**24,140**) of non-redundant, label-independent and non-repeatable crosslinked peptides of FDR ≤ 0.01 (**Table S1c**), the number (**9,289**) of non-redundant, label-independent and repeatable crosslinked peptides of FDR ≤ 0.1 (**Table S1c**). The XLPep stands for XL-peptides.

(B) The accumulation (blue) and cumulating contribution (orange) curve shows a steady increase of the number of non-redundant, label-independent and repeatable XL-peptides (**9,289**) from 12 experimental replicates of 6 biological replicates (**Table S1c**).

(C) A Venn diagram shows the total number of repeatable XL-peptides without (7,754) and with (5,340) formaldehyde pre-fixation (FDR ≤ 0.01), respectively (Table S1c-d).

(D) Diagram represents the data processing procedure of XL-peptides. The 5,340 XL-peptides (corresponding to 2,540 proteins) were classified into 3,283, 1,399 and 658 XL-peptides between two nuclear proteins, one nuclear protein and one non-nuclear proteins and two non-nuclear proteins, respectively (Table S1e-g). The 3,283 XL-peptides between two nuclear proteins could be further divided into 1,622 inter-crosslinks and 1,661 intra-crosslinks. The 1,622 inter-crosslinks of two nuclear proteins, 1,661 intra-crosslinks of two nuclear proteins as well as 1,399 inter-crosslinks of one nuclear and non-nuclear protein could be converted into 1,297 hetero-PPIs of two nuclear proteins (1,052 nuclear proteins), 533 homo-PPIs of nuclear proteins (533 nuclear proteins) as well as 1,227 hetero-PPIs of one nuclear protein and one non-nuclear protein (420 nuclear proteins), respectively (Table S1i). The hetero-PPI and homo-PPI is defined as protein interaction between two proteins with distinct and identical polypeptide sequences, respectively.

(E) Venn diagram showing three categories of 1,427 nuclear proteins: nuclear proteins-nuclear proteins hetero-PPI (1,052, brown), nuclear proteins-nuclear proteins homo-PPI (533, blue) and nuclear proteins-nonnuclear protein hetero-PPI (420, grey). Lines mark the number of nuclear proteins from the combinatorial categories. The classified data are listed in **Table S1i**.

(F) A Venn diagram shows the number of control- (1,155) and hormone-related (1,181) PPIs (1,297) occurred among 1,052 nuclear proteins of different sequences, respectively (Table S1h).

(G) Diagram represents the data processing procedure of quantitative XL-peptides. The 3,283 XL-peptides contained 1,259 quantifiable XL-peptides (see Star Methods for detail). After XIC-based quantification, there were 129 and 217 hormone significantly down- and up-regulated XL-peptides, respectively. The 129 and 217 hormone significantly down- and up-regulated XL-peptides were corresponding to 115 and 121 nuclear proteins, respectively. The 129 hormone down-regulated crosslinks could be separated into 24 inter- and 105 intra-crosslinks, which could be further converted into 20 hetero- (40 nuclear proteins) and 82 homo-PPIs (82 nuclear proteins), respectively. The 217 hormone up-regulated crosslinks could be separated into 48 inter- and 169 intra-crosslinks, which could be further converted into 35 hetero- (48 nuclear proteins) and 89 homo-PPIs (89 nuclear proteins), respectively. The data are listed in Table S1l.

**Supplemental Figure S6**

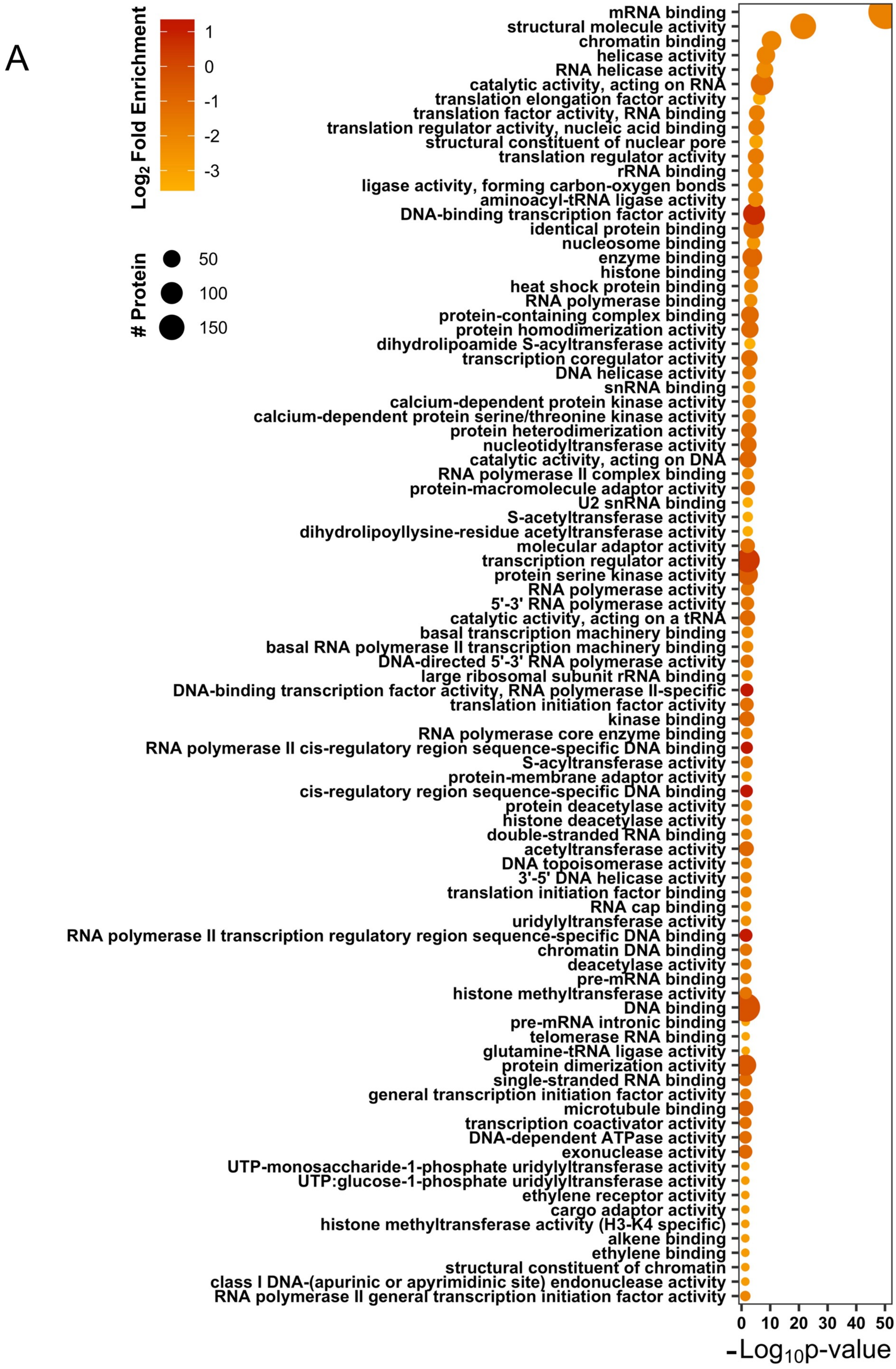

#### Supplemental Figure S6

#### Molecular function analysis of crosslinked nuclear proteins

(A) Bubble plot shows molecular functions enriched from the 1052 nuclear proteins involved in hetero-PPI. Y-axis labels represent the names of molecular functions while the minus base 10 logarithm of *p* value is shown on X-axis. The size and color of the bubble represents number of proteins enriched in each molecular function and their base 2 logarithm of fold enrichment, respectively (**Table S1m**).

### Supplemental Figure S7

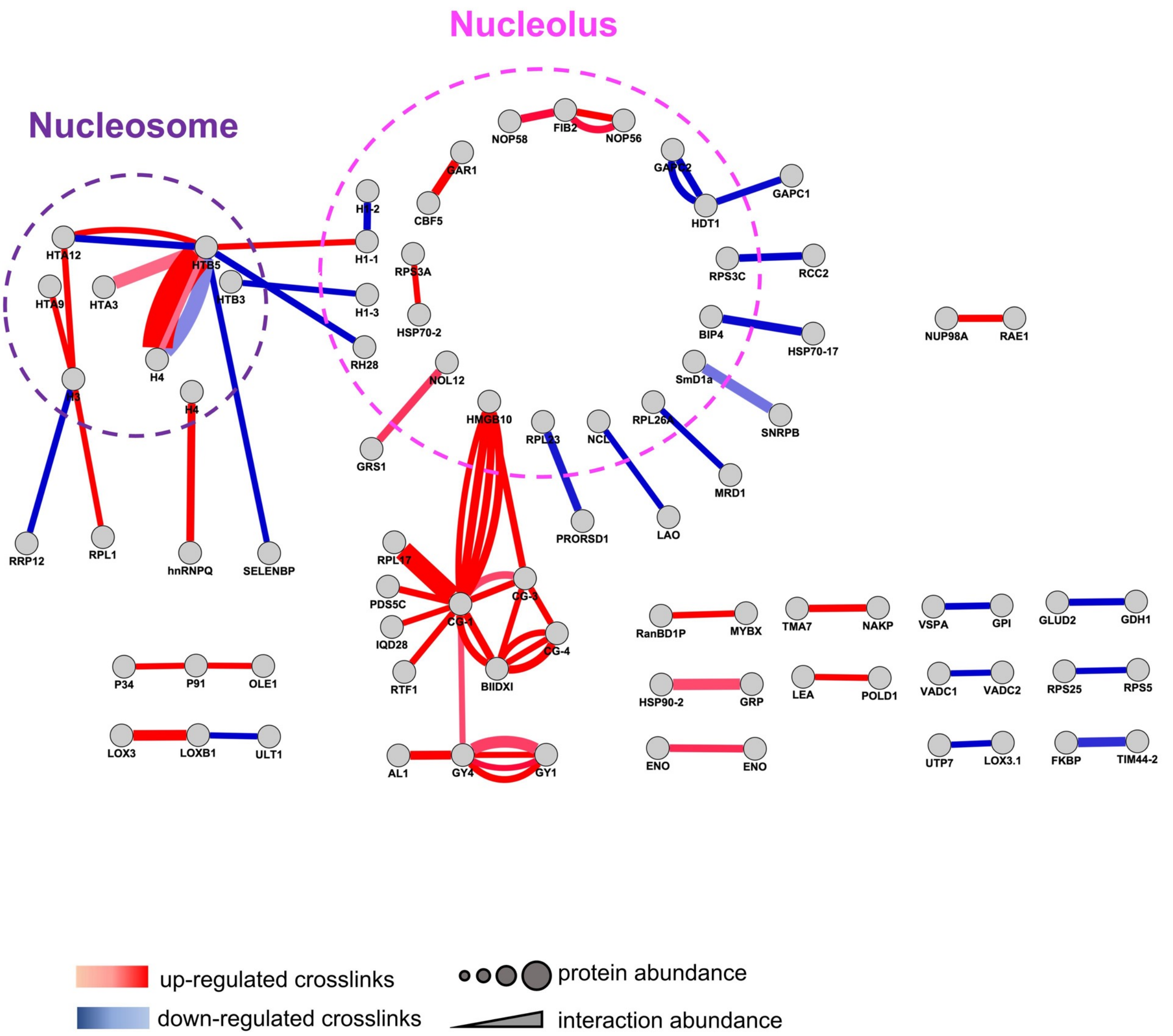

**Supplemental Figure S7**

**Network of quantitative crosslinks among nuclear proteins**

Diagram represents a network of 81 crosslinked distinct nuclear proteins and the abundance of crosslinks. The dot and line represents the protein and significantly regulated crosslinks among nuclear proteins, respectively. The size of the dot represents the protein abundance determined by Proteomic Ruler. The red and blue line represents the hormone significantly up- and down-regulated crosslinks, respectively. The thickness of the color line represents the PSM counts or abundance of the crosslink. The purple and pink dashed circle marks the core histones and nucleolus proteins, respectively. The quantitative data are listed in **Table S1l.**

### Supplemental Figure S8

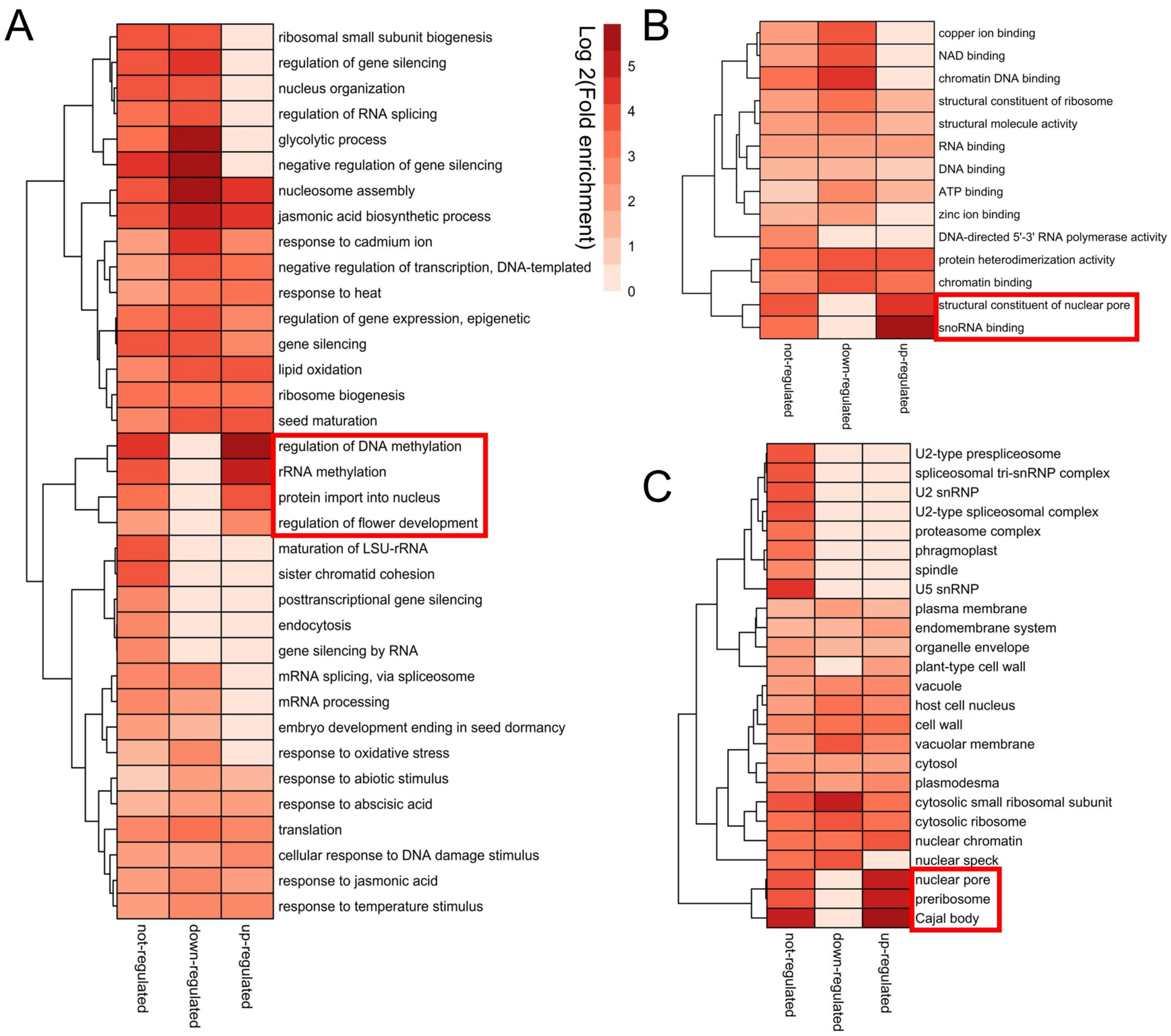

#### Supplemental Figure S8

#### GO analysis of the proteins from hormone significantly regulated crosslinks

(A – C) The heatmap of biological process (A), molecular function (B) and cellular component (C) analysis of proteins from hormone non- (283), down- (115 proteins) and up-regulated (121 proteins) crosslinks (Table S1l). Row represents the name of GO term while column marks the different group of proteins. The color represents the base 2 logarithm of fold enrichment of each GO term (**Table S1n**).

**Supplemental Figure S9**

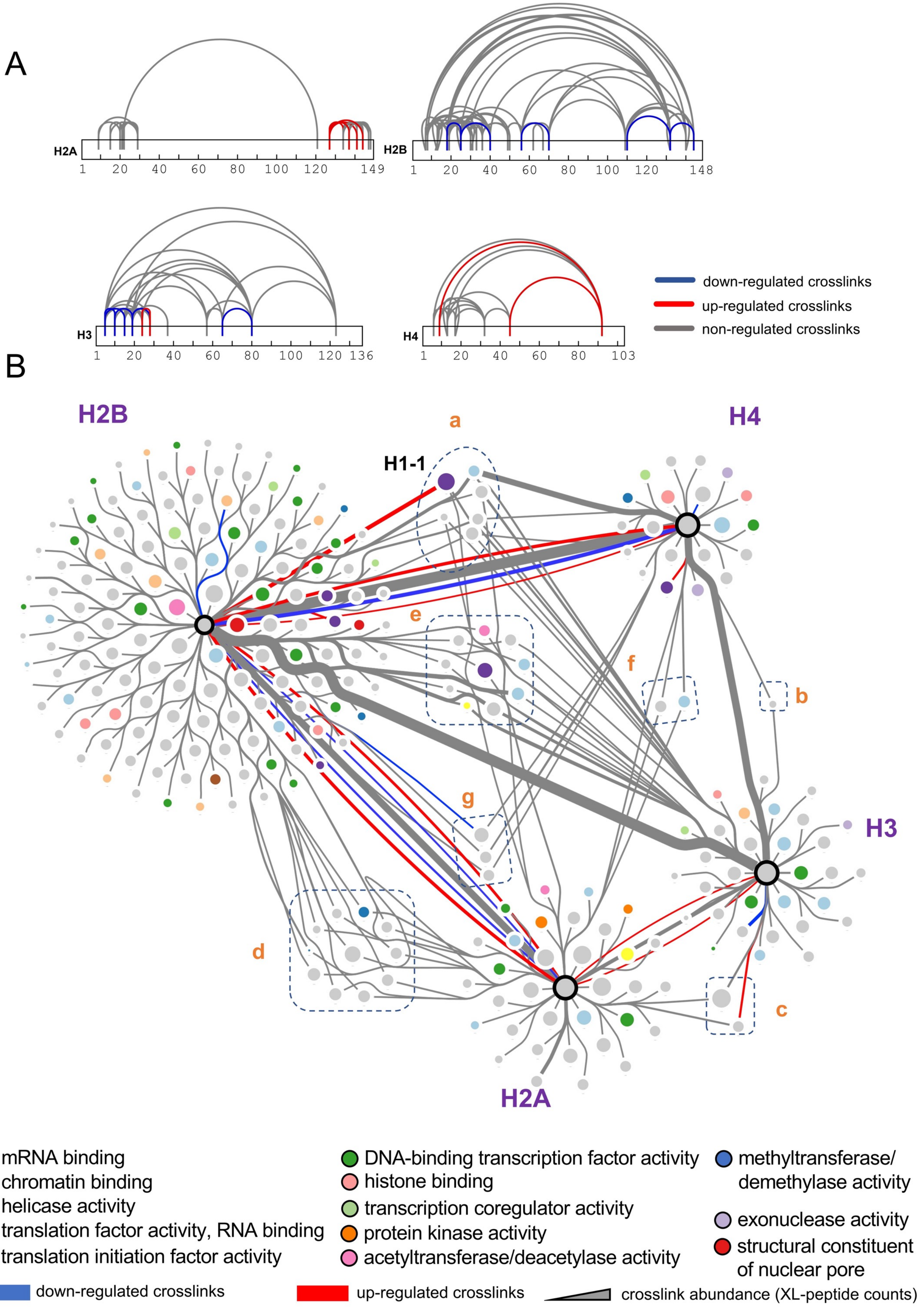

#### Supplemental Figure S9

#### The intra-crosslinks and interactome of core histone proteins

(A) Crosslinking map of 200 intra-crosslinks of core histone proteins (H2A, H2B, H3 and H4). The blue, red and grey lines indicate the hormone down-, up- and non-regulated crosslinks. The data are listed in Table S2a.

(B) The histone proteins are labelled with black circle while 256 interactors of histone octamers are shown as colored dots and grey dots Lines indicate the inter-crosslinks among core histone proteins (87 inter-crosslinks) as well as between histones and their interactors (436 inter-crosslinks). The thickness of line represents the abundance of the crosslinks, which is calculated by the PSM counts. The blue and red edges represent the hormone down- and up- regulated crosslinks, respectively. The light blue-, purple-, wheat-, yellow-, brown-, dark green-, light coral- ,light green-, orange-, pink-, dark blue-, light purple- and red-colored node represents mRNA binding, chromatin binding, helicase activity, translation factor activity (RNA binding), translation initiation factor activity, DNA-binding transcription factor activity, histone binding, transcription coregulator activity, protein kinase, acetyltransferase/deacetylase activity, methyltransferase/demethylase activity, exonuclease activity and structural constituent of nuclear pore, respectively. The common interactors of two, three distinct histone proteins are marked by dark blue dashed boxes and circles, respectively. The data are listed in Table S2a-c.

**Supplemental Figure S10**

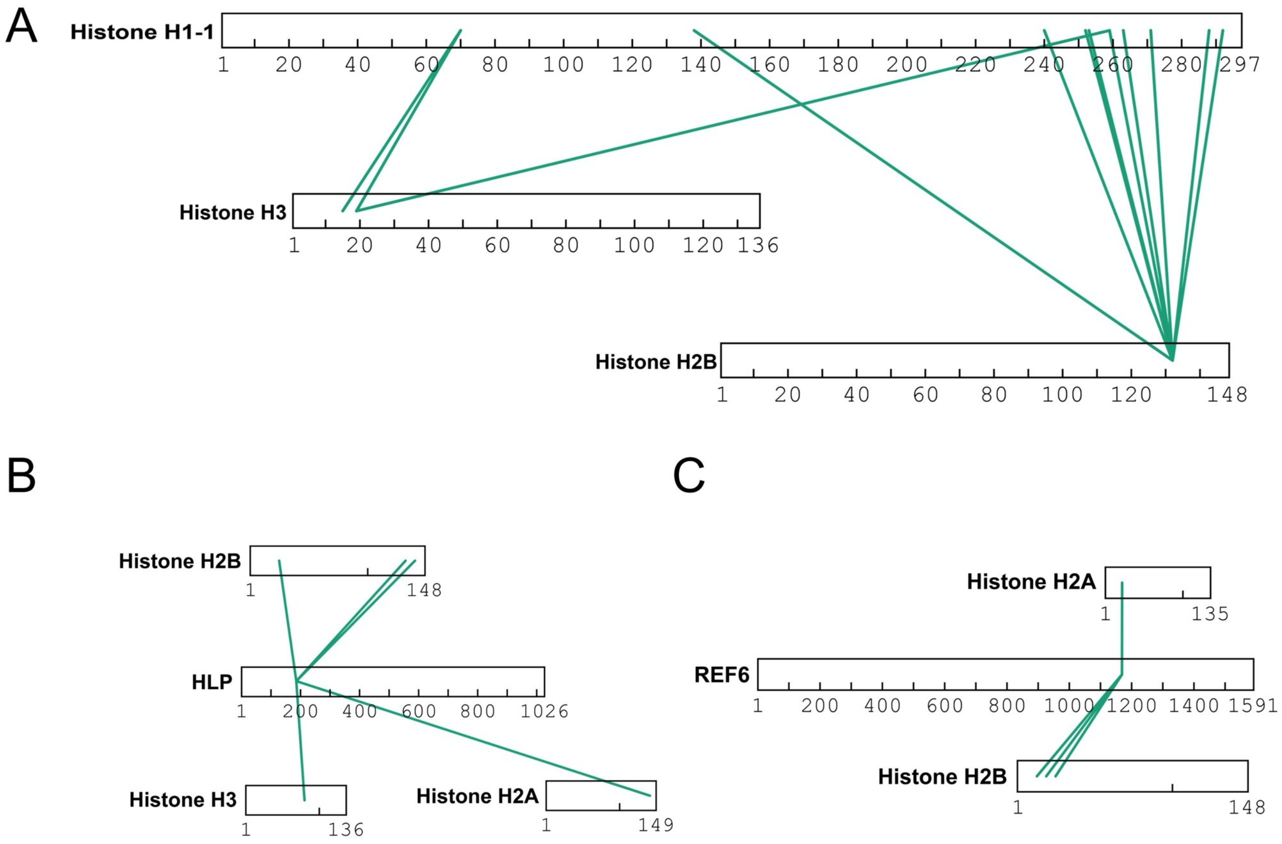

#### Supplemental Figure S10

(A) Crosslinking map of interactions in between histone H1-1 and core histone proteins (histone H2B and histone H3). Green line represents 12 inter-crosslinks of histone H1-1 (**Table S2c**).

(B) Crosslinking map of interactions in between HLP (Homebox-like Superfamily Protein) and core histone proteins (histone H2A, histone H2B, and histone H3). Green line represents 5 inter-crosslinks of HLP (**Table S2c**).

(C) Crosslinking map of interactions in between REF6 (Relative of Early Flowering 6) and core histone proteins (histone H2A, histone H2B). Green line represents 4 inter-crosslinks of REF6 (**Table S2c**).

### Supplemental Figure S11

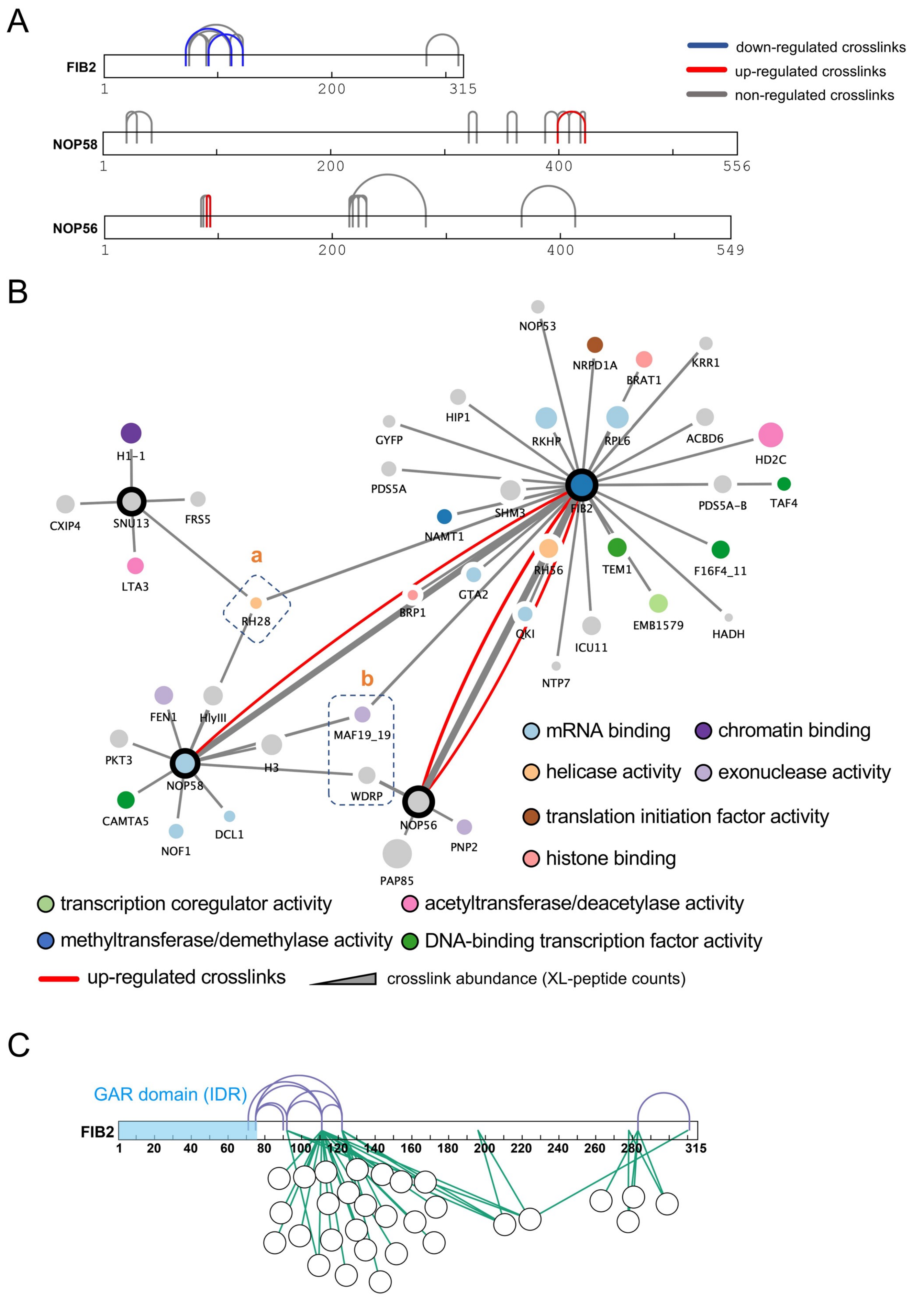

#### Supplemental Figure S11

#### The intra-crosslinks and interactome of core proteins of box C/D snoRNP

(A) Crosslinking map of 33 intra-crosslinks of FIB2, NOP58 and NOP56. The blue, red and grey lines indicate the hormone down-, up- and non-regulated crosslinks. The data are listed in Table S3a.

(B) The box C/D snoRNP proteins are labelled with black circle while 41 interactors of them are shown as colored dots and grey dots. Lines indicate the inter-crosslinks among box C/D snoRNP proteins (11 inter-crosslinks) as well as between box C/D snoRNP proteins and their interactors (57 inter-crosslinks). The thickness of line represents the abundance of the crosslinks, which is calculated by the PSM counts. The red edges represent the hormone up-regulated crosslinks. The light blue-, purple-, wheat-, brown-, dark green-, light coral-, light green-, pink-, dark blue- and light purple-colored node represents mRNA binding, chromatin binding, helicase activity, translation initiation factor activity, DNA-binding transcription factor activity, histone binding, transcription coregulator activity, acetyltransferase/deacetylase activity, methyltransferase/demethylase activity and exonuclease activity, respectively. The common interactors of two distinct box C/D snoRNP proteins are marked by dark blue dashed boxes. The data are listed in Table S1l, S3a-b.

(C) Crosslinking maps of interactions in between FIB2 and its interactors as well as intra-crosslinks of FIB2. Purple and green line represents intra- and inter-crosslinks of FIB2, respectively (**Table S3a-b**). The blue region indicates the GAR (Glycine- and Arginine-rich, 1-75aa) domain of FIB2.

**Supplemental Figure S12**

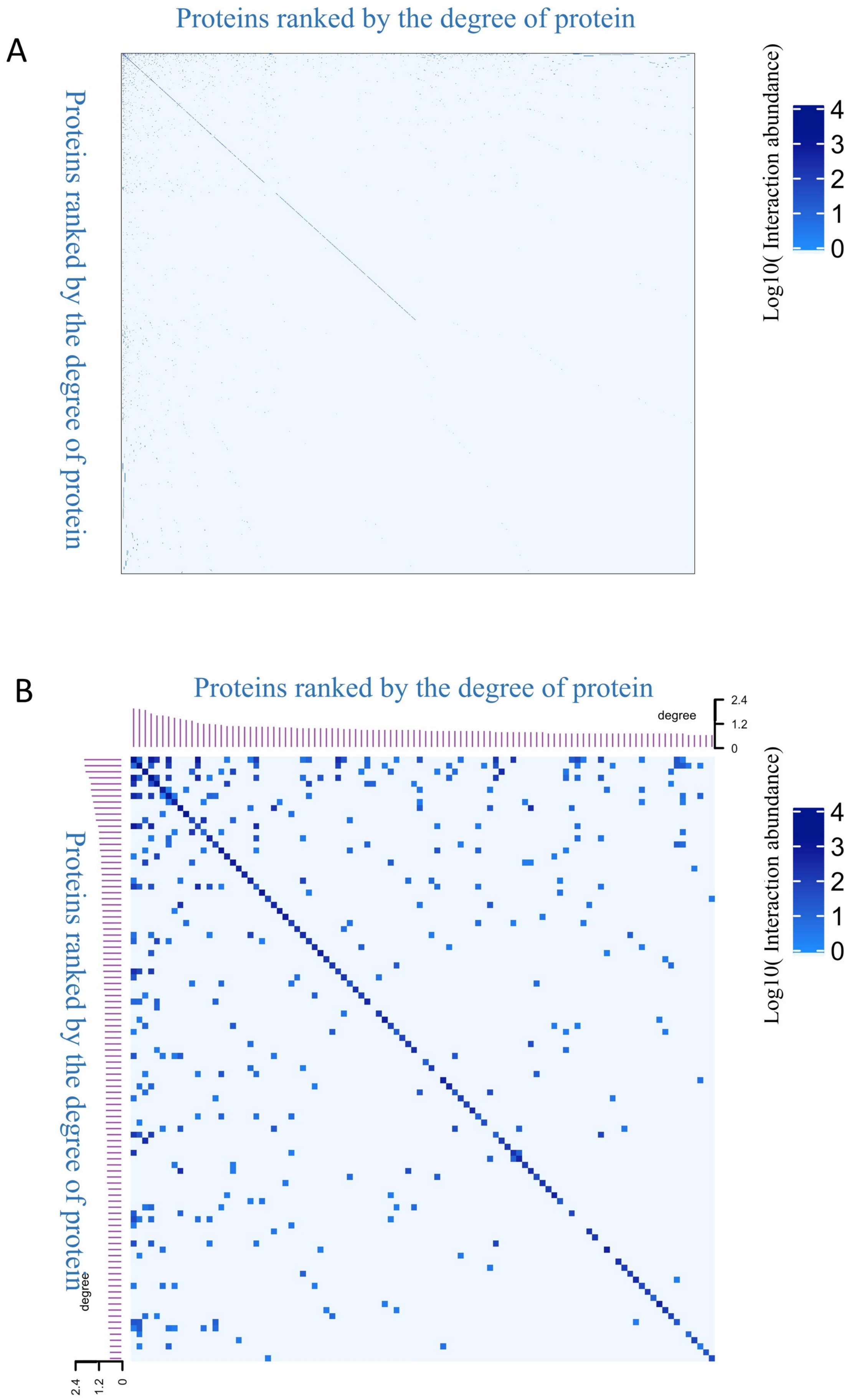

#### Supplemental Figure S12

#### Heatmap of PSM count of XL-peptide derived from nuclear protein

Heatmap (A) and the close-up view of Heatmap (B) of inter- and intra-crosslinks (or PSM counts of XL-peptides) were generated from 1,052 nuclear proteins and 100 most interactive nuclear proteins, respectively. The color intensity of the dot represents the abundance of the protein interaction (PSMs of XL-peptides). Protein components are ranked based on the degree of proteins (Table S1g-h).

**Supplemental Figure S13**

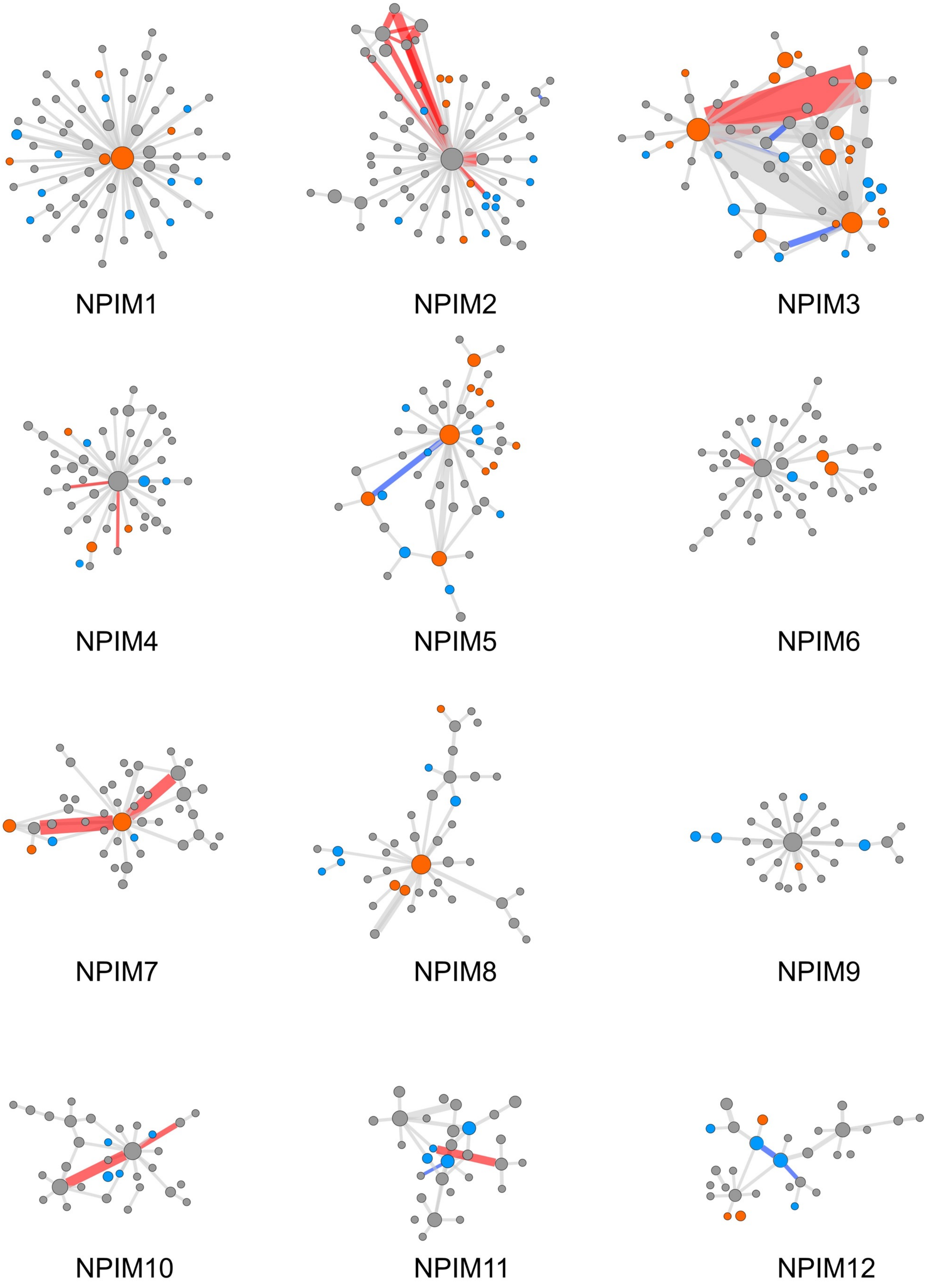

**Supplemental Figure S13**

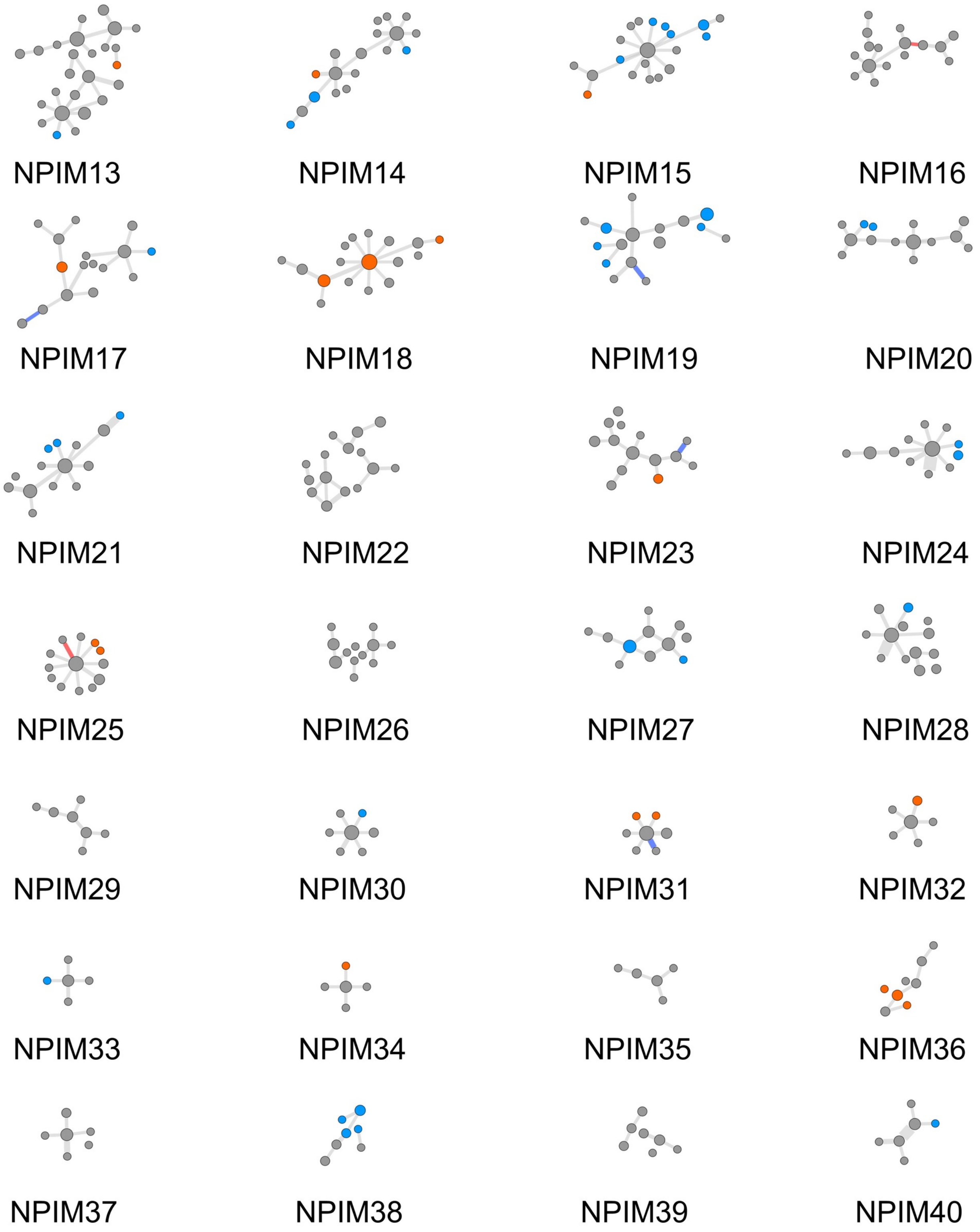

**Supplemental Figure S13**

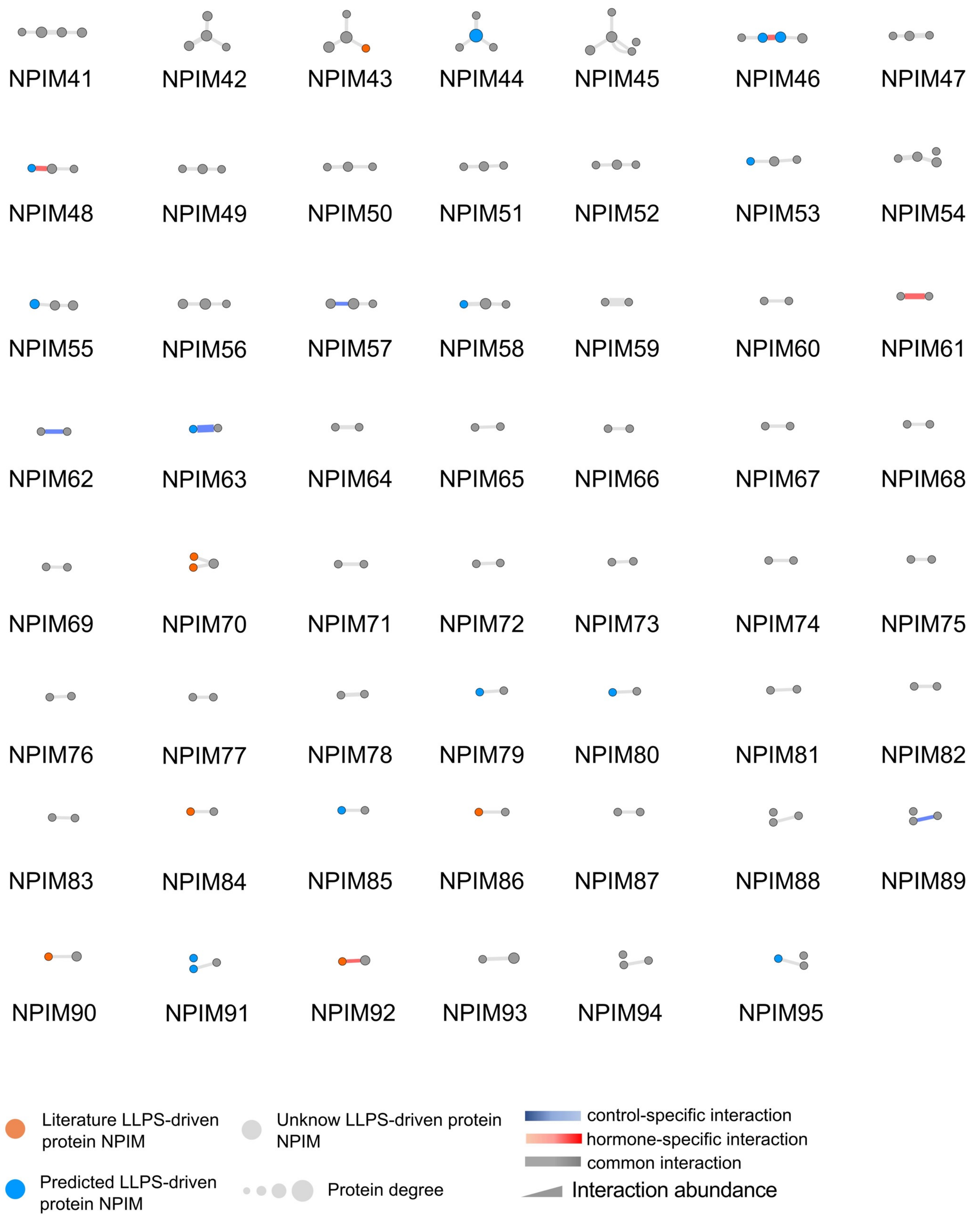

#### Supplemental Figure S13

#### Diagram of nuclear protein interaction module

The diagrams of nuclear protein interaction modules (NPIMs) were generated from PPI data using MONET toolbox. The number of NPIM was assigned by the MONET. The node and edge of module stands for modular protein component and protein-protein interaction, respectively. The size of node stands for the degree of protein. The thickness of edge represents the abundance of protein interaction. The orange, light blue and light grey node represents the literature-annotated condensate-forming, the intrinsically disordered region (IDR)-containing and unknown putative condensate-forming proteins, respectively. The blue, red and grey edge represents the control-specific, hormone-specific and common protein interaction **(Table S4c).**

**Supplemental Figure S14**

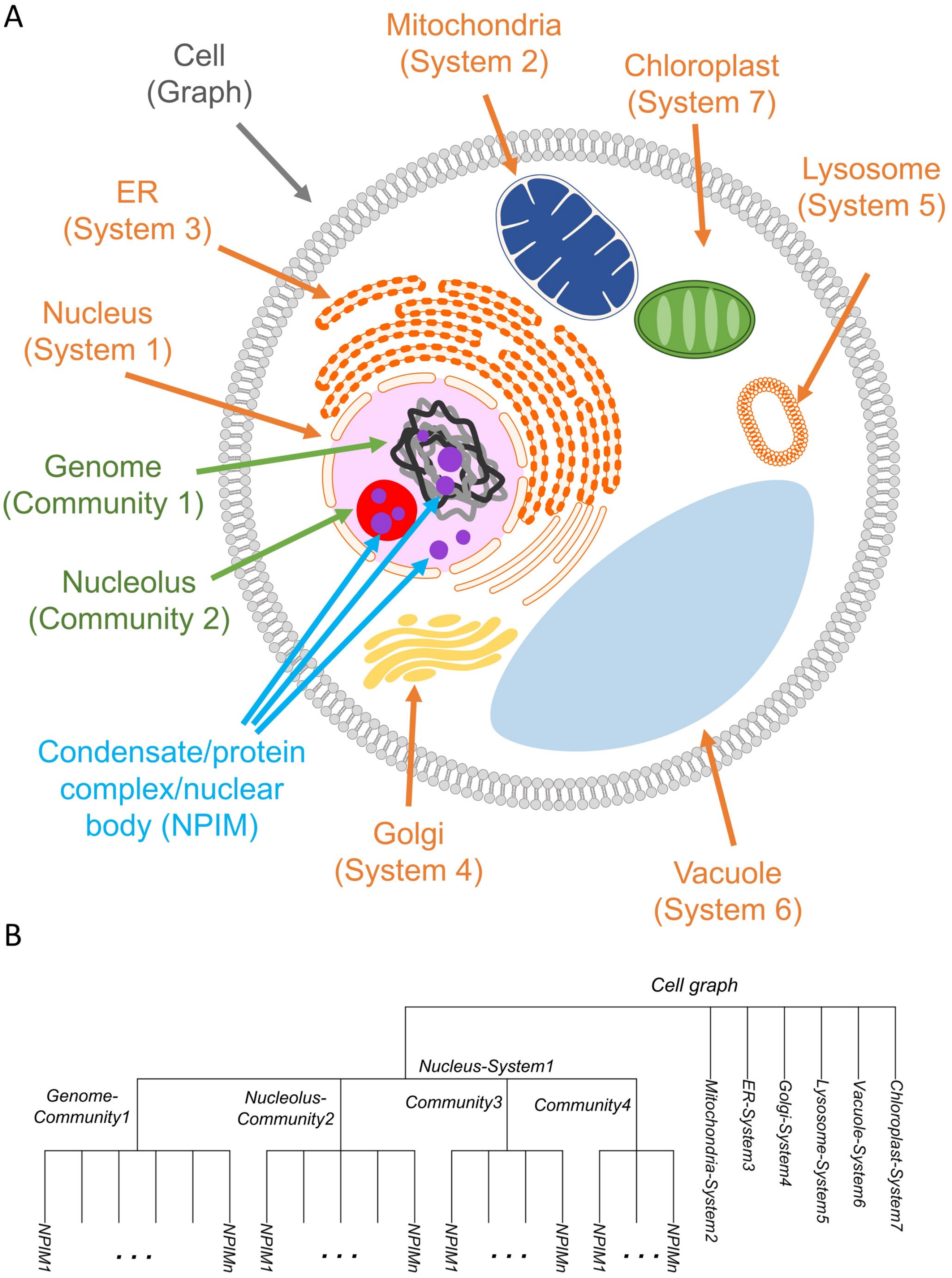

#### Supplemental Figure S14

#### The graph structure of cell

(A) Diagram represents a cell graph. The systems are defined as the organelles of the cell. System 1, 2, 3, 4, 5, 6, and 7 represents the nucleus, mitochondria, ER, Golgi apparatus, lysosome, vacuole, and chloroplast, respectively. The communities are defined as the sub-compartments of the nucleus. Community 1 and 2 represent the genome and nucleolus, respectively. The modules (or called NPIMs) are defined as the sub-compartments of the community. The module represents condensate, protein complex or nuclear body.

(B) Phylogenetic tree revealing the hierarchical and intricate organization of a cell graph. The bottom, middle, and top branch of the tree represents the NPIM, community, and system of the cell graph.

**Supplemental Figure S15**

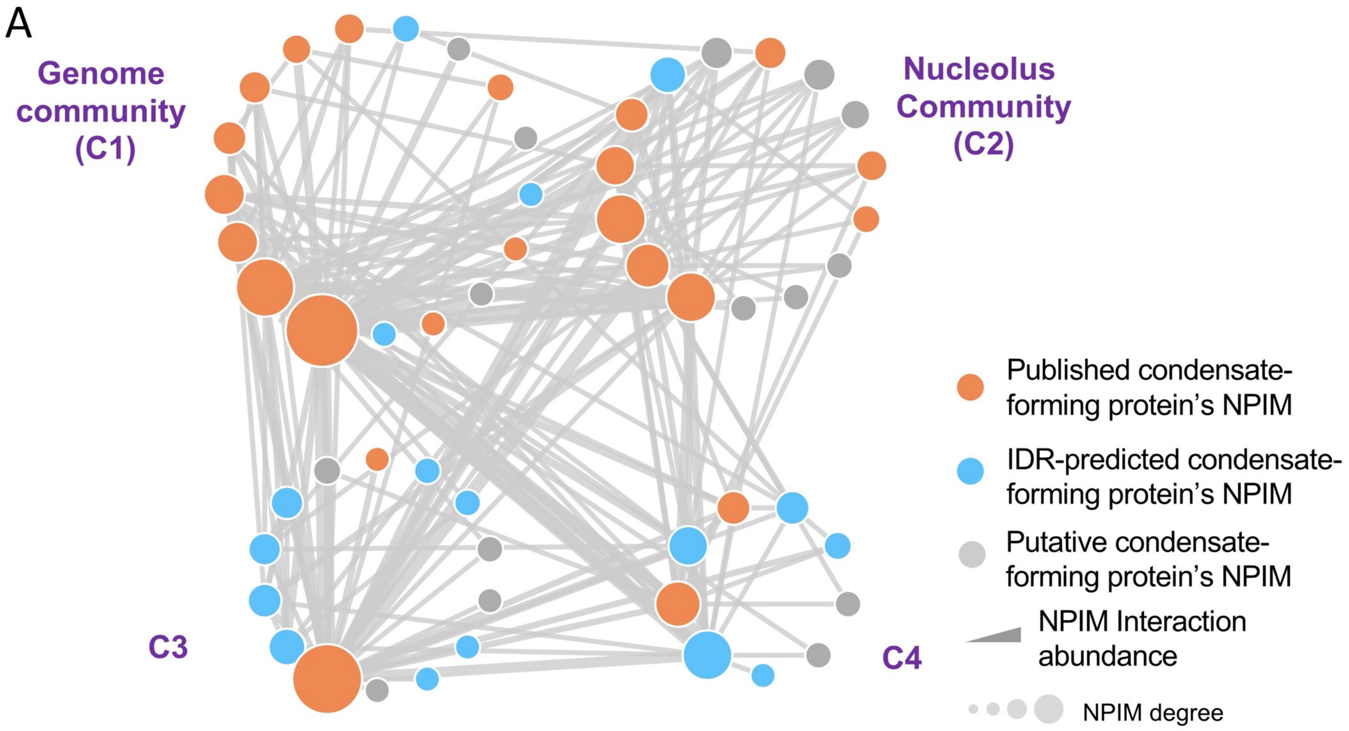


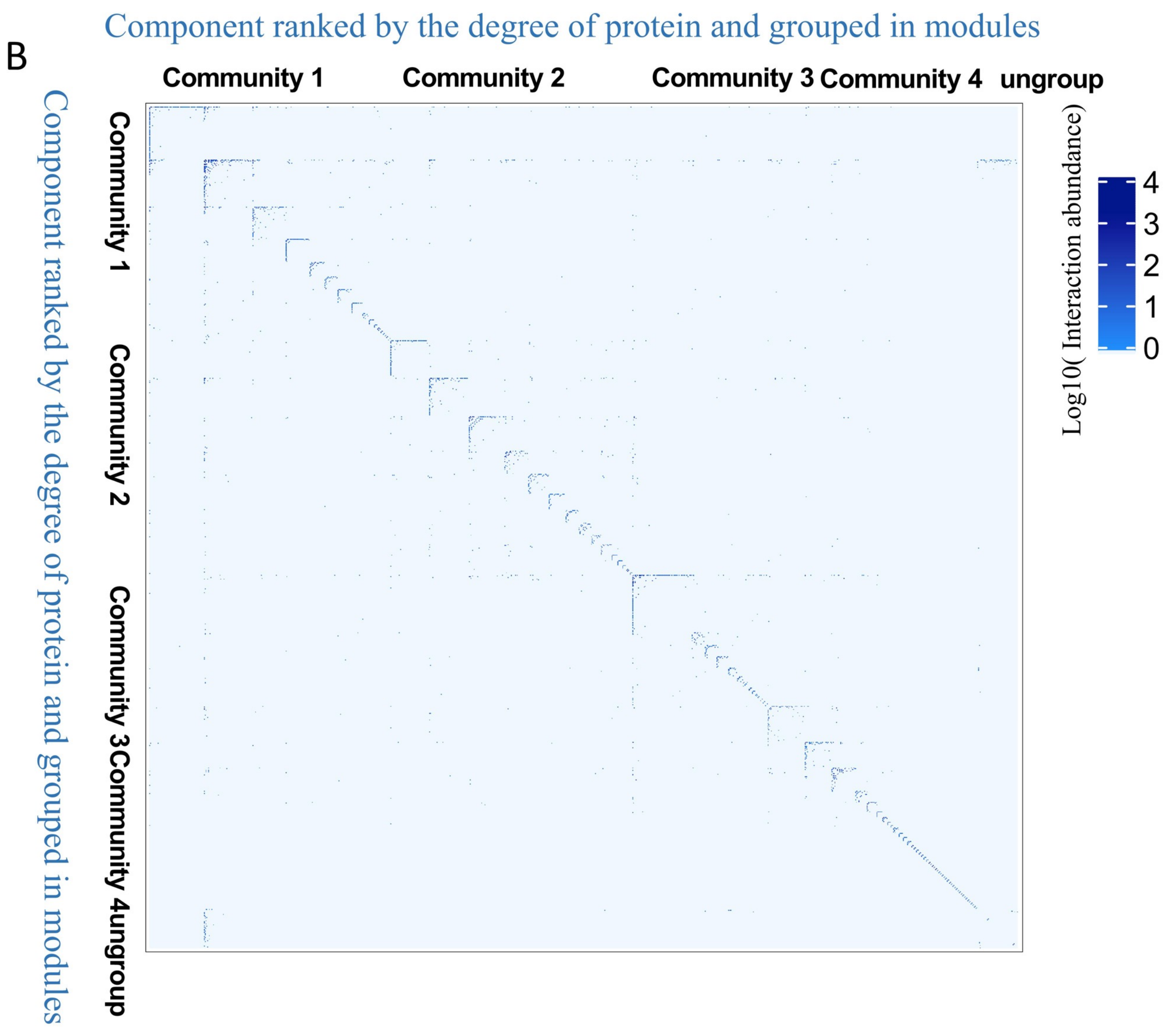

#### Supplemental Figure S15

#### The hierarchical organization of proteins into modules and communities in nucleome

(A) Diagram represents a system of 4 communities present in nucleome. The node and edge represents the NPIM and the interaction among NPIMs, which is defined as module-module interaction (MMI), within the nucleus graph, respectively. The higher order module of NPIMs is defined as community. The size of the node represents the degree of NPIM. The orange, light blue and light grey node represents the literature-annotated condensate-forming, the intrinsically disordered region (IDR)-containing and unknown putative condensate-forming NPIMs, respectively. The C1 and C2 community is defined as the genome community and nucleolus community, respectively. Both C3 and C4 communities represent NPIMs that might locate within interchromatin spaces. The thickness of line represents the abundance of interaction between two NPIMs, which is a count of PPIs between these two NPIMs (**Table S4c**).

(B) Heatmap of inter- and intra-crosslinks (or PSM counts of XL-peptides) were generated from 1052 nuclear proteins (Table S1g-h). Nuclear protein's PPI data were grouped into 95 NPIMs, which were further organized into 4 distinct communities using MONET toolbox. The color intensity of the dot represents the abundance of protein interaction (PSMs of XL-peptides). Protein components are ranked based on the degree of proteins within each module. (**Table S4c**).

### Supplemental Figure S16

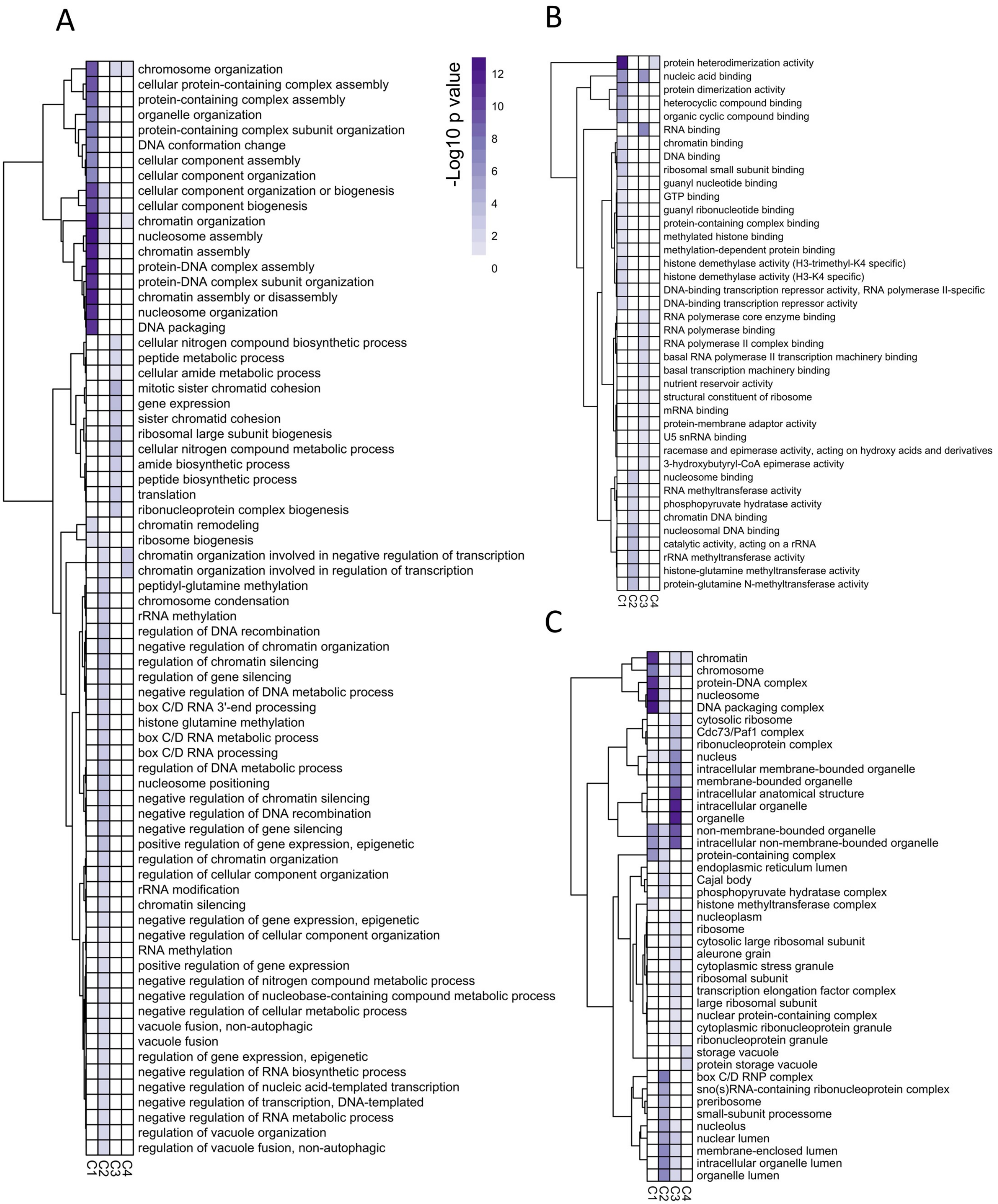

#### Supplemental Figure S16

#### GO analysis of four communities

(A – C) The heatmap shows the biological process (A), molecular function (B) and cellular component (C) analysis of four communities. The top 10% most interactive proteins (highest protein degree) in each community were used for the analysis. Row represents the name of GO term while column marks the different community of proteins. The color stands for the minus base 10 logarithm of *p* value (**Table S4d**).

**Supplemental Figure S17**

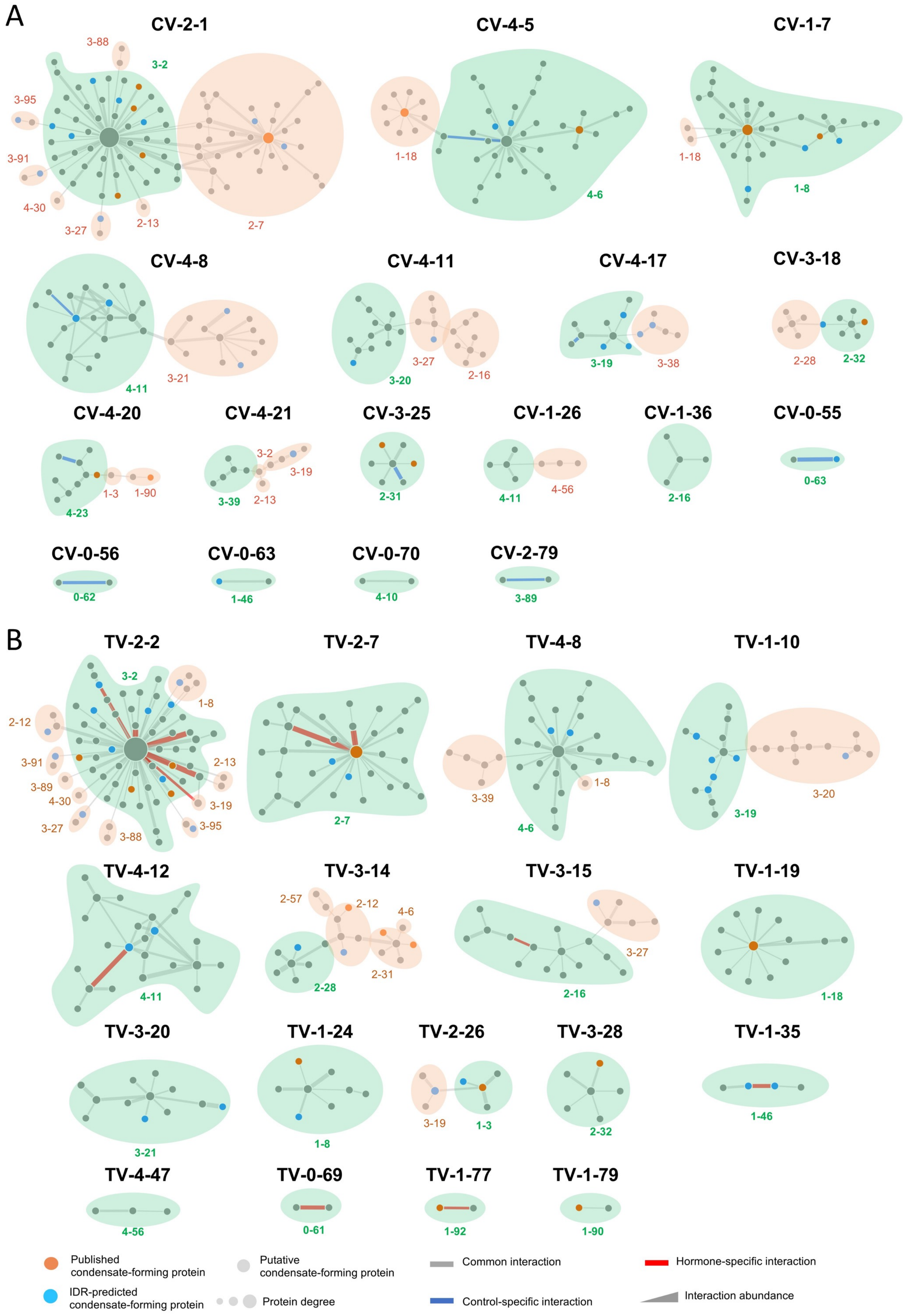

#### Supplemental Figure S17

#### Control- and Hormone-specific module variants

(A - B) Diagrams present Control-specific (A) and Hormone-specific (B) module variants. The letter marks the graph index of master module. The green and brown letter represents the scaffold and client graph index of master module, respectively. The light brown region denotes the components which belong to the same master module. The orange, light blue and light grey node represents the literature-annotated condensate-forming, the intrinsically disordered region (IDR)-containing and unknown putative condensate-forming proteins, respectively. The size of the nodes represents the protein degree. The blue, red and grey edges represent the Control-specific, Hormone-specific and common protein interaction, respectively (**Table S5e-g).**

**Supplemental Figure S18**

A

| Control-specific module variants | Master modules |
|---|---|
| *Module CV-2-1* | NPIM **3-2**, 2-7, 2-13, 3-27, 4-30, 3-88, 3-91, 3-95 |
| *Module CV-4-5* | NPIM **4-6**, 1-18 |
| *Module CV-1-7* | NPIM **1-8**, 1-18 |
| *Module CV-4-8* | NPIM **4-11**, 3-21 |
| *Module CV-4-11* | NPIM **3-20,** 2-16, 3-27 |
| *Module CV-4-17* | NPIM **3-19**, 3-38 |
| *Module CV-3-18* | NPIM **2-32**, 2-28 |
| *Module CV-4-20* | NPIM **4-23,** 1-3, 1-90 |
| *Module CV-4-21* | NPIM **3-39,** 3-2, 2-13, 3-19 |
| *Module CV-3-25* | NPIM **2-31** |
| *Module CV-1-26* | NPIM **4-11**, 4-56 |
| *Module CV-1-36* | NPIM **2-16** |
| *Module CV-0-55* | NPIM **0-63** |
| *Module CV-0-56* | NPIM **0-62** |
| *Module CV-0-63* | NPIM **1-46** |
| *Module CV-0-70* | NPIM **4-10** |
| *Module CV-2-79* | NPIM **3-89** |

B

| Hormone-specific module variants | Master modules |
|---|---|
| *Module TV-2-2* | NPIM **3-2**, 1-8, 2-12, 2-13, 3-19, 3-27, 4-30, 3-88, 3-89, 3-91, 3-95 |
| *Module TV-2-7* | NPIM **2-7** |
| *Module TV-4-8* | NPIM **4-6**, 1-8, 3-39 |
| *Module TV-1-10* | NPIM **3-19**, 3-20 |
| *Module TV-4-12* | NPIM **4-11** |
| *Module TV-3-14* | NPIM **2-28**, 4-6, 2-12, 2-31, 2-57 |
| *Module TV-3-15* | NPIM **2-16**, 3-27 |
| *Module TV-1-19* | NPIM **1-18** |
| *Module TV-3-20* | NPIM **3-21** |
| *Module TV-1-24* | NPIM **1-8** |
| *Module TV-2-26* | NPIM **1-3**, 3-19 |
| *Module TV-3-28* | NPIM **2-32** |
| *Module TV-1-35* | NPIM **1-46** |
| *Module TV-4-47* | NPIM **4-56** |
| *Module TV-0-69* | NPIM **0-61** |
| *Module TV-1-77* | NPIM **1-92** |
| *Module TV-1-79* | NPIM **1-90** |

Master module means that the integrated whole module was constructed using both Control and Hormone-treated PPI data. CV, Control module variant; TV, Treatment module variant.

#### Supplemental Figure S18

#### Relationship between the specific module variants and master modules

(A - B) Tables contain relationship between the Control-specific module variants (A) and master modules as well as that between Hormone-specific module variants (B) and master modules. The purple and brown graph index represents the corresponding scaffold and client master module, respectively, of the specific module variants (**Table S5e-g).**

### Supplemental Figure S19

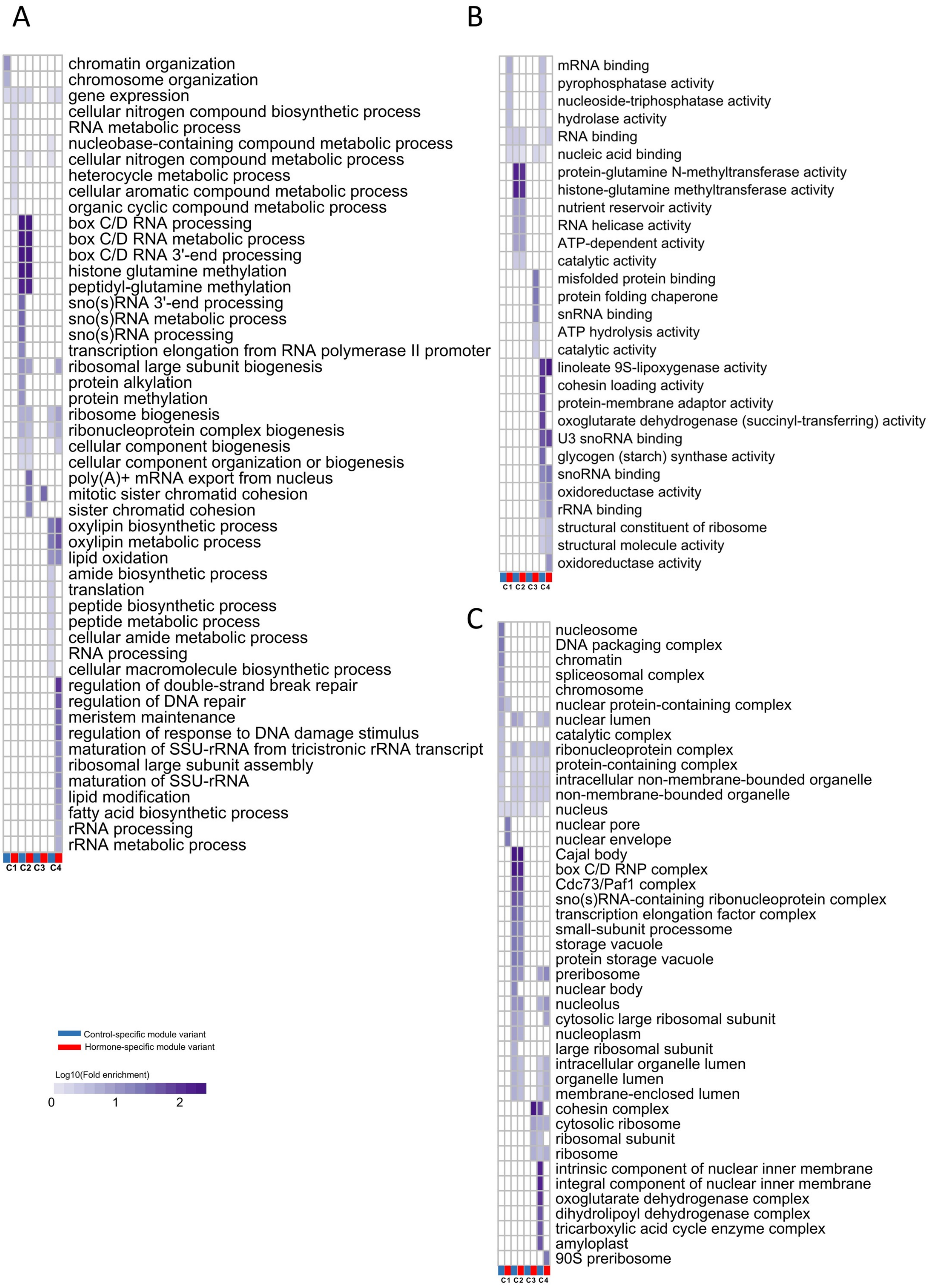

#### Supplemental Figure S19

#### GO comparison of Control- and Hormone-specific module variants in each community

(A - C) The heatmap of biological process (A), molecular function (B) and cellular component (C) analysis of the proteins obtained from control- and hormone-specific module variants in each community. Row represents the name of GO term while column marks the different group of proteins. The purple color represents the base 10 logarithm of fold enrichment of each GO term. The dark blue and red rectangles annotate the group of Control- and Hormone-specific modules, respectively. C1, C2, C3 and C4 represents the community 1, 2, 3 and 4, respectively (**Table S5h**)

**Supplemental Figure S20**

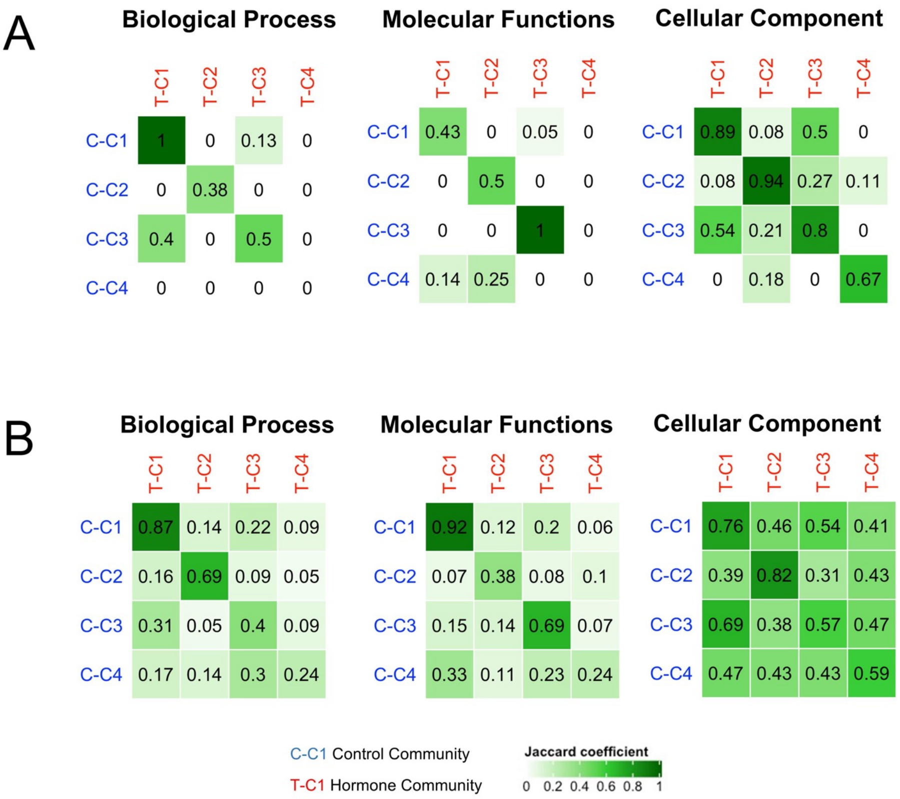


A — Biological Process

| | T-C1 | T-C2 | T-C3 | T-C4 |
|---|---|---|---|---|
| C-C1 | 1 | 0 | 0.13 | 0 |
| C-C2 | 0 | 0.38 | 0 | 0 |
| C-C3 | 0.4 | 0 | 0.5 | 0 |
| C-C4 | 0 | 0 | 0 | 0 |

A — Molecular Functions

| | T-C1 | T-C2 | T-C3 | T-C4 |
|---|---|---|---|---|
| C-C1 | 0.43 | 0 | 0.05 | 0 |
| C-C2 | 0 | 0.5 | 0 | 0 |
| C-C3 | 0 | 0 | 1 | 0 |
| C-C4 | 0.14 | 0.25 | 0 | 0 |

A — Cellular Component

| | T-C1 | T-C2 | T-C3 | T-C4 |
|---|---|---|---|---|
| C-C1 | 0.89 | 0.08 | 0.5 | 0 |
| C-C2 | 0.08 | 0.94 | 0.27 | 0.11 |
| C-C3 | 0.54 | 0.21 | 0.8 | 0 |
| C-C4 | 0 | 0.18 | 0 | 0.67 |

B — Biological Process

| | T-C1 | T-C2 | T-C3 | T-C4 |
|---|---|---|---|---|
| C-C1 | 0.87 | 0.14 | 0.22 | 0.09 |
| C-C2 | 0.16 | 0.69 | 0.09 | 0.05 |
| C-C3 | 0.31 | 0.05 | 0.4 | 0.09 |
| C-C4 | 0.17 | 0.14 | 0.3 | 0.24 |

B — Molecular Functions

| | T-C1 | T-C2 | T-C3 | T-C4 |
|---|---|---|---|---|
| C-C1 | 0.92 | 0.12 | 0.2 | 0.06 |
| C-C2 | 0.07 | 0.38 | 0.08 | 0.1 |
| C-C3 | 0.15 | 0.14 | 0.69 | 0.07 |
| C-C4 | 0.33 | 0.11 | 0.23 | 0.24 |

B — Cellular Component

| | T-C1 | T-C2 | T-C3 | T-C4 |
|---|---|---|---|---|
| C-C1 | 0.76 | 0.46 | 0.54 | 0.41 |
| C-C2 | 0.39 | 0.82 | 0.31 | 0.43 |
| C-C3 | 0.69 | 0.38 | 0.57 | 0.47 |
| C-C4 | 0.47 | 0.43 | 0.43 | 0.59 |

#### Supplemental Figure S20

#### GO comparison of Control and Hormone communities

(A - B) The heatmap shows the comparison of biological process (left panel), molecular function (middle panel) and cellular component (right panel) analysis of proteins derived from both Control (rows; blue letters) and Hormone (columns; red letters) communities. The top 10% (A) and 50% (B) most interactive proteins (highest protein degree) in each community were used for the analysis. The similarity among the communities is evaluated by Jaccard Coefficient (JC) of related GO terms. The green color of the heatmap stands for the level of Jaccard Coefficient. C1, C2, C3 and C4 represents the community 1, 2, 3 and 4, respectively. Ungrouped are those modules that fail to be integrated into a community. The prefixes of communities, “C-” and “T-”, stand for Control and Hormone dataset, respectively.  (Table S5i-l).

#### Supplemental Figure S21

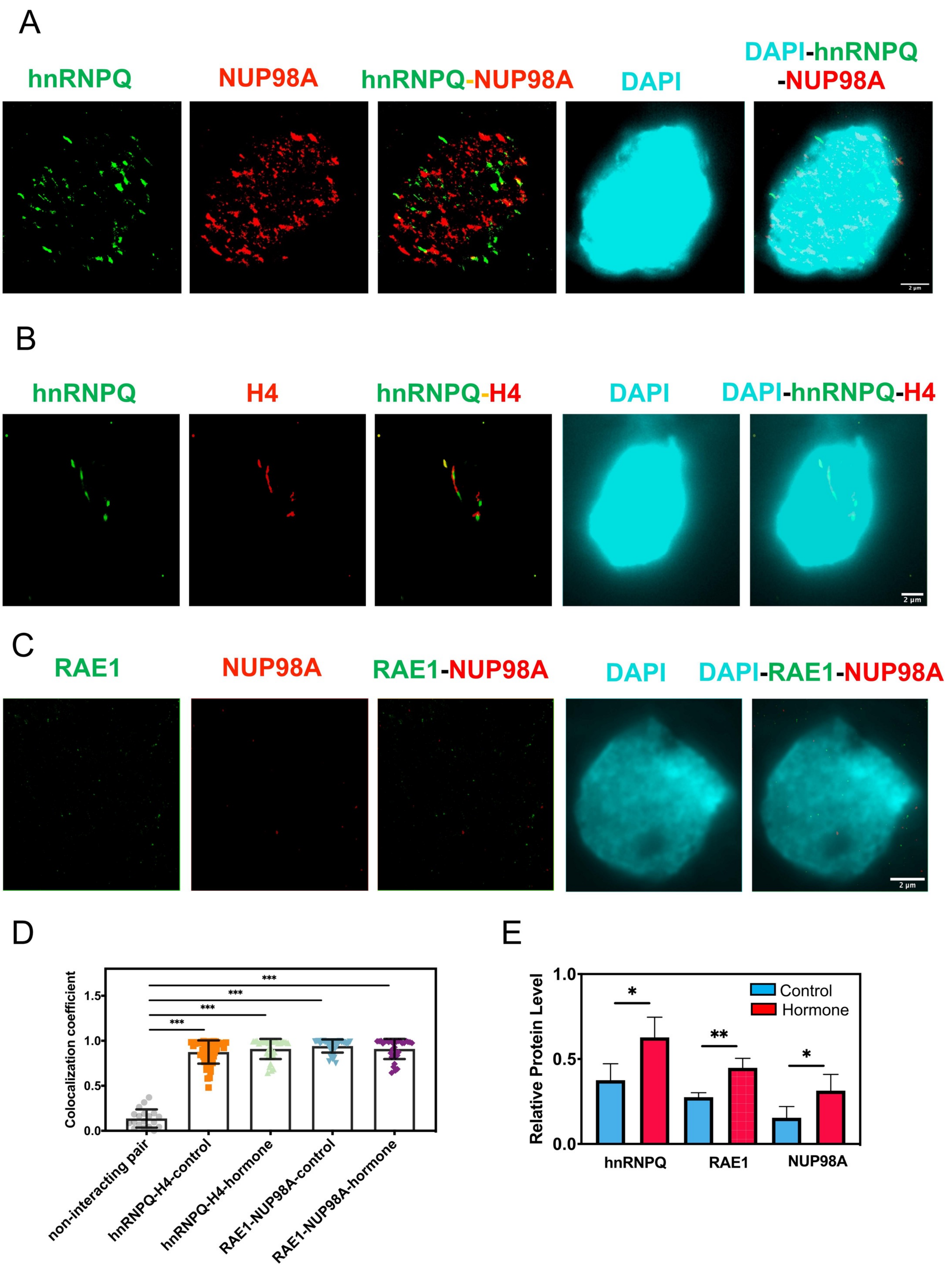

#### Supplemental Figure S21

(A) STORM super-resolution imaging of a non-interacting protein pair of hnRNPQ with NUP98A. HnRNPQ and NUP98A protein is shown as green (from Alexa Fluor 568 dye) and red color (from Alexa Fluor 750 dye), respectively. The colocalized proteins are shown in yellow color. DAPI-stained nuclei are marked in cyan color.

(B - C) STORM super-resolution imaging of PPIs of hnRNPQ-histone H4 pair and RAE1-NUP98A pair with 1:10 antigen peptides-titration. Both hnRNPQ and RAE1 proteins are shown as green color (from Alexa Fluor 568 dye), whereas H4 and NUP98A proteins shown as red color (from Alexa Fluor 750 dye). The colocalized proteins are shown in yellow color. DAPI-stained nuclei are marked in cyan color.

(D) Bar chart of the colocalization coefficients (MOC, Manders' Overlap Coefficient) of PPIs, hnRNPQ-H4 protein pair and RAE1-NUP98A protein pair, under both non-hormone treatment control and hormone treatment. The STORM super-resolution imaging data of the non-interacting protein pair of hnRNPQ-NUP98A proteins were used as the PPI control. Average values and error bars (±SEM) are shown. n.s., *, **, and *** as $p \geq 0.05$, $p < 0.05$, $p < 0.01$, and $p < 0.001$, respectively.

(E) Bar chart of comparison of the nuclear protein level of hnRNPQ, RAE1 and NUP98A under both control and hormone treatment. Average values and error bars (±SEM) are shown. n.s., *, **, and *** as $p \geq 0.05$, $p < 0.05$, $p < 0.01$, and $p < 0.001$, respectively.

**Supplemental Figure S22**

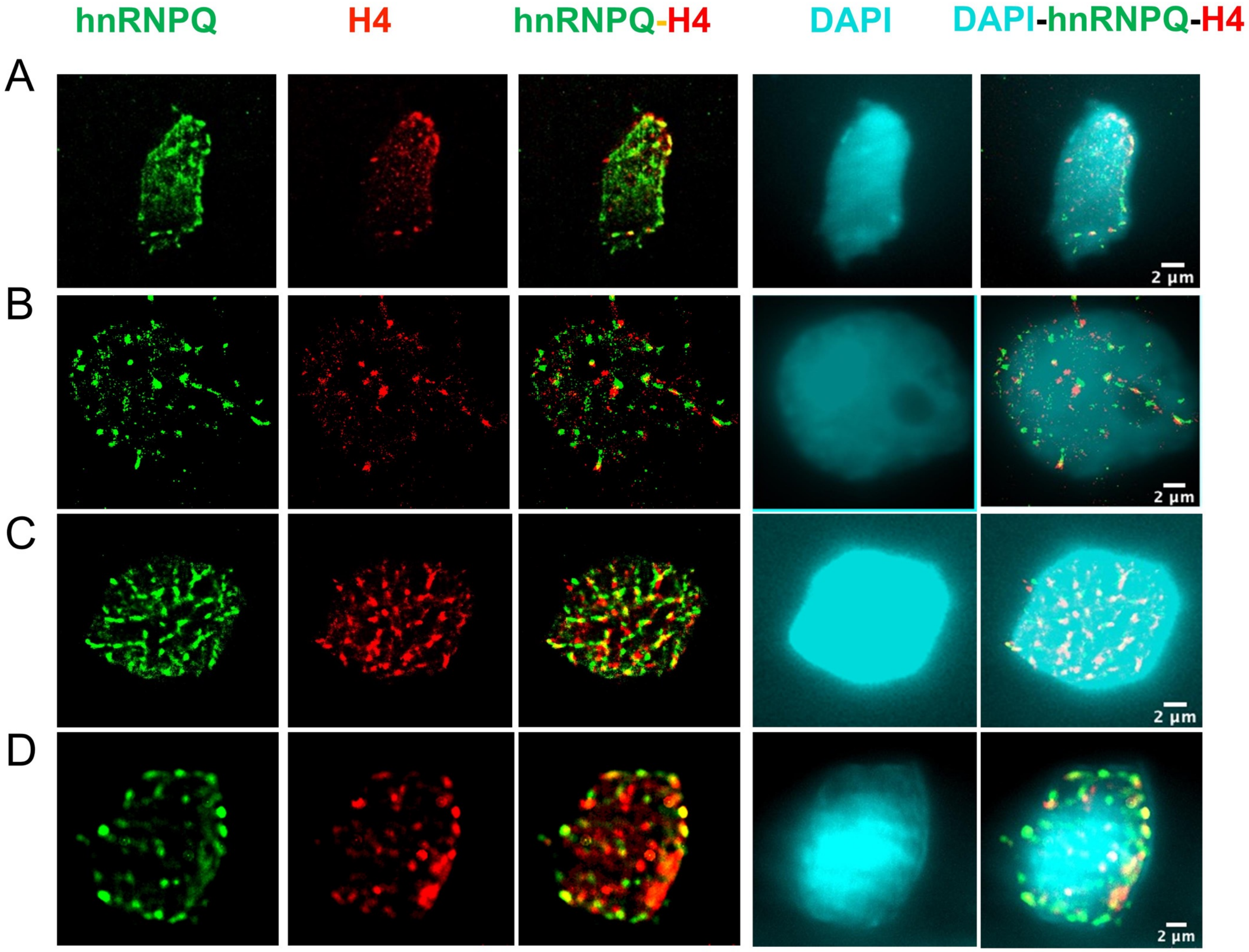

#### Supplemental Figure S22

#### Super-resolution microscope of protein hnRNPQ and histone H4 under control condition

(A - D) STORM super-resolution imaging of PPIs of hnRNPQ with histone H4 under non-hormone treatment control condition. HnRNPQ and histone H4 protein is shown as green (from Alexa Fluor 568 dye) and red color (from Alexa Fluor 750 dye), respectively. The colocalized protein pairs are shown in yellow color. DAPI-stained nuclei are marked in cyan color.

**Supplemental Figure S23**

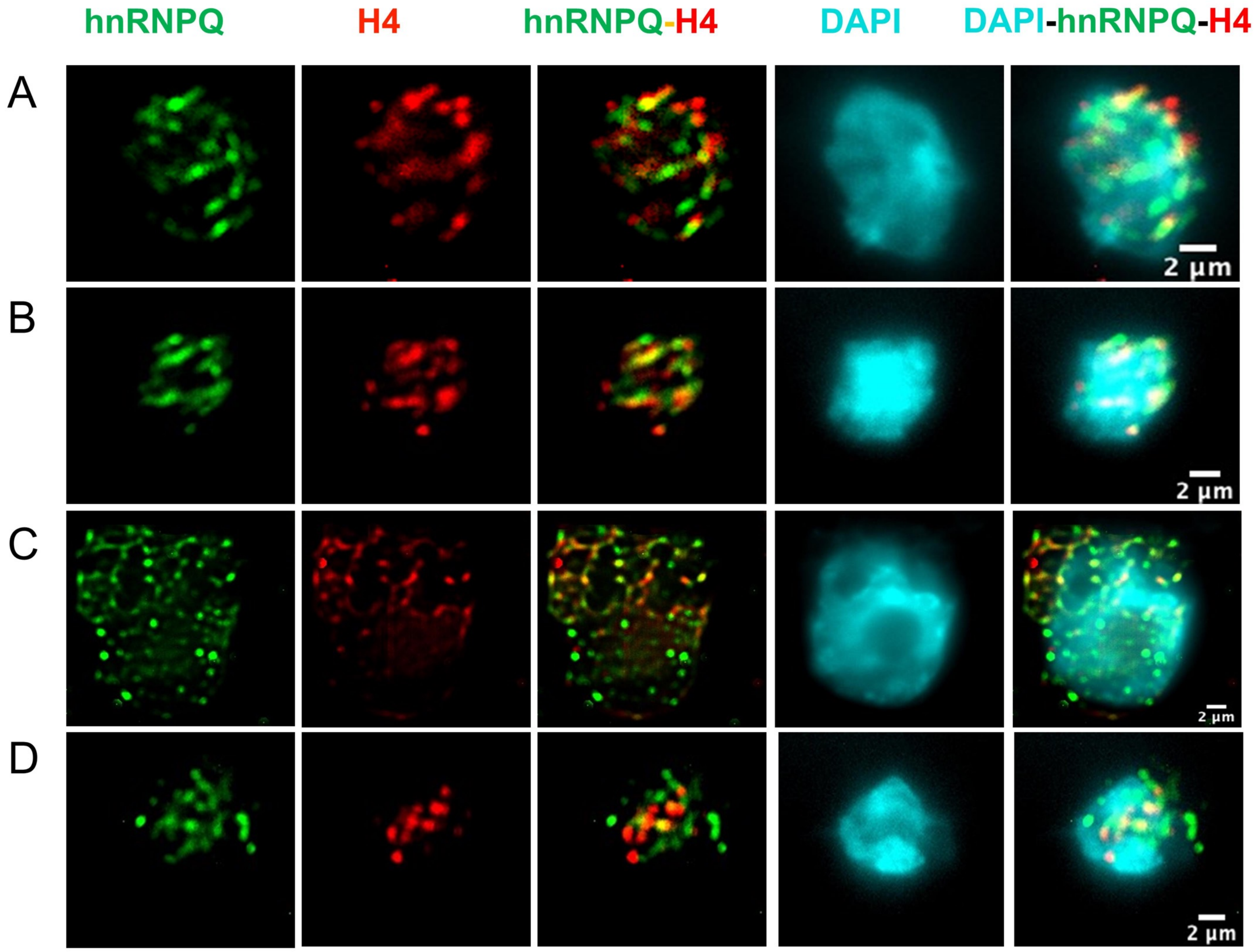

#### Supplemental Figure S23

#### Super-resolution microscope of protein hnRNPQ and histone H4 under hormone condition

(A - E) STORM super-resolution imaging of PPIs of hnRNPQ with histone H4 under hormone treatment condition. HnRNPQ and histone H4 protein is shown as green (from Alexa Fluor 568 dye) and red color (from Alexa Fluor 750 dye), respectively. The colocalized protein pairs are shown in yellow color. DAPI-stained nuclei are marked in cyan color.

**Supplemental Figure S24**

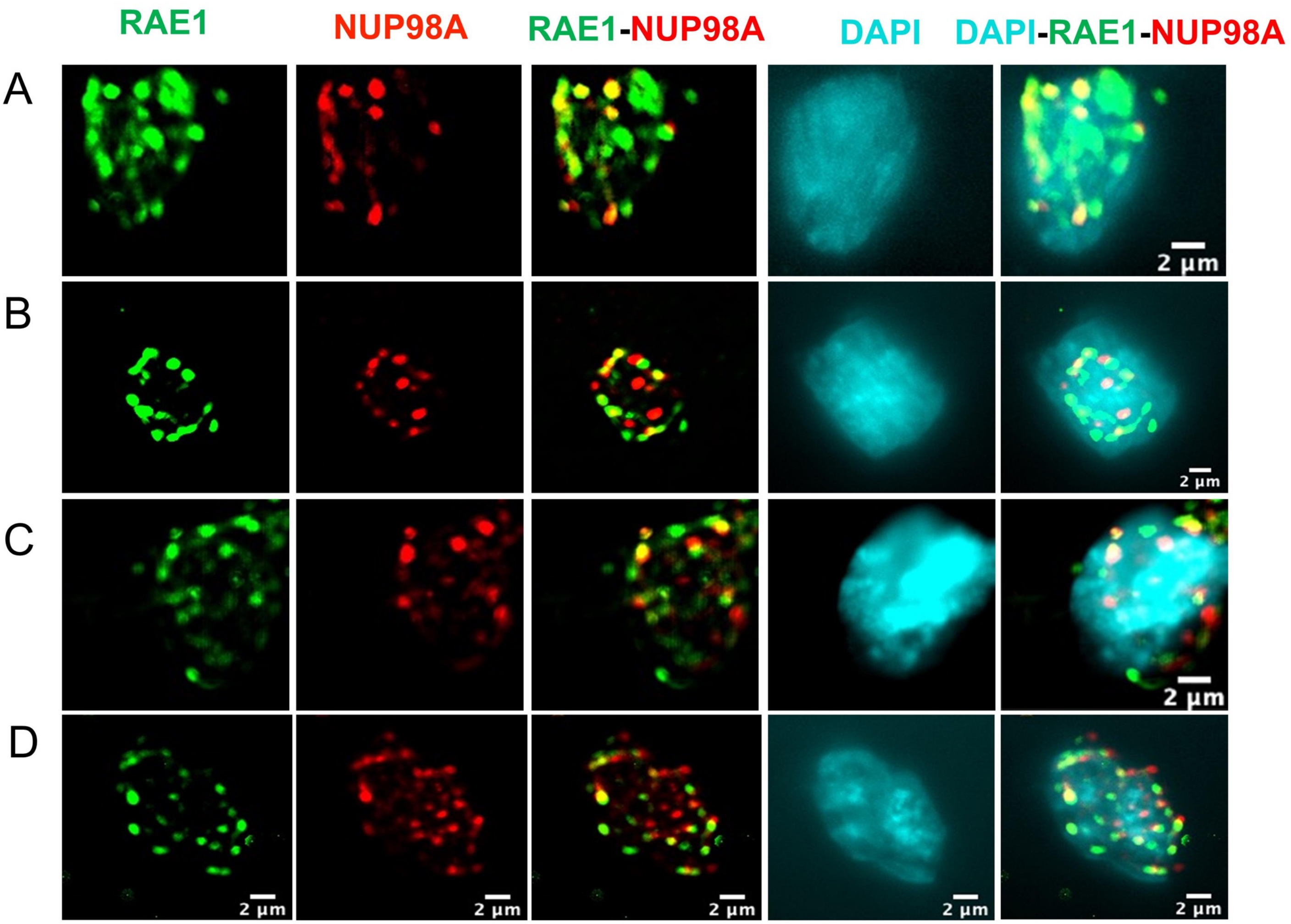

#### Supplemental Figure S24

#### Super-resolution microscope of protein RAE1 and NUP98A under control condition

(A - E) STORM super-resolution imaging of PPIs of RAE1 with NUP98A under non-hormone treatment control condition. RAE1 and NUP98A protein is shown as green (from Alexa Fluor 568 dye) and red color (from Alexa Fluor 750 dye), respectively. The colocalized protein pairs are shown in yellow color. DAPI-stained nuclei are marked in cyan color.

**Supplemental Figure S25**

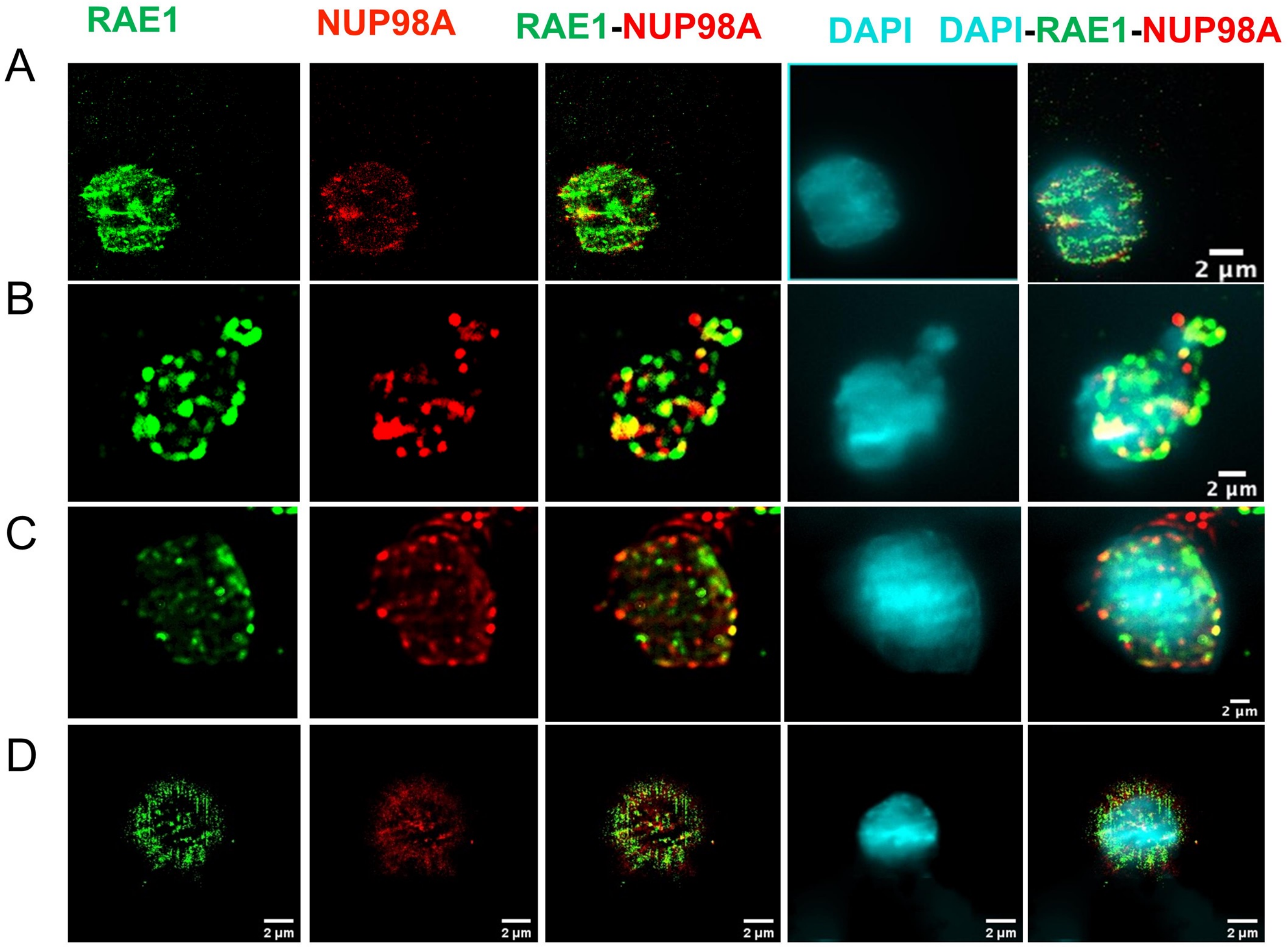

#### Supplemental Figure S25

#### Super-resolution microscope of protein RAE1 and NUP98A under hormone condition

(A - E) STORM super-resolution imaging of PPIs of RAE1 with NUP98A under hormone treatment condition. RAE1 and NUP98A protein is shown as green (from Alexa Fluor 568 dye) and red color (from Alexa Fluor 750 dye), respectively. The colocalized protein pairs are shown in yellow color. DAPI-stained nuclei are marked in cyan color.

#### Supplemental Figure S26

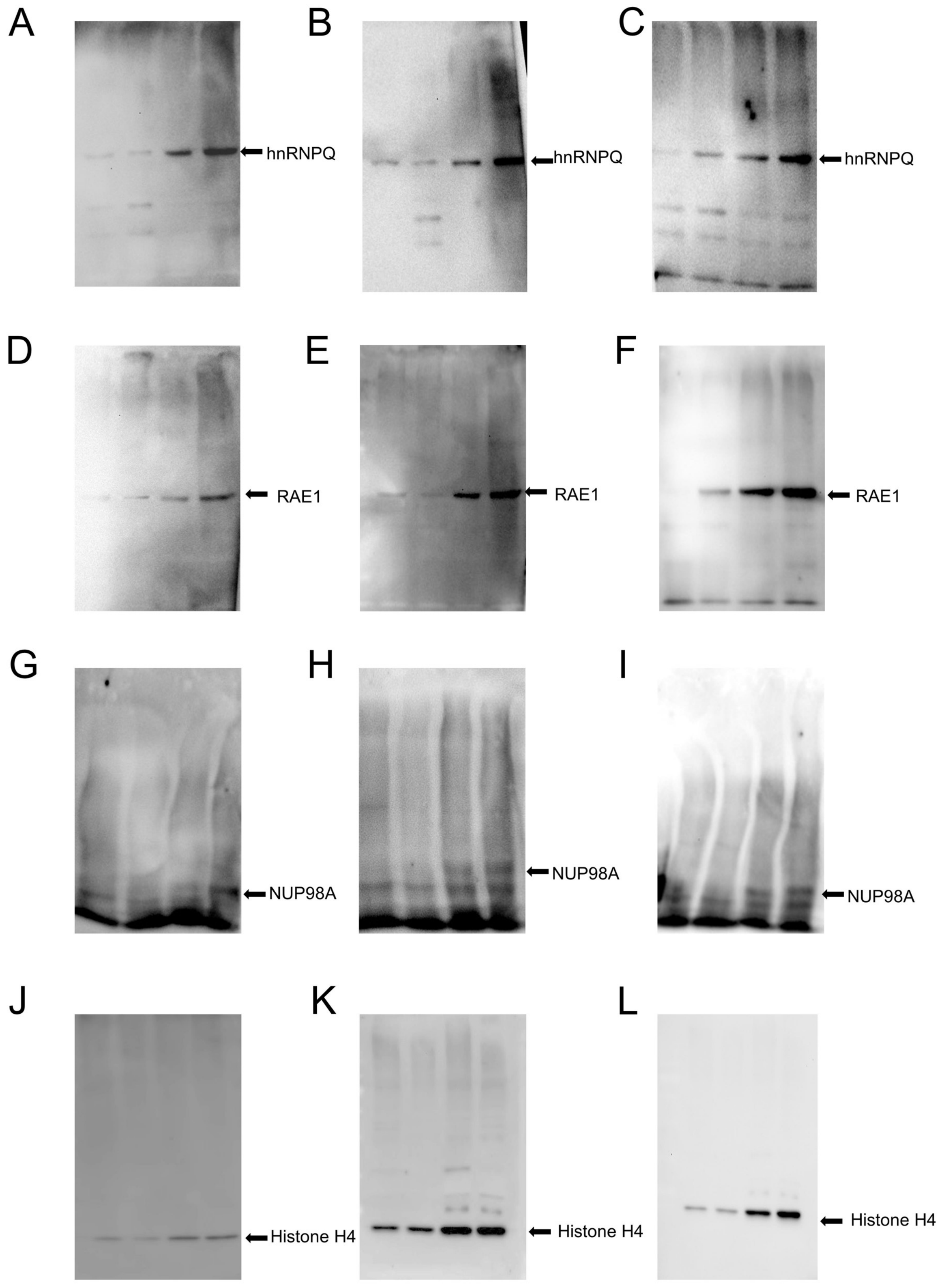

#### Supplemental Figure S26

#### Western blot analysis of the selected nuclear proteins

(A - L) Western blot analysis of nuclear proteins extracted from nuclei of both non-hormone treated control and hormone-treated tissues. Protein hnRNPQ, H4, RAE1 and NUP98A was identified using anti-hnRNPQ, -H4, -RAE1 and -NUP98A polyclonal antibodies, respectively.

**Supplemental Figure S27**

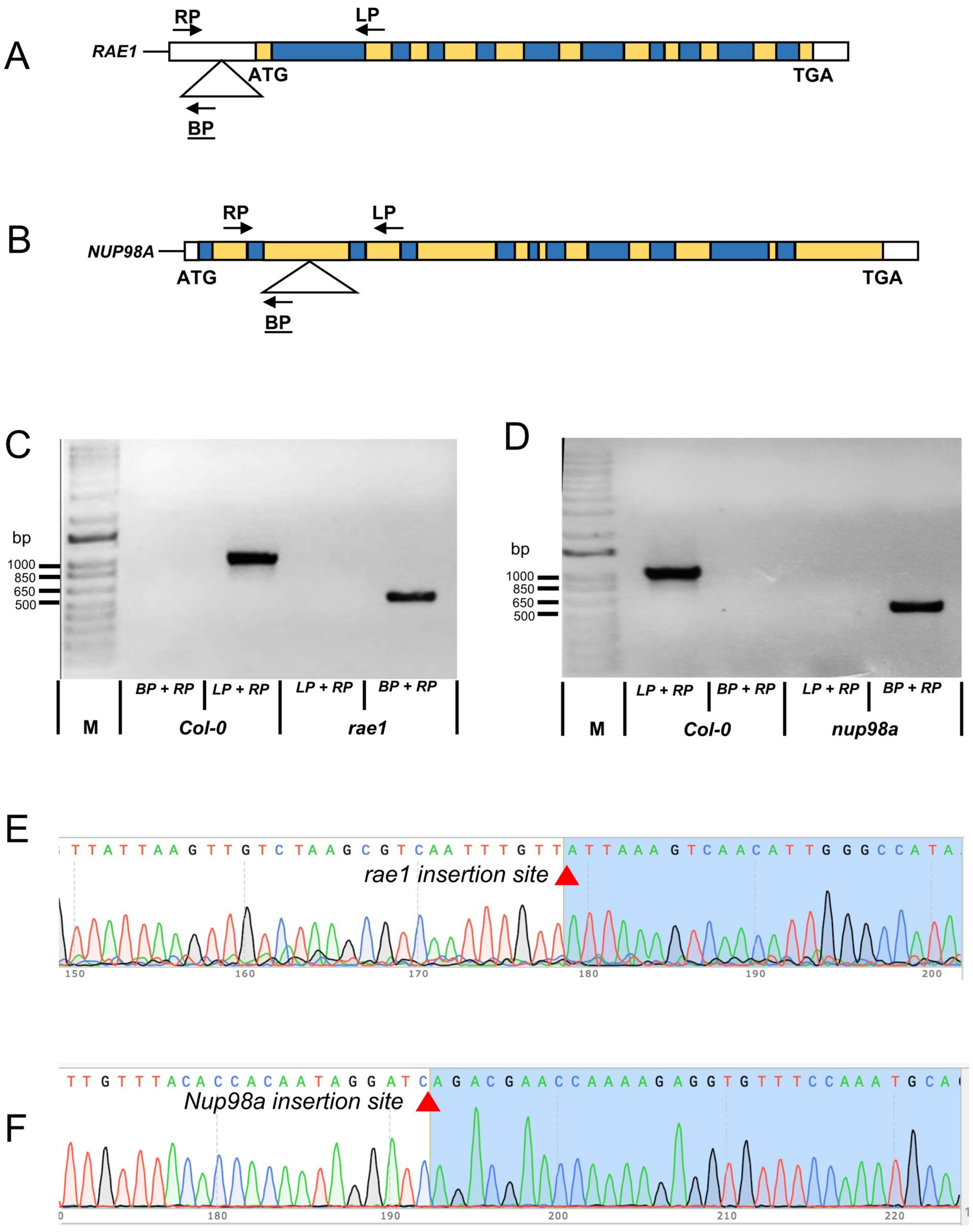

#### Supplemental Figure S27

#### Genotyping of T-DNA insertional mutants

(A - B) Schematic diagram of RAE1 (A) and NUP98A (B) gene structure. Triangles represent T-DNA insertion sites in mutants. Primers used for genotyping detection are represented by arrows. LP, RP and BP arrow stands for left genomic, right genomic and T-DNA border primer, respectively. White, yellow and blue box with black border marks the 5' or 3' UTR, exon and intron region, respectively. ATG and TGA indicates the start and stop codon, respectively.

(C - D) Genotyping of *rae1* (C) and *nup98a* (D) mutants by PCR. Primer pairs used to amplify fragment are BP + RP and LP + RP, respectively. The M stands for the DNA maker.

(E - F) DNA sequencing verification of T-DNA insertion sites in *rae1* (E) and *nup98a* (F) mutants. The insertion site is indicated with red triangles.

**Supplemental Figure S28**

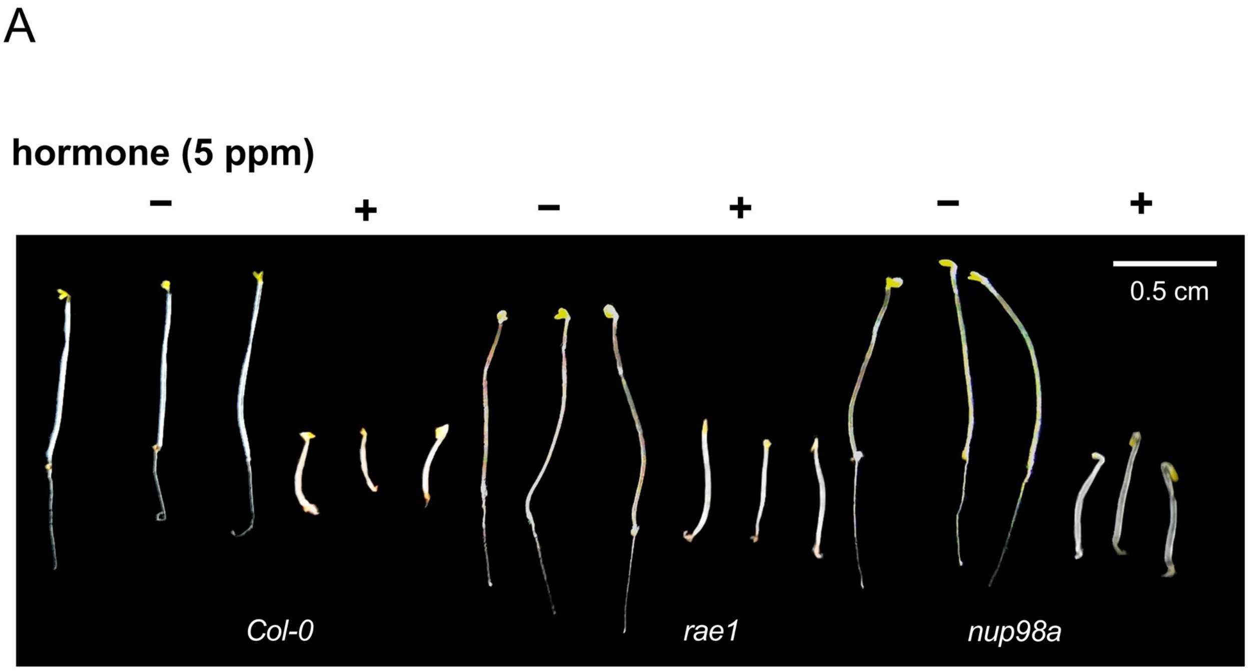

#### Supplemental Figure S28

#### Triple response assay of *rae1* and *nup98a* mutants

Triple response phenotypes of the loss-of-function mutants, *rae1* and *nup98a*, of the model plant Arabidopsis. Representative phenotypes of the etiolated seedlings of 4-day-old *Col-0*, *rae1* and *nup98a* Arabidopsis grown without ( - ) and with hormone treatment ( + ).

### Supplemental Figure S30

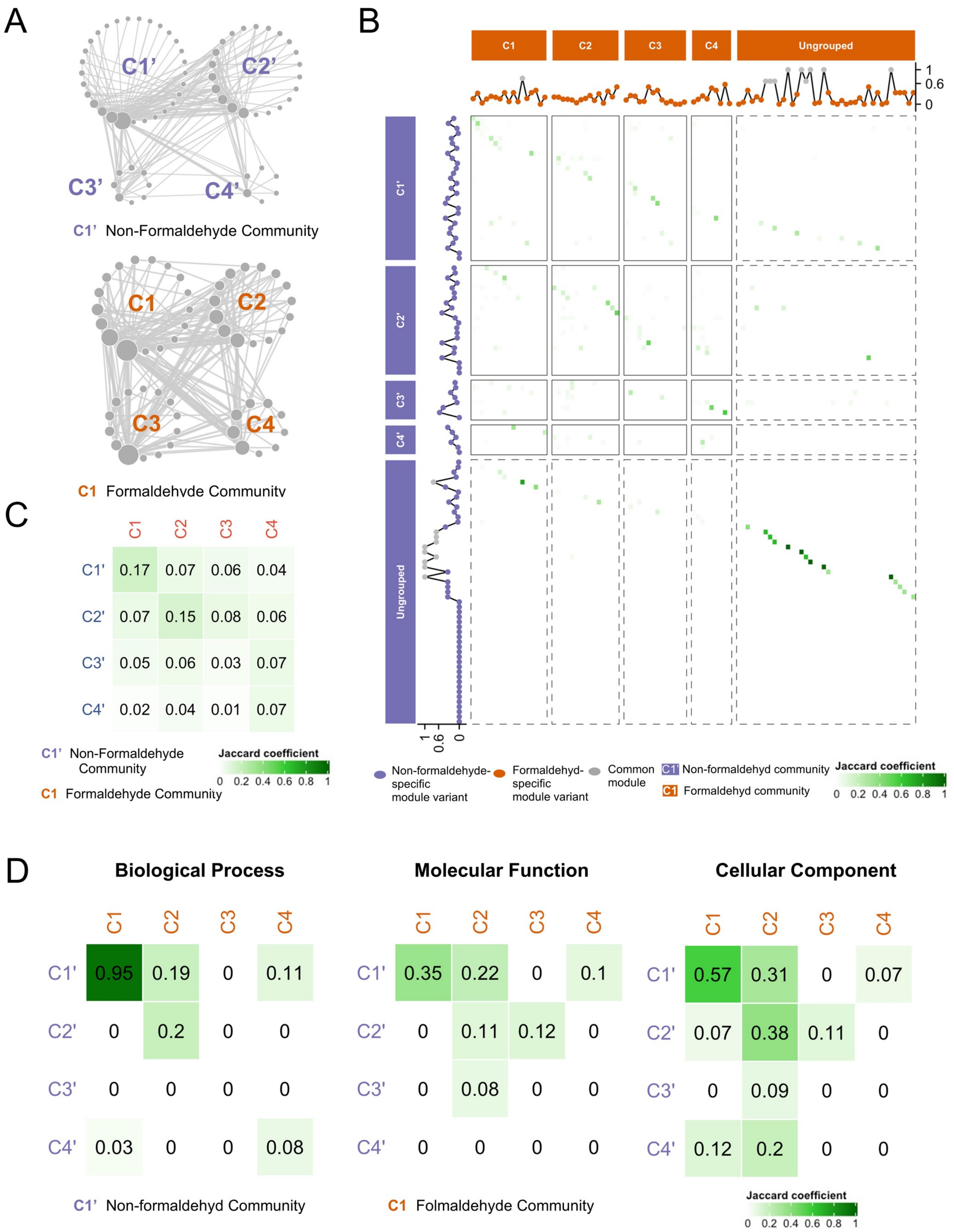

#### Supplemental Figure S30

**Heatmap and GO analysis of modules and communities generated by PPIs derived from non-formaldehyde-crosslinked and formaldehyde-crosslinked XL-peptides**

(A) The schematic topological graph represents communities of non-formaldehyde-crosslinked data (generated by 1,398 Arabidopsis ortholog PPIs from non-formaldehyde-crosslinked XL-peptides, 997 Arabidopsis ortholog proteins; left panel; violet) and formaldehyde-crosslinked data (generated by 1,211 Arabidopsis ortholog PPIs from formaldehyde-crosslinked XL-peptides, 877 Arabidopsis ortholog proteins; right panel; orange). The node and edge represents the NPIM (or variant of NPIM) and the interaction among modules (MMI, module-module interaction) within the nucleus graph, respectively. The higher order module of modules is defined as community. The size of the node represents the degree of the module. The thickness of the line represents the abundance of module-module interaction, which is the count of PPIs between these two modules. Non-formaldehyde and formaldehyde stands for non-formaldehyde-crosslinked and formaldehyde-crosslinked data, respectively. The data are in Table S8a-d.

(B) The heatmap represents the module comparison between non-formaldehyde-crosslinked data (generated by 1,398 Arabidopsis ortholog PPIs from non-formaldehyde-crosslinked XL-peptides; 997 Arabidopsis ortholog proteins; rows; violet) and formaldehyde-crosslinked data (generated by 1,211 Arabidopsis ortholog PPIs from formaldehyde-crosslinked XL-peptides, 877 Arabidopsis ortholog proteins; columns; orange). The similarity between two modules is evaluated by Jaccard Coefficient (JC), which is determined by the number of identical components divided by the total number of unique components. The green color of the heatmap stands for the level of Jaccard Coefficient. The violet and orange dots stand for the non-formaldehyde-crosslinked-specific module variants (JC <0.6), formaldehyde-crooslinked-specific module variants (JC<0.6), and common modules (JC>0.6), respectively (Table S8e). C1, C2, C3 and C4 represents the community 1, 2, 3, and 4, respectively. Ungrouped are those modules that fail to be integrated into a community.

(C) The heatmap represents the community comparison between non-formaldehyde-crosslinked (rows; violet letters) and formaldehyde-crosslinked (columns; orange letters) communities. The similarity between two modules is evaluated by Jaccard Coefficient (JC), which is determined by the number of identical components divided by the total number of unique components. The green color of the heatmap stands for the level of Jaccard Coefficient. C1, C2, C3 and C4 represents the community 1, 2, 3 and 4, respectively. The data are in Table S8c-d.

(D) The heatmap shows the comparison of biological process (left panel), molecular function (middle panel), and cellular component (right panel) analysis of proteins derived from non-formaldehyde-crosslinked data (rows; violet letters) and formaldehyde-crosslinked data (columns; orange letters) communities. The top 10% of most interactive proteins (highest protein degree) in each community were used for the analysis. The similarity among the communities is evaluated by the Jaccard Coefficient (JC) of related GO terms. The green color of the heatmap stands for the level of the Jaccard Coefficient (Table S8f-g).

**Supplemental Figure S31**

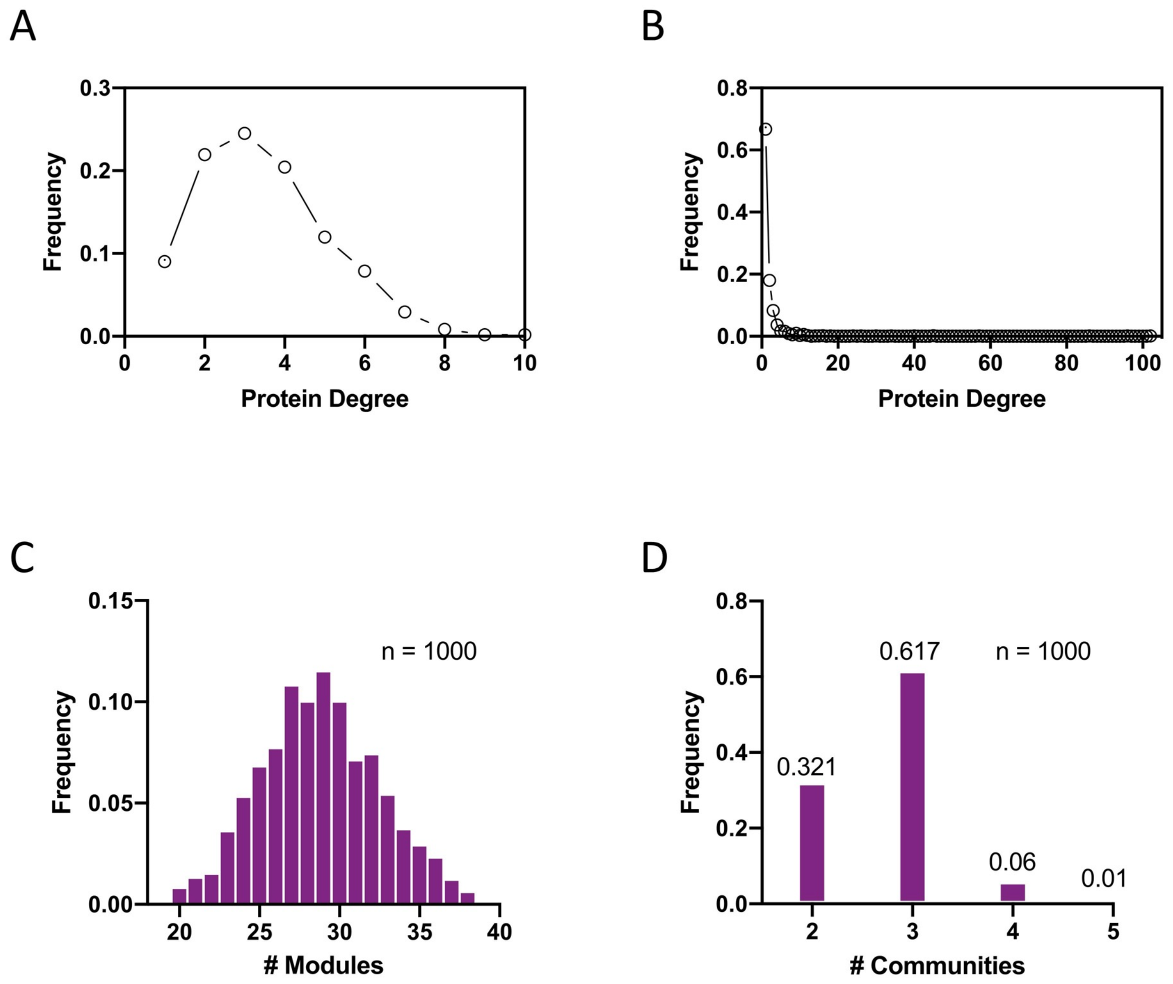

#### Supplemental Figure S31

#### Modules and communities from randomly generated PPIs by Erdös-Rényi random network

(A) Line chart shows the protein degree distribution based on random PPI network. The frequency stands for the proportion of proteins with a specific number of protein degree. The random PPI network is an Erdös-Rényi random network generated by equivalent number of nodes (1,052) and edges (1,297) with the nuclear PPI network in this study (**Table S1h**; Erdös and Rényi, 1959).

(B) Line chart shows the degree distribution of proteins based on PPIs in this study (**Table S1h-j)**. The frequency stands for the proportion of proteins with a specific number of protein degree.

(C - D) Histograms show the distribution of number of modules (C) and communities (D) generated by 1,000 random tests. The frequency stands for the proportion of tests with a specific number of modules (C) or communities (D). The n indicates the 1000 random tests. The details of random tests are described in Star Methods.

**Supplemental Figure S32**

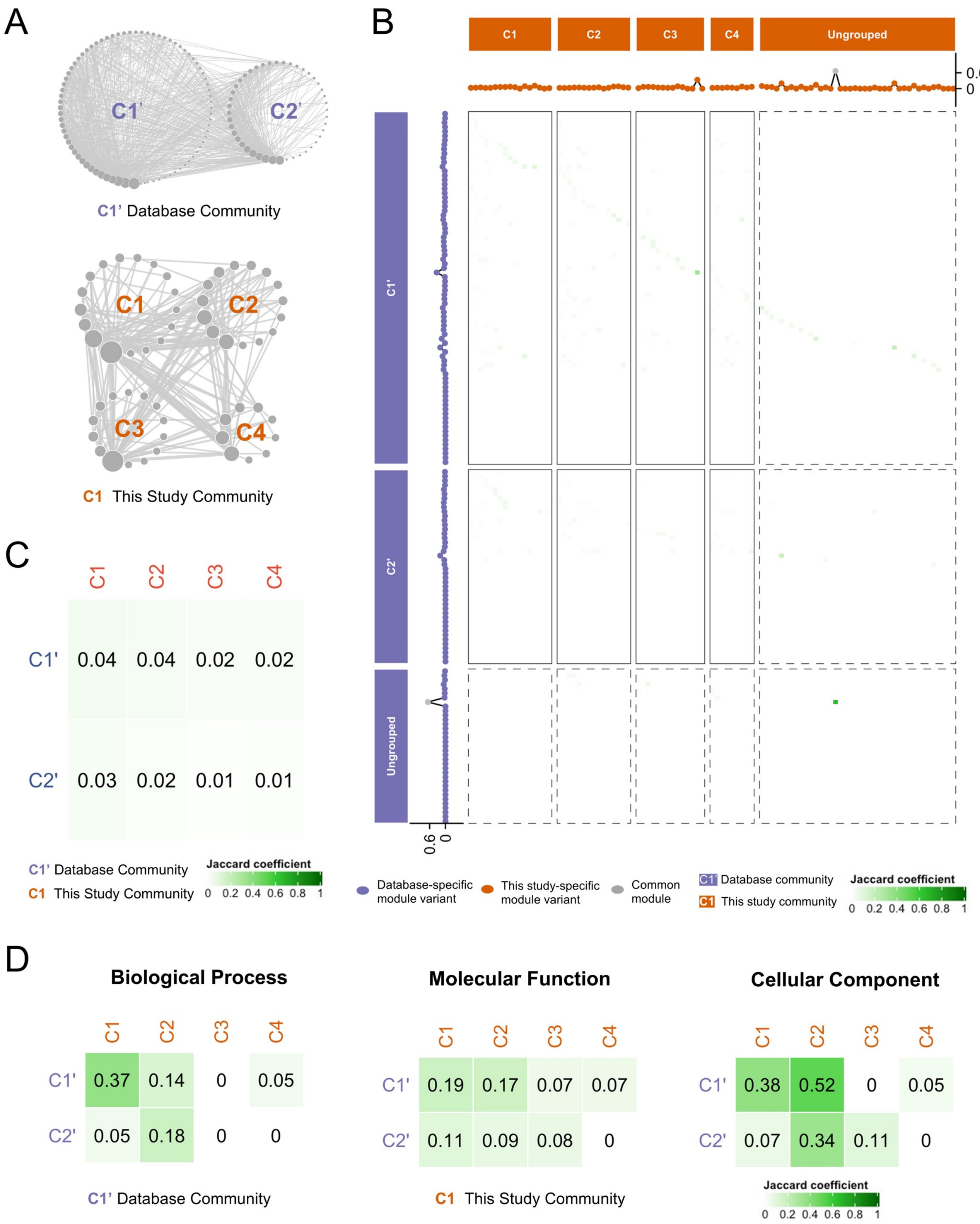

#### Supplemental Figure S32

**Heatmap and GO analysis of modules and communities generated by PPIs of database and this study**

(A) The schematic topological graph represents communities of database data (generated by 4,960 Arabidopsis PPIs from BioGrid database, 2,067 Arabidopsis proteins; left panel; violet) and FA-crosslinked data (generated by 1,211 Arabidopsis ortholog PPIs from this study, 877 Arabidopsis ortholog proteins; right panel; orange). The node and edge represents the NPIM (or variant of NPIM) and the interaction among modules (MMI, module-module interaction) within the nucleus graph, respectively. The higher order module of modules is defined as community. The size of the node represents the degree of the module. The thickness of the line represents the abundance of module-module interaction, which is the count of PPIs between these two modules. The data are in Table S9a-d.

(B) The heatmap represents the module comparison between database data (generated by 4,960 Arabidopsis PPIs from BioGrid database, 2,067 Arabidopsis proteins; rows; violet) and this study data (generated by 1,211 Arabidopsis ortholog PPIs from this study, 877 Arabidopsis ortholog proteins; columns; orange). The similarity between two modules is evaluated by Jaccard Coefficient (JC), which is determined by the number of identical components divided by the total number of unique components. The green color of the heatmap stands for the level of Jaccard Coefficient. The violet and orange dots stand for the database-specific module variants (JC <0.6), this study-specific module variants (JC<0.6), and common modules (JC>0.6), respectively (Table S9e). C1, C2, C3 and C4 represents the community 1, 2, 3, and 4, respectively. Ungrouped are those modules that fail to be integrated into a community.

(C) The heatmap represents the community comparison between the database (rows; violet letters) and this study (columns; orange letters) communities The similarity between two modules is evaluated by Jaccard Coefficient (JC), which is determined by the number of identical components divided by the total number of unique components. The green color of the heatmap stands for the level of Jaccard Coefficient. C1, C2, C3 and C4 represents the community 1, 2, 3 and 4, respectively. The data are in Table S9c-d.

(D) The heatmap shows the comparison of biological process (left panel), molecular function (middle panel), and cellular component (right panel) analysis of proteins derived from database (rows; violet letters) and this study (columns; orange letters) communities. The top 10% of most interactive proteins (highest protein degree) in each community were used for the analysis. The similarity among the communities is evaluated by the Jaccard Coefficient (JC) of related GO terms. The green color of the heatmap stands for the level of the Jaccard Coefficient (Table S9f-g).

#### Supplemental Figure S33

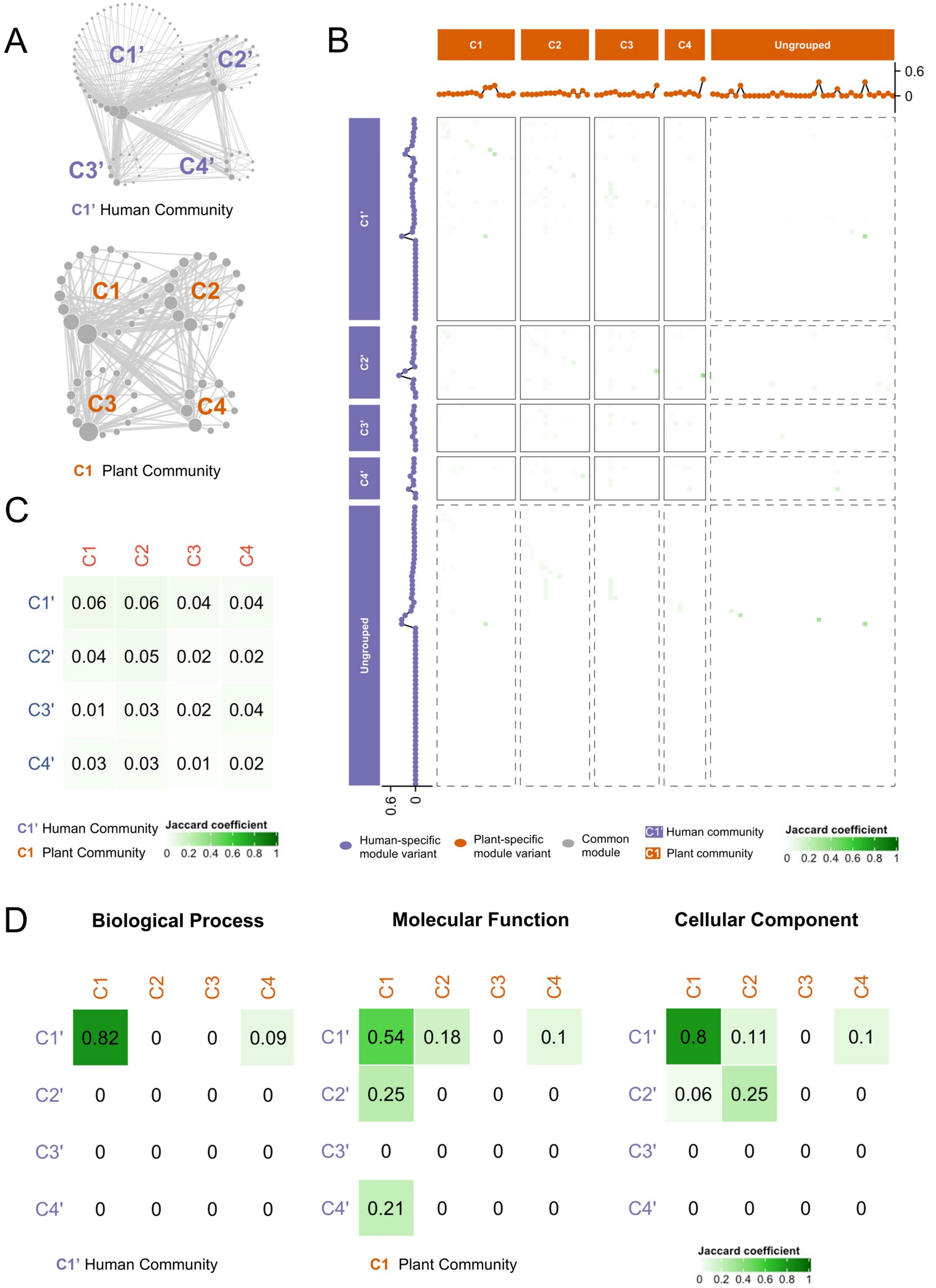

#### Supplemental Figure S33

**Heatmap and GO analysis of modules and communities generated by PPIs derived from human and plant nuclear XL-peptides**

(A) The schematic topological graph represents communities of human data (generated by 3,312 PPIs from human nuclear XL-peptides, 1580 proteins; left panel; violet) and plant data (generated by 1,211 Arabidopsis ortholog PPIs from plant nuclear XL-peptides, 877 Arabidopsis ortholog proteins; right panel; orange). The node and edge represents the NPIM (or variant of NPIM) and the interaction among modules (MMI, module-module interaction) within the nucleus graph, respectively. The higher order module of modules is defined as community. The size of the node represents the degree of the module. The thickness of the line represents the abundance of module-module interaction, which is the count of PPIs between these two modules. The data are in Table S10a-d.

(B) The heatmap represents the module comparison between human data (generated by 3,312 PPIs from human nuclear XL-peptides, 1580 proteins; rows; violet) and plant data (generated by 1,211 Arabidopsis ortholog PPIs from plant nuclear XL-peptides, 877 Arabidopsis ortholog proteins; columns; orange). The similarity between two modules is evaluated by Jaccard Coefficient (JC), which is determined by the number of identical components divided by the total number of unique components. The green color of the heatmap stands for the level of Jaccard Coefficient. The violet and orange dots stand for the human-specific module variants (JC <0.6), plant-specific module variants (JC<0.6), and common modules (JC>0.6), respectively (Table S10e). C1, C2, C3 and C4 represents the community 1, 2, 3, and 4, respectively. Ungrouped are those modules that fail to be integrated into a community.

(C) The heatmap represents the community comparison between the human (rows; violet letters) and plant nuclear (columns; orange letters) communities. The similarity between two modules is evaluated by Jaccard Coefficient (JC), which is determined by the number of identical components divided by the total number of unique components. The green color of the heatmap stands for the level of Jaccard Coefficient. C1, C2, C3 and C4 represents the community 1, 2, 3 and 4, respectively. The data are in Table S10c-d.

(D) The heatmap shows the comparison of biological process (left panel), molecular function (middle panel), and cellular component (right panel) analysis of proteins derived from the human (rows; violet letters) and plant nuclear (columns; orange letters) communities. The top 10% of most interactive proteins (highest protein degree) in each community were used for the analysis. The similarity among the communities is evaluated by the Jaccard Coefficient (JC) of related GO terms. The green color of the heatmap stands for the level of the Jaccard Coefficient (Table S10f-g).
